# Neural Processes with Normalizing Flows for Wheat Height Estimation

**DOI:** 10.64898/2026.06.24.734247

**Authors:** Mike Boss, Michele Volpi, Lukas Roth

## Abstract

In this work, we investigate modeling plant traits over time using neural processes, a class of machine learning models that learn distributions over functions. Plant growth is an inherently stochastic process with complex dynamics measured mostly at irregular times throughout the growing seasons. While individual trait trajectories may be simple, their distributions are shaped by complex interactions between genotype, environment, and other factors.

In particular, we focus on plant height in wheat, a deceptively simple-looking trait with complex dynamics. To model these trajectory distributions, we evaluate neural processes and in particular extensions using normalizing flows, with different combinations of genotype and environmental covariates. For controlled evaluations, we generate synthetic wheat height trajectories calibrated against Swiss weather station records and the Field Phenotyping Platform 1.0 (*FIP1*) wheat dataset. To fully evaluate these trajectory distributions, we use signatures, vector representations of sequential data, together with the signature maximum mean discrepancy (*Sig-MMD*) and its recently introduced censored variant (*CSig-MMD*). *Sig-MMD* enables direct pathwise comparison of predicted and simulator trajectory distributions, while *CSig-MMD* focuses this comparison on the tail, including lodged trajectories. Together, these metrics allow us to assess whether the models capture the full distribution of growth trajectories, including rare outcomes. We compare the neural processes with a lodging-mixture spline baseline and apply them to the *FIP1* data.

## 1 Introduction

Plant trait measurements are inherently stochastic due to the nature of the growth process and the noisy measuring methods. In addition, measurements are often only taken at non-uniform, sparse intervals throughout the season [52]. For phenotyping, researchers are interested in the true underlying development rather than superficial variations that obscure the genuine differences between genotypes. However, even seemingly simple traits, such as plant height in wheat, have complex dynamics. For example, lodging events, i.e., the bending of the plants towards the end of the season, occur suddenly and are hard to predict as they are primarily caused by the height of the plants and random weather events [5].

Such events introduce uncertainty even under identical conditions. Moreover, growth depends on conditions that are only partly known, such as soil and management. The growth of a plot is one of many outcomes that could have occurred under the same known conditions. Plant growth can therefore be viewed as a stochastic process whose realizations are the growth curves. While traits of interest may be derived from the measured growth process, ideally models encode the underlying growth dynamics in a representation learned directly from the data [42, 45, 47]. Such representations allow for comparison of genotypes based on their growth characteristics rather than pre-selected traits [44]. Because growth is stochastic, the learned representation must capture this uncertainty to fully reflect the growth dynamics [46]. A deterministic representation models only the most likely outcome, and so would miss, for example, the lodging probability that distinguishes genotypes. Capturing such genotype differences is part of modeling the genotype-environment interaction that underlies prediction for new, unobserved environments, a long-standing goal in plant breeding [33].

Since trait measurements are always only a partial observation of an underlying stochastic growth process, neural processes (*NP*s) provide a natural framework for modeling plant growth with neural networks probabilistically [13]. In *NP*s, a subset of the data, referred to as the *context set C*, is used to predict the *target set T*. Similar to variational auto-encoders (*VAE*s) [21], *NP*s introduce a latent representation *z*, samples of which are then used to make stochastic predictions. In practice, one trained model then predicts the trajectory distribution of a plot from its genotype and environment before any height is measured, and sharpens this prediction as measurements arrive during the season, without refitting. This allows genotypes to be screened from their covariates under recorded weather, or the response of one genotype to be compared across temperature records, with the lodging probability as part of the prediction.

In standard *NP*s, both the latent distributions and the predictive likelihood are typically modeled as Gaussian distributions [13, 25]. This diagonal Gaussian assumption may limit the expressiveness of the model [27]. A diagonal Gaussian cannot capture non-Gaussian, multi-modal dynamics such as potential lodging, nor the correlations between multiple traits [43]. Normalizing flows (*NF*s) offer a way to increase the expressiveness of simple base distributions by transforming them into expressive target distributions with correlated dimensions. Such *NF*s can be applied to the Gaussian latent distributions used by *NP*s as seen in [27].

In this work, we analyze the use of *NP*s to model plant traits over time, specifically wheat height measurements. To investigate how well these models capture the underlying dynamics of plant growth, we introduce a fully synthetic wheat height generation model. We compare different *NP* variants, including different extensions with *NF*s. A lodging-mixture spline model with genotype and environment effects serves as the baseline, and the models are also trained and evaluated on the Field Phenotyping Platform 1.0 (*FIP1*) data. All variants share the same transformer backbone, shown in Figure 1.

**Figure 1:**
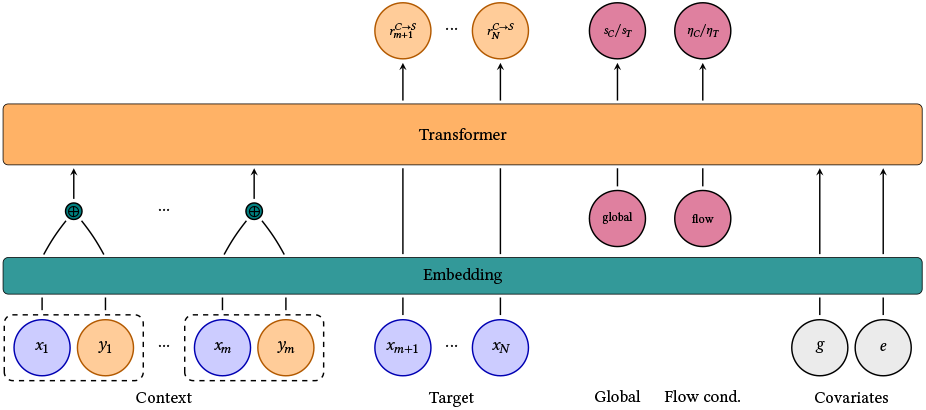
Shared transformer encoder, run in two passes. The *context pass* receives the context observations and query tokens at the prediction locations, producing the context summary *s*_*C*_, the flow-conditioning representation η_*C*_ for flow variants, and the per-target representations 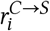 of the queried points *S*. The *target pass*, used only in training, receives the target observations and produces only the target summary *s*_*T*_ and η_*T*_. The covariates are the genotype-parameter vector *g* and the temperature trajectory *e*.

To compare these models, we assess whether they reproduce the distribution of full growth trajectories rather than only per-time-point marginals. Pointwise or marginal metrics can miss trajectory-level stochastic events such as lodging. Therefore, we compare the predicted and simulator trajectory distributions directly with signature maximum mean discrepancy (*Sig-MMD*) [8], a divergence based on the signature kernel. Signatures can capture the geometry of trajectories and correlations between multiple traits, allowing trajectories to be compared pathwise rather than as collections of individual points. We further incorporate censored signature maximum mean discrepancy (*CSig-MMD*) [34] for better tail-event evaluation and outlier detection, such as lodging events. *CSig-MMD* is especially relevant here, since breeders are typically interested in the extreme phenotypes [3, 5], which it evaluates from the full trajectory shapes. The synthetic generator, the models, and the evaluation code are available at http://hdl.handle.net/20.500.11850/806730.

## 2 Preliminaries

The preliminaries are introduced in the context of individual plant trait estimation, where for each experimental unit there is a set of pairs {(*x*_*i*_, *y*_*i*_)}_*i*∈ℐ_. Here, *x*_*i*_ are the measurement time points, *y*_*i*_ the corresponding measurements, and ℐ the index set for the specific experimental unit. Experiment data consists of a collection *D* of such measurement sets, one for each experimental unit. In the context of wheat height estimation, the unit is a plot of a single genotype grown over one season, and *x*_*i*_ and *y*_*i*_ are days and heights, respectively.

Each growth curve is one realization of the growth process, and the *NP* models learn the distribution over such realizations. The *NP*s are extended with *NF*s to increase their expressiveness. The models are evaluated with two signature-based metrics, *Sig-MMD* and its tail-focused variant *CSig-MMD*, which build on the components introduced below.

### 2.1 Neural Processes

A stochastic process can be defined as *X*: *J* × Ω → *S*, where *J* is an index set, *S* is the state space, and Ω is the sample space of a suitable probability space [9]. Given any ω ∈ Ω, *t* ↦ *X* (*t*, ω) is a function that is a realization of the process over the domain. Plant growth can be described with *J* as the set of all time points during the season, *S* as the set of all possible heights, and the realizations as possible growth curves.

*NP*s approximate stochastic processes by modeling a predictive conditional distribution *p*_θ_(*y*_*T*_ ∣ *x*_*T*_, *x*_*C*_, *y*_*C*_), where θ denotes the parameters of a neural network. Each data index set ℐ is split into the *context set C* and the *target set T*, for which commonly *C* ⊆ *T* = ℐ. The predictive distribution is modeled using a latent variable model with a global latent variable *z* as:

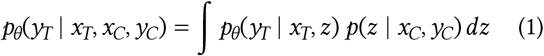

where *p*(*z* ∣ *x*_*C*_, *y*_*C*_) is the true conditional prior distribution over the latent variable, and *p*_θ_(*y*_*T*_ ∣ *x*_*T*_, *z*) is the conditional likelihood of the predictions.

The marginal predictive distribution requires integrating over the latent variable *z*, which is generally intractable because the true context-conditioned prior is unknown and the neural-network parameterization of the conditional likelihood *p*_θ_(*y*_*T*_ ∣ *x*_*T*_, *z*) is nonlinear [13]. Therefore, *NP*s use a shared encoder network with parameters ϕ to parameterize a context-conditioned prior *q*_ϕ_(*z* ∣ *x*_*C*_, *y*_*C*_) and a target-conditioned variational posterior *q*_ϕ_(*z* ∣ *x*_*T*_, *y*_*T*_). The model is trained by variational inference, maximizing the evidence lower bound (*ELBO*) with respect to both θ and ϕ [13, 25]:

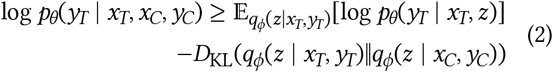

where *D*_KL_ is the Kullback–Leibler divergence (*KL*).

### 2.2 Normalizing Flows

*NP*s assume a Gaussian latent distribution, which can be a limitation. *NF*s can increase the expressiveness of these distributions by transforming a simple *base* distribution *p*_0_(*z*_0_), usually a standard Gaussian, into an expressive *target* distribution through an invertible transformation *f*_*c*_. Both the base distribution and the transformation may be conditioned on auxiliary variables *c* [51]. The invertibility enables both sampling and exact density evaluation through the change-of-variables formula. For *z* = *f*_*c*_(*z*_0_),

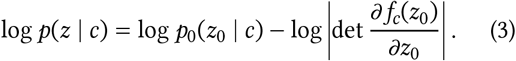

Equivalently, for a given *z*, the density can be evaluated by mapping 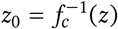.

Some architectures exhibit a computational asymmetry, being cheaper to evaluate in one direction than in the other. Autoregressive flows, for instance, evaluate one direction in a single pass but require a separate sequential pass per latent dimension in the other [31].

### 2.3 Signatures

Each realization of the growth process can be described as a continuous path *y* : [*a, b*] → ℝ^*d*^ mapping time *t* to trait values *y*_*t*_. Its shape can be described by the *signature* [7], a hierarchical summary of the path built from iterated integrals and organized into levels of increasing order.

Each level captures progressively finer detail about the path’s shape. The *n*-th level contains all *d*^*n*^ integrals of order *n*:

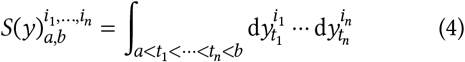

where *t*_1_ < … < *t*_*n*_ are ordered values of *t*, and 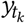 denotes the path evaluated at time *t*_*k*_. The signature captures a path’s shape but not its speed, making it invariant to reparametrization. For a one-dimensional path, it reduces to the net change alone. These limitations can be addressed by *augmenting* the path with additional dimensions [8]. Adding a time channel makes the signature sensitive to the timing of changes, while the lead-lag transform increases its sensitivity to the path’s geometric variation.

The full signature is infinite-dimensional. *Truncated signatures* retain only the first *N* levels and can be used directly as features, while the *signature kernel* [39, 26] uses the kernel trick to implicitly compute the inner product of infinite-dimensional signatures [8].

### 2.4 Maximum Mean Discrepancy

Comparing the predicted and simulator trajectory distributions requires a distance between distributions. The maximum mean discrepancy, *MMD* [17], measures the discrepancy between two distributions **P** and **Q** using a positive semi-definite kernel *k*:

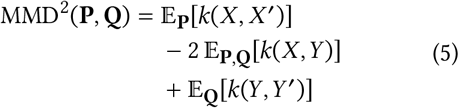

where *X, X* ^′^ ∼ **P** and *Y, Y* ^′^ ∼ **Q** are independent samples.

### 2.5 Mahalanobis Distance

Identifying rare, extreme trajectories requires measuring how far each one lies from a reference distribution. The Mahalanobis distance provides such a measure, accounting for the spread and correlations of that distribution. Let *x* ∈ ℝ^*d*^ be a feature vector, and let µ and Σ denote the mean vector and covariance matrix of a reference distribution in feature space. The Mahalanobis distance [28] of *x* from this reference distribution is defined as

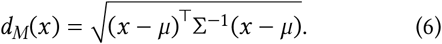

### 2.6 Kernel Normalization

Normalizing the signature kernel places every path on a common scale, so its overall size no longer dominates the comparison of shape. For a positive semi-definite kernel *k* with *k*(*X, X*), *k*(*Y, Y*) > 0, the normalized kernel is defined by [16]

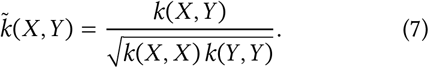

In particular, 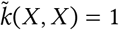.

### 2.7 Importance Sampling

A model that only produces whole trajectories cannot be conditioned on observed points directly. Importance sampling conditions it by weighting each draw with the likelihood of the observations under that draw, so that resampling the draws with these weights yields draws from the posterior given the observations [14]. For a pool *p* of candidate trajectories *H*_*c*_ and heights *H*_*i*_ observed at *x*_*i*_, *i* ∈ *C*, with independent Gaussian noise of scale σ_obs_, the weights are

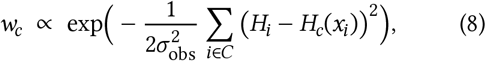

normalized over the candidates. The effective number of draws, 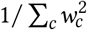, falls as the observations constrain the posterior, so the pool must be large enough to contain draws that explain them.

## 3 Data

We are interested in evaluating how well models capture the underlying dynamics of plant growth. Unfortunately, real-world high-throughput phenotyping data is sparse, noisy, and not available in large enough quantities for controlled distributional comparisons [3]. In addition, even simple real-world dynamics are influenced by many factors [49, 37]. For example, lodging depends on genotype, environmental conditions, and specific weather events.

To address these issues, synthetic wheat height trajectories can be generated, calibrated against real data. The generator has two parts, a temperature generation model and a height generation model, and is intentionally simpler than full weather or crop growth models [32, 18]. The goal is not to reproduce wheat development in full detail, but to create controlled trajectory distributions whose sources of variation are known and can be sampled repeatedly. Within this synthetic setting, genotypic information is represented by the simulator’s parameter vectors. They serve as simplified substitutes for real genotype representations, rather than observed marker data. Environmental covariates are synthetic full-season temperature trajectories.

*P*_sim_ denotes distributions defined by this synthetic simulator. A condition is *u* = (*g, e*), where *g* is the simulator-defined genotype-parameter vector and *e* is the generated temperature trajectory. For a fixed condition *u*, the non-lodged growth curve is deterministic, and *P*_sim_(**H** ∣ *u*) denotes the distribution of underlying height trajectories **H** obtained by resampling lodging. Noisy observations are generated from these trajectories by adding independent observation noise. Marginal distributions such as *P*_sim_(**H**), *P*_sim_(**H** ∣ *g*), and *P*_sim_(**H** ∣ *e*) are obtained by marginalizing over the unspecified covariates.

### 3.1 Temperature Generation

The synthetic temperature generator produces hourly temperature trajectories for arbitrary year-site combinations. The model additively decomposes hourly temperature at site *s*, year *y*, day *d*, and hour *h* as:

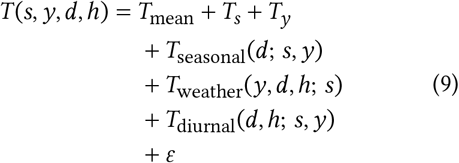

where *T*_mean_ is the global mean, 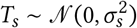 a site offset, 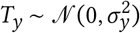 a year anomaly, and 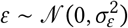 measurement noise.

The seasonal component *T*_seasonal_ follows a two-harmonic Fourier model whose amplitude and phase vary by site and year, capturing regional differences and inter-annual variability. Weather variability *T*_weather_ is modeled as an hourly first-order autoregressive (*AR(1)*) process with both shared and site-specific variation. The diurnal component *T*_diurnal_ produces an asymmetric daily cycle whose amplitude and shape vary seasonally through a shared seasonal factor based on a two-harmonic fit of the diurnal ranges. Further details are given in Appendix G.

### 3.2 Height Generation Model

The height generation model takes temperature as input [36] and consists of two components: a growth rate surface and a developmental clock.

Development is tracked by thermal time [50, 23], the cumulative sum of the positive part of hourly temperatures *T*_*t*_, where *t* indexes the hours of the season consecutively:

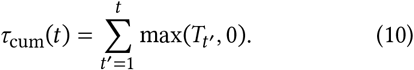

Normalizing by a thermal time budget τ_max_ and clipping at maturity gives a developmental index

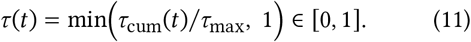

Growth ceases at maturity, where τ = 1 and *R*(*T*, 1) = 0. The index τ is the fraction of the thermal-time budget used since a fixed start day. It is a quantity of the simulator, not a phenological scale.

The growth rate is modeled as a tensor product of cubic B-splines over temperature and developmental stage:

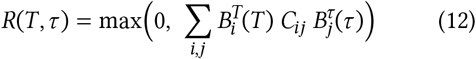

where 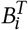 and 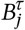 are cubic B-spline basis functions over the temperature and developmental-stage axes, and the control point matrix **C**, with entries *C*_*ij*_, weights their tensor products to define the shape of the response surface. Since τ depends on temperature through thermal time, warmer conditions both increase the instantaneous growth rate and advance development, causing the surface to be traversed faster. The growth model first produces a lodging-free height trajectory *H* (*d*) by summing hourly growth rates:

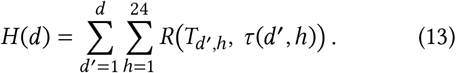

To simulate diverse synthetic genotypes, the interior control points of **C** are sampled around a calibrated mean response surface. Figure 2 shows one such sampled surface. The correlated prior yields smooth genotype-specific variations (Figure 3). Boundary control points are fixed at zero, enforcing no growth at the temperature limits and at the beginning and end of development. The outer temperature knots set a minimum and a maximum temperature (0 and 35 ^°^C) at and beyond which the growth rate is zero. They are chosen to lie roughly within the range reported for wheat [50, 36] and restrict the calibration to biologically plausible growth responses. The thermal-time budget τ_max_ is sampled from a calibrated truncated-normal distribution.

**Figure 2:**
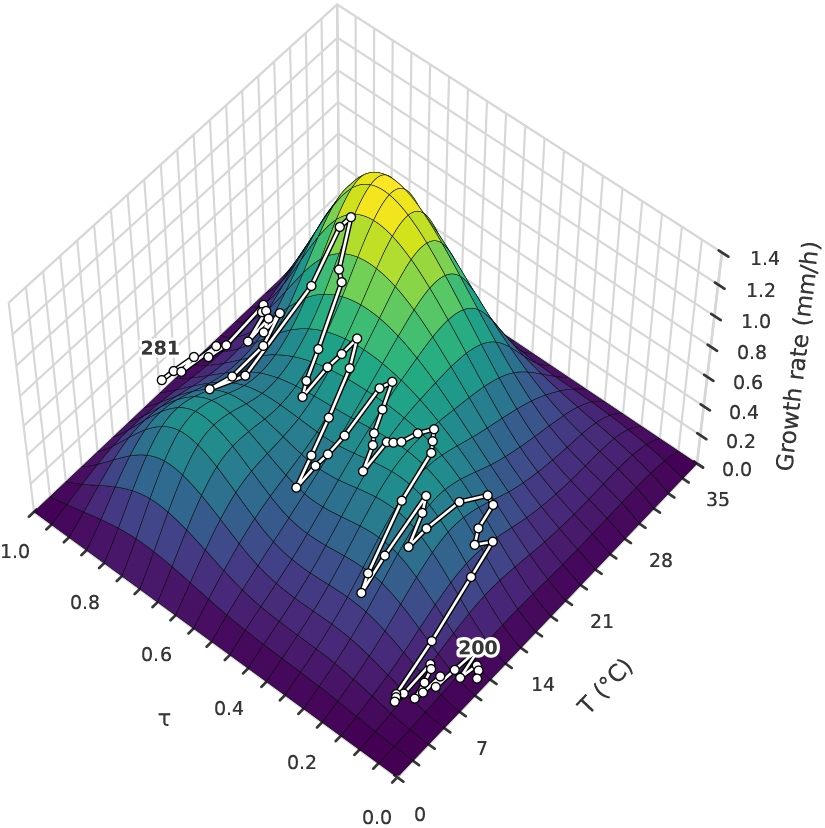
Growth rate surface *R*(*T*, τ) of one synthetic genotype, sampled from the calibrated generator, as a tensor product of cubic B-splines over temperature *T* and developmental index τ, the fraction of the simulator’s thermal-time budget used since a fixed start day. The surface defines the instantaneous growth rate, which is summed over hourly temperatures to obtain plant height (eqs. (12) and (13)). The white path shows one synthetic season on this surface, with one dot per day at the daily mean temperature and the developmental index reached by the end of that day. Its labels give the start day and the day of maturity, counted from the first of September.

**Figure 3:**
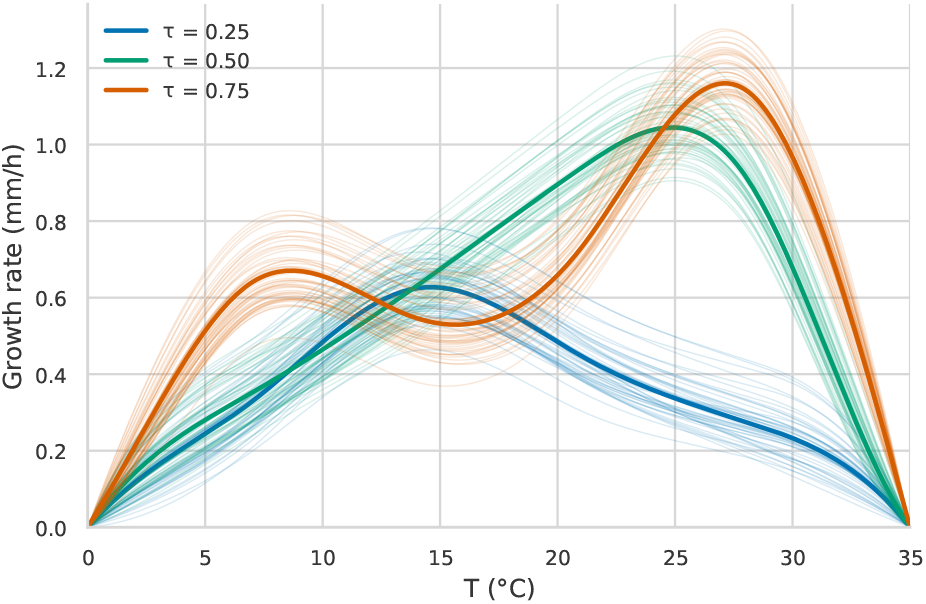
Variation in genotype-specific growth-rate curves. Thin lines show growth-rate curves sampled from the synthetic generator as functions of temperature *T* at three equally spaced values τ ∈ {0.25, 0.5, 0.75} of the developmental index, the fraction of the thermal-time budget used since a fixed start day. Thick lines show the corresponding means. In simulations with the calibrated model, plants have reached about 17 %, 52 %, and 87 % of their final height at these values, which places them at the start, in the middle, and near the end of stem elongation.

Lodging is modeled as a stochastic event whose probability increases with maximum height and which can occur only after maximum height has been reached. After generating the clean growth curve *H* (*d*), whether lodging occurs is sampled using a probability based on *h*_max_ = max_*d*_ *H* (*d*) (Appendix H). If it occurs, the trajectory drops over a short random transition to a random fraction of *h*_max_. Thus, for a fixed genotype and environment, resampling lodging produces the remaining trajectory variability.

Observed synthetic heights are obtained by adding independent Gaussian observation noise to the sampled trajectories. Numerical generator parameters are listed in Appendix A.

### 3.3 Synthetic Sample Construction

Synthetic samples are generated by first sampling a yearsite-specific temperature trajectory and genotype-specific height-model parameters, then generating the height trajectory from both. For model inputs, genotype information is provided as the normalized genotype-parameter vector and environmental information as the corresponding hourly temperature trajectory.

### 3.4 Calibration

Both generation models are calibrated against real-world data: the temperature model against Swiss weather station records and the height model against the *FIP1* dataset [38]. The height calibration used 3,806 *FIP1* plots across five harvest years (2016, 2017, 2018, 2021, and 2022). The temperature calibration used 151 complete station-year records from eight winter wheat variety testing sites over harvest years from 2004 to 2024 (Appendix G.1).

For the height model, genotype-specific control point matrices are drawn from a Gaussian prior centered on a shared mean surface, with smooth spatial correlations along the temperature and developmental-stage dimensions. The calibrated quantities are the mean surface, the prior amplitude and kernel length scales, and the distribution of τ_max_. These are jointly optimized using Optuna [1], where the objective compares simulated and observed distributions using the *MMD* (Section 2.4) on height trajectories, maximum heights, and growth cessation times. The objective is the unweighted sum of these three distributional terms, with additional guardrail penalties for finite trajectories, plausible mean height, and the expected tendency for warmer seasons to reach maturity earlier and therefore produce shorter plants [50, 24]. The lodging parameters are fixed rather than calibrated, so that the lodging rate can be set and varied across datasets, and *FIP1* contains too few lodging events to constrain them.

To quantify how well the calibrated generator matches *FIP1*, simulated and observed trajectories are compared per year with the *Sig-MMD* of Section 2.4. The scores are read against the discrepancy between genotype groups of the same year and the discrepancy between years, both computed from the observed plots (Appendix A.1). The height distributions were fitted on 2017, 2018, 2021, and 2022, and the 2016 plots entered only the growth-cessation term of the objective. In these four years, the simulator lies between the two references.

The calibration matches the plot population of each year without using genotype identity, so it does not constrain how the fitted spread divides between genotypes and between plots of the same genotype. Appendix A.2 therefore compares simulated and observed genotypes in the spread of three traits, maximum height, growth-cessation day, and elongation rate, and in the stability of genotype rankings across years. The simulator is close to *FIP1* in the spread of maximum height and in rank stability, whereas its spread of growth-cessation day is too wide and its spread of elongation rate too narrow in most years.

## 4 Model Architectures

All *NP* variants share a transformer encoder-decoder backbone [48]. They are transformer-based versions of the standard neural-process models: conditional neural process (*CNP*) [12], attentive conditional neural process (*ACNP*), latent neural process (*LNP*), and attentive neural process (*ANP*) [20]. The variants differ along two axes. The first axis is *conditional* versus *latent*. Conditional models use a context summary directly, while latent models sample a global latent variable *z*. The second axis is *global* versus *per-target*. Global models decode each target using only its location embedding, while per-target models also encode the target locations as query tokens to produce per-target representations. The latent variants are further extended by applying *NF*s to their Gaussian latent distributions on the prior, the posterior, or both.

### 4.1 Encoder-decoder backbone

The encoder, depicted in Figure 1, is similar to those of other attention-based *NP*s [20, 29, 30]. It embeds observation pairs {(*x*_*i*_, *y*_*i*_)}, a learnable global token, and, for the flow variants, a learnable flow-condition token. When supplied, the genotype-parameter vector *g* and temperature trajectory *e* are embedded as covariate tokens. Each observation token is the sum of its *x*_*i*_ and *y*_*i*_ embeddings. The global token yields a summary representation of the encoded set. Encoding the context set *C* gives *s*_*C*_, and, during training, encoding the target set *T* gives *s*_*T*_. In the flow variants, the flow-condition token likewise yields η_*C*_ and η_*T*_.

Per-target models additionally encode each prediction location as a query token, which the transformer maps to a per-target representation 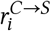, where *C* → *S* indicates that the query attends to the observations of the context set *C* and is computed for the queried points in the scoring set *S* (Section 5). The attention masks are given in Appendix D.

The decoder, depicted in Figure 4, predicts each 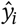 from two inputs. One is a set-level summary shared across all predictions, either the context representation *s*_*C*_ for conditional models or a latent sample *z* for latent models. The other is the per-target input *t*_*i*_, which is 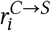 in pertarget models or the embedded prediction location *x*_*i*_ in global models. Both depend only on information available at inference.

**Figure 4:**
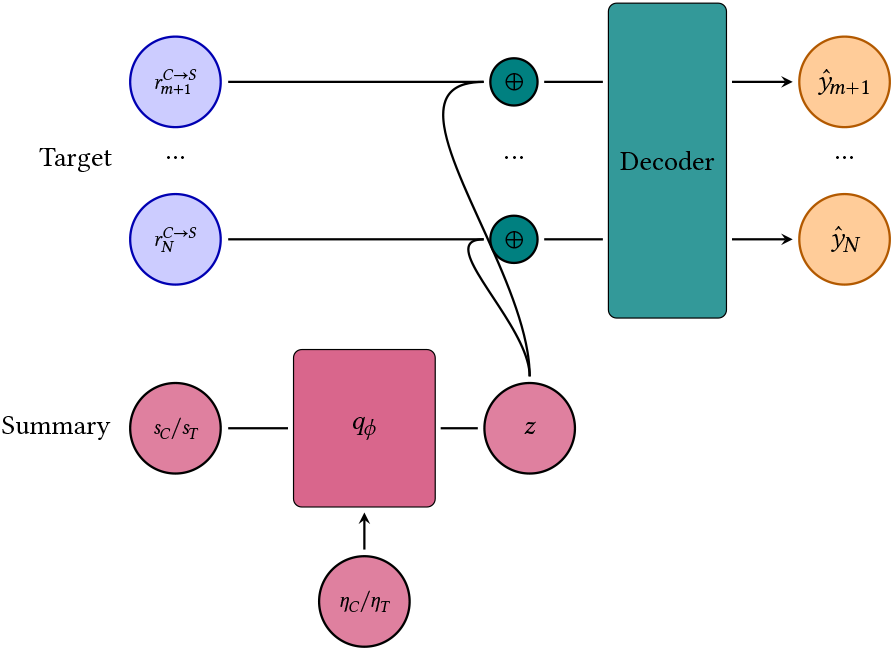
Decoder structure. Conditional models decode from the context summary *s*_*C*_. Latent models decode from a sample *z* of the latent distribution *q*_ϕ_, parameterized by *s*_*C*_ for the prior and, in training, by the target summary *s*_*T*_ for the posterior. In flow variants, the flow-conditioning representations η_*C*_ and η_*T*_ from the encoder condition the prior and posterior flows. The per-target input is the prediction-location embedding for global models and 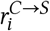 from the context pass for per-target models.

### 4.2 Base variants

The variants differ in how they use these representations (Table 1). The conditional variants, *CNP* and *ACNP*, pass *s*_*C*_ to the decoder directly and have no latent variable sampling. The latent variants, *LNP* and *ANP*, instead use the summary representations to parameterize diagonal Gaussians over a global latent *z*. Here *s*_*C*_ parameterizes the prior *q*_ϕ_(*z* ∣ *x*_*C*_, *y*_*C*_), and, during training, *s*_*T*_ parameterizes the posterior *q*_ϕ_(*z* ∣ *x*_*T*_, *y*_*T*_). The normalizing-flow variants of Section 4.3 transform these latent distributions.

**Table 1:**
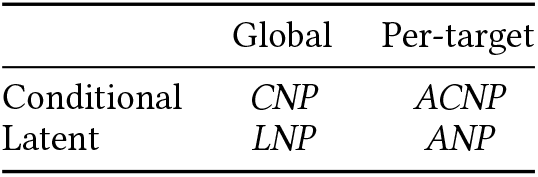
The four base variants as transformer-based neural-process models. Rows indicate whether the decoder uses the context summary directly (conditional) or samples a global latent variable *z* (latent). Columns indicate whether prediction locations are represented only by their location embeddings (global) or encoded as query tokens that produce per-target representations 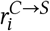 (pertarget). *CNP, ACNP, LNP*, and *ANP* abbreviate the conditional, attentive conditional, latent, and attentive neural process.

|  | Global | Per-target |
| --- | --- | --- |
| Conditional | *CNP* | *ACNP* |
| Latent | *LNP* | *ANP* |

A conditional variant predicts the targets independently,

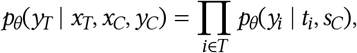

while a latent variant marginalizes the same factorized likelihood over the prior,

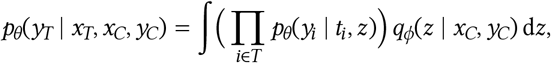

recovering the latent-variable model of eq. (1).

### 4.3 Normalizing-flow latent distributions

Increasing the expressiveness of the prior is uncommon in latent-variable models such as *VAE*s, but can be beneficial [2, 4, 19]. It may be particularly beneficial in *NP*s, where the prior and posterior latent distributions differ only in their conditioning sets. Since the prior is conditioned on less information than the posterior, it may need to model a potentially more complex and multi-modal distribution. For instance, when the context holds only pre-lodged observations, the prior must cover both outcomes, whereas the posterior, which also potentially sees the lodged points, could be simpler. The prior is also the only distribution used at inference, since predictions sample *z* from *q*_ϕ_(*z* ∣ *x*_*C*_, *y*_*C*_), while the posterior *q*_ϕ_(*z* ∣ *x*_*T*_, *y*_*T*_) is only used during training. Its expressiveness therefore directly bounds the predictive distribution.

The Gaussian assumption is therefore relaxed by transforming the latent distributions with *NF*s. Flow variants are defined only for latent models, since the conditional variants have no latent distribution. For the prior, *s*_*C*_ parameterizes the Gaussian base distribution and η_*C*_ conditions the flow. For the posterior, *s*_*T*_ and η_*T*_ play the corresponding roles. A conditional flow *f*_*c*_ maps a Gaussian base sample *z*_0_ ∼ *q*_0,ϕ_(*z*_0_ ∣ *s*) to the latent *z* = *f*_*c*_(*z*_0_), with density given by the change of variables of Section 2.2. Following [22], the flow is partially amortized through the flow-conditioning representations η_*C*_ and η_*T*_, rather than fully amortized by predicting its parameters per sample [35].

The flow can wrap the prior, the posterior, or both. The resulting model names follow the pattern

{LNP, ANP}-NF-{Prior, Posterior, Prior-Posterior}.

Here *Prior-Posterior* denotes separate flows on both the prior and posterior. The placement interacts with the asymmetric cost of evaluating a flow forward versus in reverse (Section 2.2). The posterior is only ever sampled, and the *KL* term needs its density solely at those samples, which the forward pass already produces, so its inverse is never evaluated. The prior, by contrast, is used in both directions. It is sampled forward to draw predictions at inference, and evaluated in reverse to score posterior samples for the *KL* during training. A posterior flow therefore must be efficient in only one direction, whereas the prior flow must be efficient in both.

## 5 Training

In standard *NP* training [25], each sampled trajectory is split into a context set *C* and a target set *T* with *C* ⊆ *T*. The prior is conditioned on the context set *C*, the posterior on the target set *T*, and the reconstruction loss is evaluated on the target set *T*.

Since *C* ⊆ *T*, the loss scores observations that are also provided as context. This can cause *in-context overfitting* [40], where the model reproduces noisy context points instead of the underlying noise-free trajectory. To mitigate this, [40] scores only the non-context targets *T* \*C*. The posterior could similarly overfit to its conditioning set *T*, which is also scored. This is less likely, as the target values influence the loss only through the latent path. The held-out constructions add points *O* outside *T*, which the posterior does not condition on.

These choices are studied by fixing the roles of the context set *C* and target set *T*, with *C* ⊆ *T*, and varying only the *scoring set S* on which the likelihood is scored. For the latent variants the training objective is

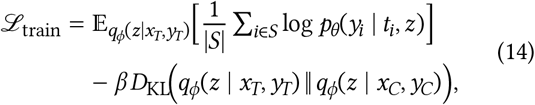

Here *t*_*i*_ is the per-target input defined above. The likelihood term is averaged over scored points. The coefficient β weights the *KL* term, and standard *NP* training is recovered by *S* = *T*. For each training trajectory, the encoder is run twice. The context pass conditions on *C* to produce the prior and the per-target inputs *t*_*i*_, while the target pass conditions on *T* only to produce the posterior *q*_ϕ_(*z* ∣ *x*_*T*_, *y*_*T*_) and, for flow variants, its flow-conditioning. Target values therefore reach the likelihood only through the latent sample *z* and never through *t*_*i*_. For the conditional variants the *KL* term vanishes and the objective reduces to the predictive log-likelihood on *S*. The flow variants use the same objective, with *NF*s applied to the latent distributions.

Four scoring sets follow from the two concerns above (Table 2). Standard *NP* training scores the full target set, *S* = *T*, and excluding the *context* gives *S* = *T* \ *C*. The analogous pair for the posterior adds a held-out set *O* with *O* ∩ *T* = ∅ and scores *S* = *T* ⋃ *O* or, excluding the *target*, only *S* = *O*.

**Table 2:** The four scoring-set constructions along two axes. The first row includes the relevant conditioning set in the loss. The second excludes it. The columns indicate whether a held-out set *O* (*O* ∩ *T* = ∅) is present, which shifts that conditioning set from the context *C* (prior) to the target *T* (posterior). The resulting scoring sets are *S* = *T, S* = *T* \ *C, S* = *T* ⋃ *O*, and *S* = *O*. The prior is conditioned on the context set *C* and the posterior on the target set *T*, with *C* ⊆ *T*.

| | only $T$ | with $O$ |
| --- | --- | --- |
| conditioning scored | $S = T$ | $S = T \cup O$ |
| conditioning excluded | $S = T \setminus C$ | $S = O$ |

## 6 Evaluation

Standard pointwise metrics and marginal scores, such as root mean squared error (*RMSE*) or per-time-step continuous ranked probability score (*CRPS*), compare predictions and observations at individual time points [15]. Rare, high-impact events such as lodging can distort these metrics. A well-calibrated model can incur a large error whenever the single observed outcome happens to be lodged. Trajectory dependencies can worsen this and may even reward poorer calibration. Because simulated lodging depends on the maximum height reached, a well-calibrated model predicts more or less lodging according to its predicted height. This is desirable, yet distorts the metrics, since a non-lodged outcome penalizes models that overpredict height while a lodged outcome penalizes those that underpredict height.

To address these issues, *trajectory distributions* have to be compared directly. Rather than comparing to a single outcome, a divergence between the predicted and simulator distributions is used. A natural candidate is the *MMD* (Section 2.4), which measures the discrepancy between two distributions through a kernel. Crucially, the kernel determines what aspects of the data the *MMD* is sensitive to. A pointwise comparison treats each time step independently. The signature kernel (Section 2.3) is used instead, comparing paths through their signatures and thus capturing trajectory geometry. Since the trajectories are one-dimensional, the signature would reduce to the net change alone. Time augmentation and the lead-lag transform are therefore applied to retain sensitivity to the full trajectory shape. The kernel is normalized (Section 2.6) to correct for the bias toward trajectories with larger height values, which produce larger increments and thus larger self-norms. The signature kernels, *Sig-MMD*, and truncated signatures are computed using pySigLib [41].

### 6.1 Simulator distributions

Distributional scores compare model samples to a simulator distribution. A single *matched* simulator distribution is used, defined so that each prediction is scored against the distribution over the trajectories consistent with the information the model was given, both its covariates and any observed height context.

The simulator is a forward model whose only inputs are a condition *u* = (*g, e*). Both the genotype *g* and the environment *e* are themselves drawn from generative distributions. A covariate regime fixes the supplied covariates, and marginalizing over the unspecified ones gives the simulator distribution consistent with them. Observed height points are outputs of this process rather than inputs, so the simulator cannot be conditioned on them.

The covariates fix a candidate pool of conditions. Let *p* denote the set of conditions compatible with the supplied covariates. The *g*+*e* regime fixes the condition, so *p* = {*u*}. The *g* regime leaves the environment free, so *p* is all conditions sharing genotype *g*. The *e* regime analogously leaves the genotype free. The ∅ regime leaves both free, so *p* is the full set of conditions. With no observed context, the matched simulator distribution is the corresponding marginal of Table 3.

**Table 3:**
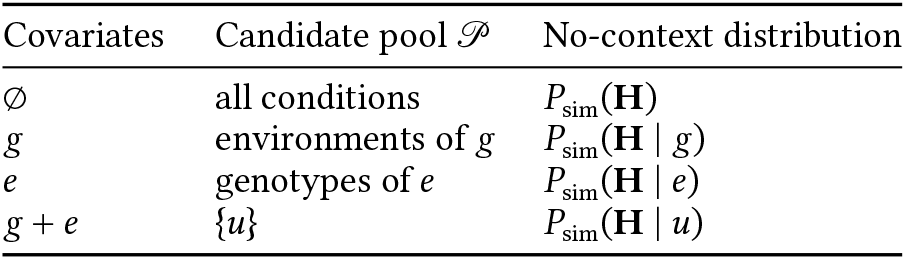
The *matched* simulator distribution by covariate regime. The candidate pool *p* is the set of conditions consistent with the supplied covariates, and the no-context distribution is the simulator marginal over that pool. Observed height context reweights the pool by eq. (8).

Observed height context narrows this pool. The context regimes reveal only pre-lodging height points, before lodging can occur, so the simulator distribution still leaves open the lodging variability to be predicted. For a fixed condition the pre-lodging growth curve is deterministic, so these context points are noisy measurements of that curve. The pool is therefore conditioned on them by importance sampling (Section 2.7), where the weight *w*_*c*_ of eq. (8) for a condition *c* uses its clean curve *H*_*c*_ and the observation-noise scale σ_obs_ of the generator. The matched simulator distribution draws a condition *c* ∝ *w*_*c*_ and then a lodging-resampled trajectory of *c*, so it samples the posterior-weighted mixture 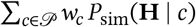. With no context the sum is empty and the weights are uniform, recovering the covariate marginal. When the covariates already fix the condition, the pool is a single element and the weights are irrelevant, recovering a comparison against that condition’s own lodging distribution *P*_sim_(**H** ∣ *u*).

This construction matches every predicted distribution to a simulator distribution built from the same information, shown in the Ground Truth column of Figure 5. As more context is observed, the weight concentrates on the conditions whose clean curves stay consistent with the observed points. Under partial covariates these are not always a single condition. The context never reveals where the curve peaks, so conditions that match the observed points but keep growing afterward stay equally consistent with them. A simulator distribution fixed to the lone true condition would instead penalize a model for the continued growth these observations leave open. Matched-reference scores compare model variants for a fixed setting, defined by the supplied covariates and observed context. Appendix I instead scores every prediction against its own condition, so that scores can be compared across covariate regimes.

**Figure 5:**
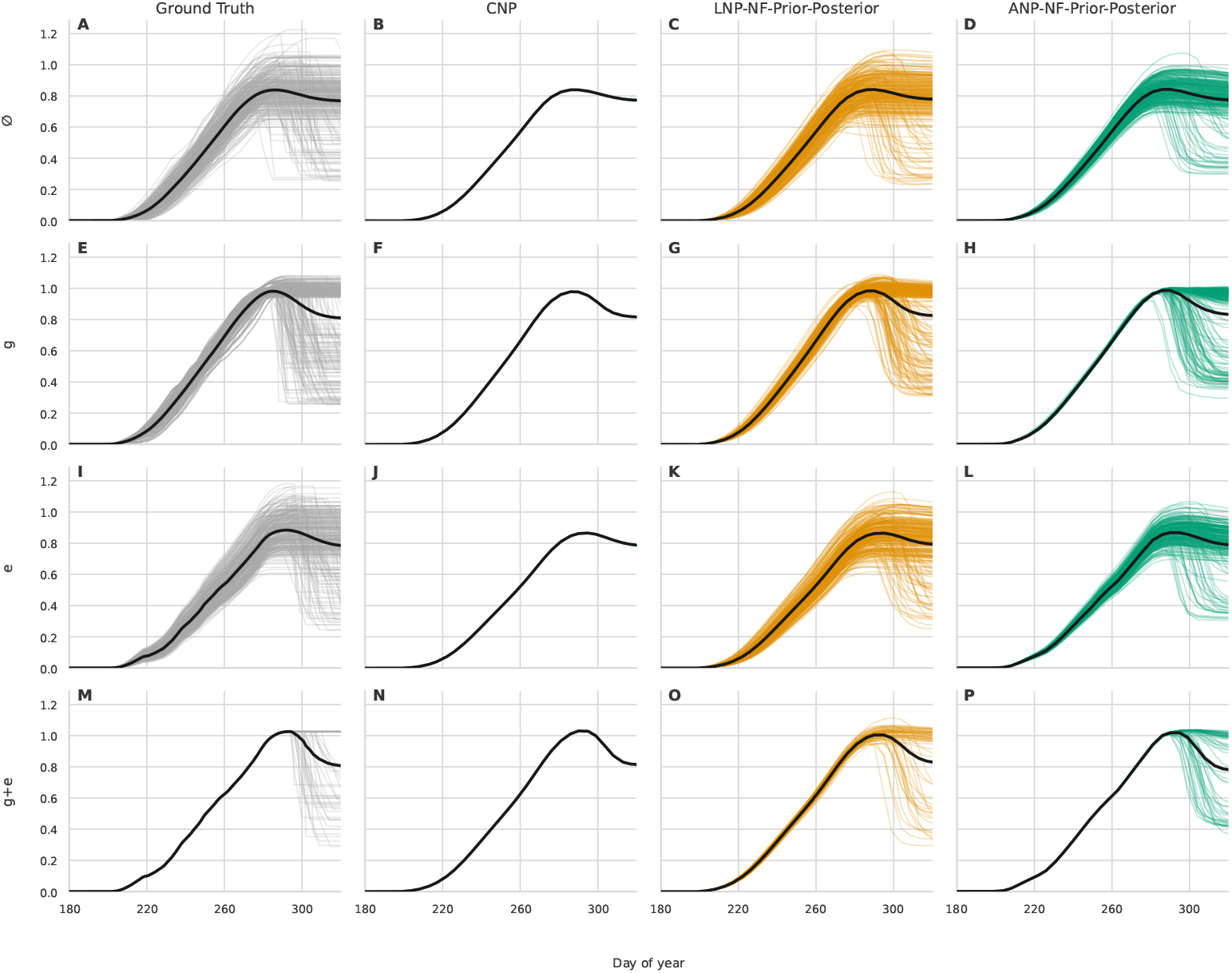
Height trajectory samples under each covariate regime. Columns show the *Ground Truth*, the conditional neural process (*CNP*), and the latent (*LNP*) and attentive (*ANP*) neural processes with normalizing flows (*NF*) on both prior and posterior (*LNP-NF-Prior-Posterior, ANP-NF-Prior-Posterior*). The *Ground Truth* column shows synthetic trajectories from the simulator that are consistent with the covariates of the row, and the other columns show model predictions. Rows show the covariate regimes with no covariates (∅), the genotype-parameter vector (*g*), the temperature trajectory (*e*), and both (*g* + *e*). The row label gives the covariates passed to the model. The *g, e*, and *g* + *e* rows use one genotype and one year-site, selected near the 90th percentile of maximum height. No height context is given. Thin lines are individual samples, and thick lines are their means. Days are counted from the first of September. Since the *CNP* has no latent variable, it produces a single predictive mean trajectory for each covariate regime.

### 6.2 Sig-MMD

Overall distributional fit is measured with *Sig-MMD* [8, 34], the *MMD* with the normalized signature kernel on time-augmented lead-lag paths. The signature kernel is evaluated in closed form, so the comparison uses the full, untruncated signature. *Sig-MMD* thus evaluates whether the model reproduces the chosen simulator trajectory distribution, capturing dependencies across time steps that pointwise metrics ignore.

### 6.3 CSig-MMD

*Sig-MMD* weighs the whole distribution, so the rare trajectories of greatest interest, such as lodging, contribute little to the overall score. *CSig-MMD* [34] is a censored variant of *Sig-MMD* that concentrates the divergence on this tail.

The tail is identified by how unusual a trajectory’s shape is. Each trajectory is summarized by its truncated signature, and its Mahalanobis distance (Section 2.5) to a robust minimum regularized covariance determinant (*MRCD*) [6] estimate fitted in this signature space measures how far into the tail it lies. The tail threshold is set on the simulator draws of the training conditions so that nearly all lodged trajectories lie beyond it.

Each trajectory is then softly censored toward a common pivot, a flat zero-height path, by an amount that grows with its distance below the tail threshold. Trajectories far in the tail are largely preserved, while typical non-tail trajectories are down-weighted toward the pivot. The truncated signatures are used only to compute tail distances and censoring weights. After censoring, the *MMD* itself is evaluated with the same normalized signature kernel as *Sig-MMD*. Thus the score is driven primarily by how well the model reproduces tail trajectories and their probability mass. The robust estimate, the tail threshold, and the steepness of the censoring boundary are given in Appendix E, and Appendix F reports how the model order depends on them and on the kernel bandwidth.

## 7 Experiments

### 7.1 Model architecture

All variants share the same transformer, described in Section 4, and differ only along the conditional/latent and global/per-target axes. The encoder uses 12 pre-norm transformer blocks with 128-dimensional hidden states and 4 attention heads, and the latent variants use a 32-dimensional global latent variable. The decoder predicts each height under a homoscedastic Gaussian likelihood with a single learned observation variance.

For the flow variants, a neural spline flow (*NSF*) [10] is applied to the Gaussian latent (Section 4.3). The prior flow uses coupling layers, which are efficient in both directions, and the forward-only posterior flow a more expressive autoregressive transform. Remaining dimensions, including the genotype-parameter vector and temperature encoders, are listed in Appendix D.

### 7.2 Baseline

As a baseline that, unlike the *CNP* and *ACNP*, can produce both lodged and non-lodged trajectories for a condition *u*, we fit a lodging-mixture spline model

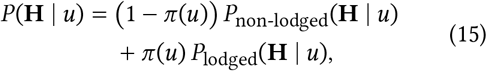

with a lodging probability π (*u*). Each growth curve is a cubic P-spline, a B-spline with a difference penalty on its coefficients [11], and these coefficients are the sum of a mean, a genotype effect, a year-site effect, and a Gaussian residual. Lodging is a single drop after the end of growth with a probability that rises with the peak height.

A neural process handles non-uniform sampling, unseen genotypes and year-sites, and context points through the same network, whereas the baseline requires an addition for each. The P-spline is fitted on the observed days of each trajectory and stays smooth where observations are missing. An unseen genotype or year-site has no fitted effect, so its effect is predicted by ridge regression on the genotype covariate or on the thermal time. The genotype covariate is the genotype-parameter vector on the synthetic data and the markers on *FIP1*. The baseline only draws whole trajectories, so it is conditioned on context points by importance sampling (Section 2.7), which resamples a pool of its draws with the weights of eq. (8), in the same way as the matched simulator distribution is built.

The baseline therefore states the form of growth, lodging, covariate effects, and conditioning, whereas the neural processes learn these forms from the data. It is strong where its forms match the data, and on the synthetic data it also profits from the evaluation itself (Appendix N).

### 7.3 Dataset construction

Each synthetic trajectory is generated from a *condition*, a unique pairing of synthetic genotype and year-site. The training sets form nested subsets of increasing size. Table 4 summarizes the training and evaluation set sizes.

**Table 4:** Synthetic dataset sizes. Conditions are unique pairings of synthetic genotype and year-site.

| Set | Genotypes | Year-sites | Conditions |
| --- | --- | --- | --- |
| *Training: one sample per condition* |  |  |  |
| 1k | 64 | 16 | 1,024 |
| 8k | 256 | 32 | 8,192 |
| 65k | 1,024 | 64 | 65,536 |
| 524k | 4,096 | 128 | 524,288 |
| *Evaluation: 64 samples per condition* |  |  |  |
| Seen | 1,000 | 64 | 64,000 |
| Genotype | 1,000 | 64 | 64,000 |
| Environment | 1,000 | 32 | 32,000 |
| Unseen | 1,000 | 32 | 32,000 |

Evaluation splits are defined relative to the 65k training set and contain 64 lodging-resampled trajectories for each fixed genotype and year-site condition. These evaluation splits mirror those of *FIP1* [38], each measuring generalization to held-out genotypes, held-out environments, or both. They are labeled *Seen, Genotype, Environment*, and *Unseen*, denoting evaluation splits rather than covariate regimes, with held-out genotypes and environments disjoint from all training datasets. The *Seen* split reuses a subset of the training genotypes and environments, the *Genotype* split uses held-out genotypes with the same environments, the *Environment* split the same genotypes with held-out environments, and the *Unseen* split holds out both, making it the hardest and most relevant case.

### 7.4 Training

All variants are trained with the neural-process objective, drawing a fresh split of each trajectory into context, target, and held-out sets at every step (Section 5). One experiment compares the scoring-set constructions directly. Everywhere else the default is the context-excluded scoring set *S* = *T* \ *C*, which keeps the context points out of the likelihood loss.

Training length is measured by the number of complete trajectories seen during optimization, sweeping 100k, 1m, 3m, and 5m and defaulting to 3m otherwise. The dataset size is swept over the four training sets and defaults to 524k. Training to 1m accounts for most of the improvement, and beyond 8k conditions the scores improve only slightly (Appendix K).

Optimization uses AdamW with batch size 64, learning rate 10^*−*3^, and cosine decay, and the latent variants additionally anneal the *KL* weight from 0 to 0.5. So that one trained model serves every covariate and context regime at evaluation, the maximum context-set size is annealed from 0 to 50 and the genotype-parameter and temperature inputs randomly dropped during training. The remaining hyperparameters are listed in Appendix C.

### 7.5 Covariate regimes

For a given genotype-environment condition, the model may receive two covariates, the normalized genotype-parameter vector *g* and the hourly temperature trajectory *e*. Four covariate regimes are evaluated, listed in Table 5 and denoted ∅, *g, e*, and *g* + *e* throughout the results. They isolate what each input contributes.

**Table 5:** Covariate regimes used for prediction and evaluation.

| Regime | Supplied |
| --- | --- |
| ∅ | none |
| $g$ | genotype-parameter vector $g$ only |
| $e$ | temperature trajectory $e$ only |
| $g + e$ | genotype-parameter vector and temperature trajectory |

### 7.6 Context regimes

For each covariate regime, three choices of observed height context are evaluated (Table 6). The *no context* regime predicts the full trajectory from covariates alone, with no observed height. The *sparse context* regime mirrors a few additional measurements taken throughout the season. The *max-height context* regime supplies all pre-lodging observations, so the model must reconstruct the deterministic growth curve from these noisy points and predict lodging, the only remaining stochastic behavior.

**Table 6:** Context regimes used at prediction time.

| Context regime | Context set |
| --- | --- |
| *no context* | $C = \emptyset$ |
| *sparse context* | 1–10 observed height points before maximum height |
| *max-height context* | all observed height points up to and including maximum height |

For a given condition, the selected context points are fixed across all model samples. Appendix J compares randomly chosen context points with points chosen by the predictive uncertainty of the models.

### 7.7 Evaluation protocol

Because the datasets are synthetic, predictions can be evaluated against the underlying true simulator trajectories. Each prediction is scored against the matched simulator distribution of Section 6.1, over the noise-free trajectories consistent with the covariates and observed context the model was given.

On the model side, the mean of each predictive Gaussian is compared at the same time points, which excludes observation noise to match the noise-free simulator distribution. The latent models produce 64 such trajectories from independent latent draws, and the conditional *CNP* and *ACNP* one each.

Since evaluating the entire candidate pool with the signature kernel is too computationally expensive, each score is estimated by Monte Carlo sampling over fixed-size blocks of model and simulator trajectories (Appendix E). *Sig-MMD* is the primary metric, while *CSig-MMD* focuses the same comparison on the tail, in particular lodging.

In addition to the distributional scores, the lodging figures summarize the fraction of predicted mean trajectories classified as lodged and the corresponding absolute height drops. The classification uses a maximum-to-final-height drop rule, counting a predicted mean trajectory as lodged when its final height is at least 0.1 m, and at least 20%, below its maximum predicted height. These cutoffs are below the minimum simulated lodging drop, so they should detect all predicted lodging events.

### 7.8 Computational cost

All experiments run on a single NVIDIA RTX 4090. Training one model on 3m trajectories takes under 3 hours. Generating 64 trajectories for each of 1,000 conditions takes about 4 seconds, and scoring them with *Sig-MMD* and *CSig-MMD* about 11 seconds.

## 8 Results

All metric values are reported as squared divergences scaled by 10^3^, with lower values corresponding to smaller discrepancy from the matched simulator distribution.

### 8.1 Qualitative trajectory distributions under covariate regimes

Figures 5 and 6 show trajectory samples for the same genotype and year-site, aggregated per row over a covariate regime. The Ground Truth column shows the matched simulator distribution (Section 6.1), which contracts from the full spread of simulator trajectories at ∅ to a single near-deterministic curve at *g* + *e*, where only lodging still varies. Figure 5 uses each row’s covariate regime as model input, whereas Figure 6 always uses both covariates. The difference is clearest for the *CNP* model. In Figure 5, under specific covariate regimes, it produces a single predictive mean trajectory. In Figure 6, because it always receives both covariates, it produces different trajectories for each covariate pair, which are then aggregated per row. Figure 5 thus shows that the *CNP* has learned the mean for each covariate regime.

**Figure 6:**
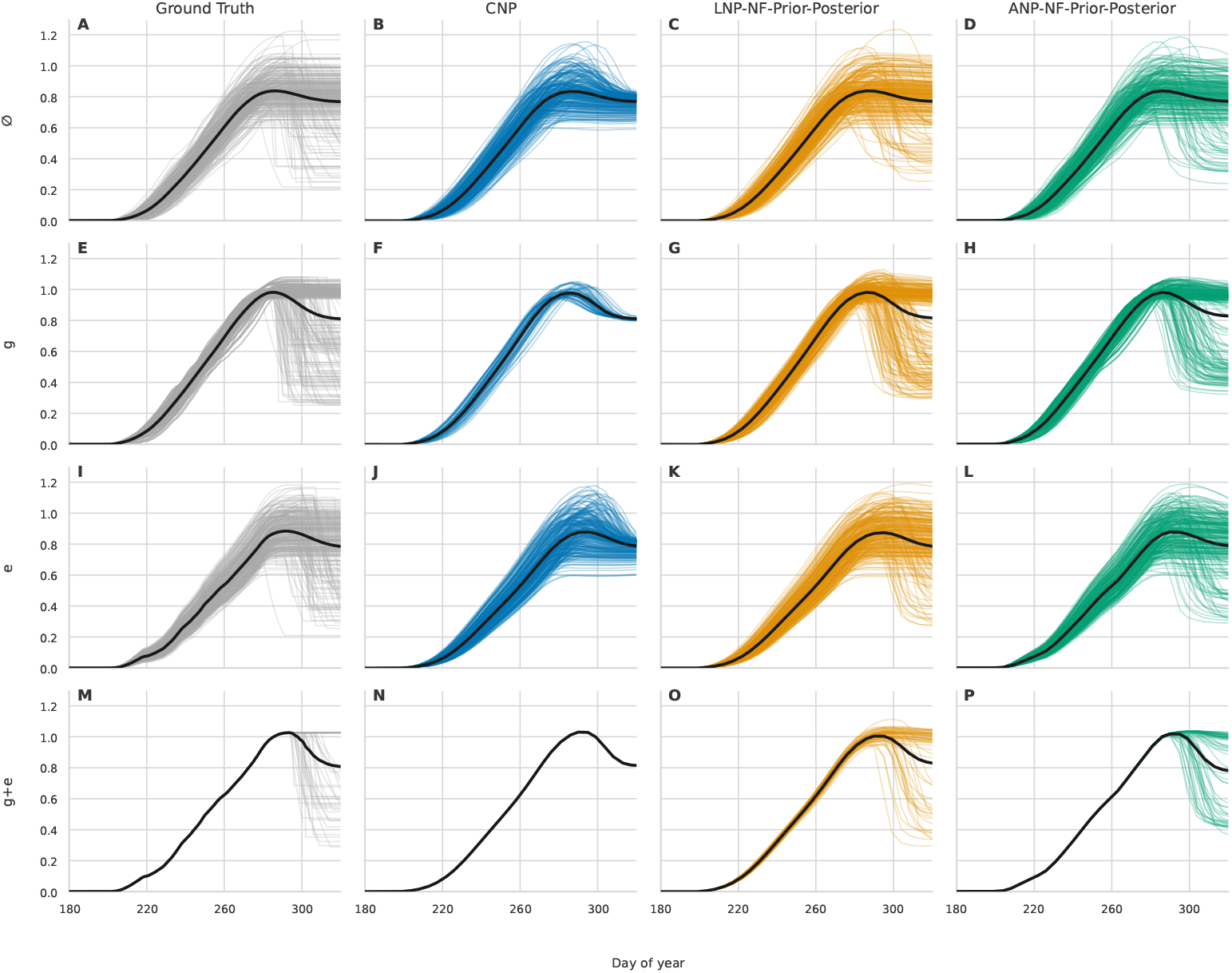
Height trajectory samples aggregated by covariate level while models receive the *g* + *e* covariate regime. Columns show the *Ground Truth*, the conditional neural process (*CNP*), and the latent (*LNP*) and attentive (*ANP*) neural processes with normalizing flows (*NF*) on both prior and posterior (*LNP-NF-Prior-Posterior, ANP-NF-Prior-Posterior*). The *Ground Truth* column shows synthetic trajectories from the simulator, and the other columns show model predictions. Rows aggregate trajectories over all conditions (∅), over the conditions of one genotype (*g*) or of one year-site (*e*), and over the single condition of this genotype and year-site (*g* + *e*), but every model receives both the genotype-parameter vector *g* and the temperature trajectory *e* of each condition. The genotype and year-site are selected near the 90th percentile of maximum height. No height context is given. Thin lines are individual samples, and thick lines are their means. Days are counted from the first of September. Since the *CNP* has no latent variable, it produces a single predictive mean trajectory for each genotype-environment pair. Because models always receive both inputs, trajectories vary within the coarser aggregation levels, except for *g* + *e*.

The *LNP-NF-Prior-Posterior* model matches the simulator distribution visually well, except for the last row with both covariates (Figure 5O), where it shows too much variance up to the maximum height of the trajectory. The *ANP-NF-Prior-Posterior* much better matches the deterministic simulator distribution in this case (Figure 5P), but in Figure 5 it does not match the simulator variance for either single-covariate regime (panels H and L), producing overly confident predictions. Figure 6, however, shows it predicts the correct distribution (panels H and L), indicating a collapse when given only partial covariates. This is also slightly less visible in the first row (Figure 5D against Figure 6D), where it does not predict many trajectories with maximum heights above one meter without covariates but does so correctly when they are given as input.

### 8.2 Scoring-set construction

This ablation varies the training scoring set *S*. Figure 7 illustrates overfitting to noisy context data. The models are given all noisy context points up to the simulator trajectory’s maximum height. The *ANP* model overfits to these context points under both scoring sets that include them (Figure 7E and G), where its predictions follow the noisy points instead of a smooth curve. The *LNP* model does not show the same issue, but its predictions are less confident overall, producing more varied trajectories.

**Figure 7:**
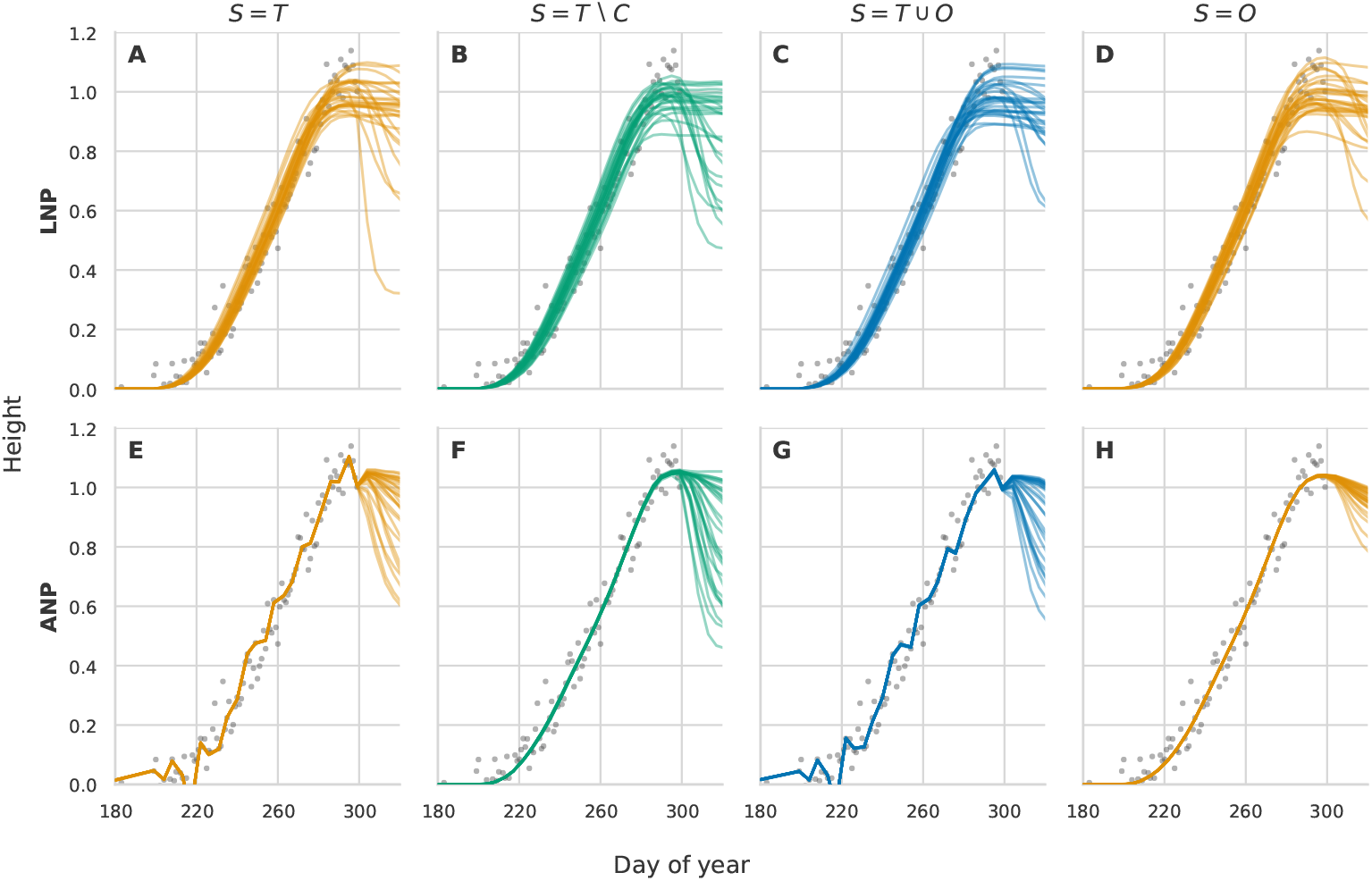
Height trajectory samples under different scoring-set constructions for the latent (*LNP*) and attentive (*ANP*) neural processes. Models receive the *max-height context* for one synthetic trajectory and no covariates (∅). Gray dots are the context points, noisy observations of one smooth simulator trajectory up to its maximum height. Lines show predicted trajectories under each scoring-set construction. The columns are the scoring sets *S* = *T, S* = *T* \ *C, S* = *T* ⋃ *O*, and *S* = *O*, where *T* is the target set, *C* the context set, and *O* a held-out set, so that the first and third column score the context points in the training loss and the other two exclude them. Days are counted from the first of September.

The same pattern appears quantitatively in Figure 8. When context points are scored, for *S* = *T* and *S* = *T* ⋃ *O*, the *ANP* model with context up to maximum height performs poorly (Figure 8B). Excluding the context, with *S* = *T* \ *C*, restores the expected benefit from additional context. The held-out constructions give no improvement and often degrade performance. *S* = *T* ⋃ *O* still scores the context and shows the same overfitting, while *S* = *O* drops the target set entirely to rely on the posterior channel alone and remains elevated. These results suggest that harmful in-context overfitting is mainly caused by the per-target context path. The posterior channel appears weaker because it is bottlenecked by *z* and regularized by the *KL* term. The context-excluded scoring set *S* = *T* \ *C* performs best, and is used in all subsequent experiments.

**Figure 8:**
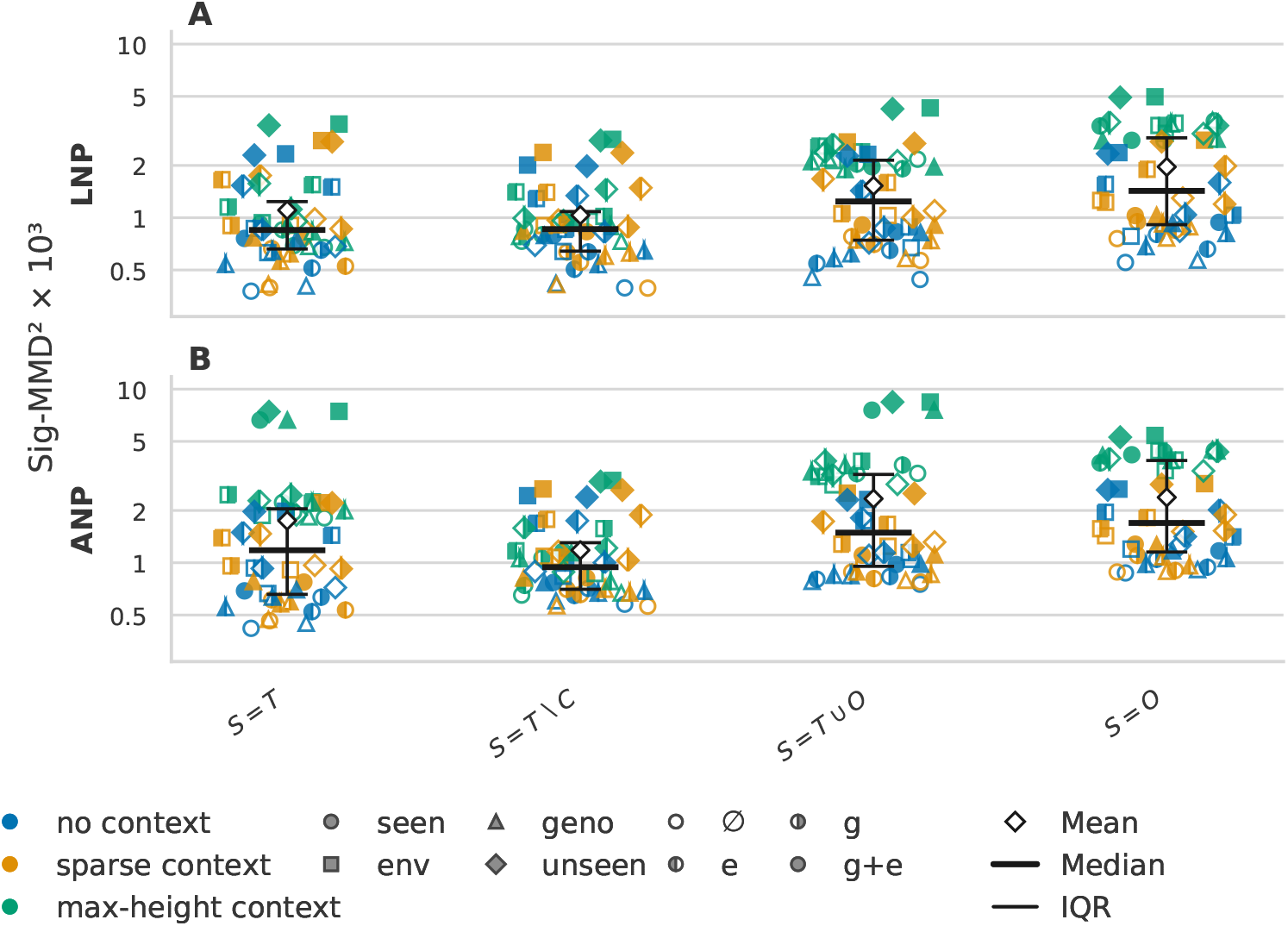
Signature maximum mean discrepancy (*Sig-MMD*) scores under the matched simulator distribution for the latent (*LNP*) and attentive (*ANP*) neural processes trained under different scoring-set constructions. The constructions are *S* = *T, S* = *T* \ *C, S* = *T* ⋃ *O*, and *S* = *O*, where *T* is the target set, *C* the context set, and *O* a held-out set. Each marker is the score of one combination of context regime (color), evaluation split (marker shape), and covariate regime (marker fill). The evaluation splits are seen genotypes and environments (*seen*), held-out genotypes (*geno*), held-out environments (*env*), and both held out (*unseen*). The covariate regimes are no covariates (∅), the genotype-parameter vector (*g*), the temperature trajectory (*e*), and both (*g* + *e*). Overlaid symbols give the mean, median, and interquartile range (*IQR*) over the markers of a scoring set. The vertical axis shows the squared *Sig-MMD* scaled by 10^3^ on a logarithmic scale. Lower is better.

### 8.3 Lodging probability and severity

The lodging measures use the ∅ covariate regime and compare the *no context* and *max-height context* columns. Lodging is investigated through the predicted lodging rate and severity. Figure 9 shows models trained on datasets with different lodging rates, the total percentage of lodged samples in each dataset. Because lodging in the synthetic generator depends only on the maximum height of the trajectory, the no-context and max-height-context rates should be very similar. Without context points, the *LNP* model matches the analytical rate quite well up to the 7% total lodging rate dataset (Figure 9A). The *ANP* model, however, does not predict lodging well without context and does not predict the full height range (Figure 9C). Both models predict similar rates with context, but these are worse than the *LNP* predictions without context. All subsequent experiments use a total lodging rate of *19*.*7%* for both training and evaluation sets.

**Figure 9:**
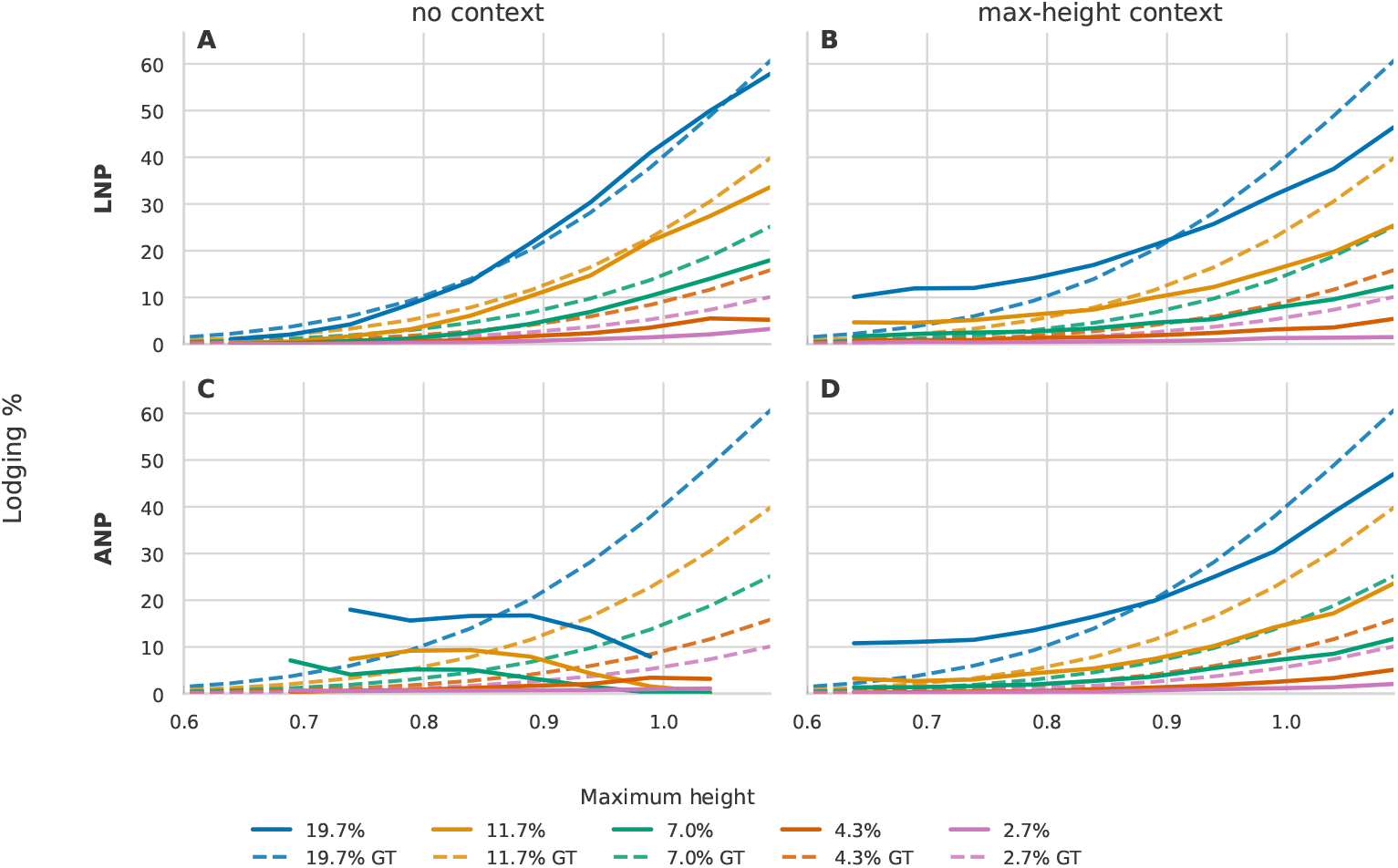
Lodging probability over maximum height for the global and per-target latent models, the latent (*LNP*) and attentive (*ANP*) neural processes, trained using different lodging scales. The legend lists the resulting overall lodging rate of the respective training set. Solid curves show lodging rates of the predicted trajectories. Predicted trajectories are counted as lodged using the maximum-to-final-height drop rule described in Section 7.7. Dashed curves show the analytical lodging probability of the respective synthetic generator, the ground truth (*GT*). Columns are context regimes. In the *no context* column models receive no height points. In the *max-height context* column they receive all noisy context points up to maximum height of the trajectory. Without context, trajectories are grouped by their predicted maximum height, and with context by their true maximum height in the simulator, in bins of 0.05 m. Bins with fewer than 1 % of the trajectories are not shown. All predictions use no covariates (∅).

So far the *LNP* and *ANP* models have been analyzed without *NF* extensions. Figure 10 shows the same lodging rate setup as Figure 9 for the expanded global and pertarget model groups. The global *LNP* and its flow variants again match the analytical lodging rate quite well without context (Figure 10A), while the per-target *ANP* and its flow variants do not (Figure 10C). For the per-target models, *NF*s can improve lodging prediction without context, but still do not capture the full height range. The conditional *CNP* and *ACNP* models, which have no latent variable, emit a single mean trajectory. Their curves show the fraction of predictive mean trajectories classified as lodged once the ratio of lodged samples bends the mean downward enough.

**Figure 10:**
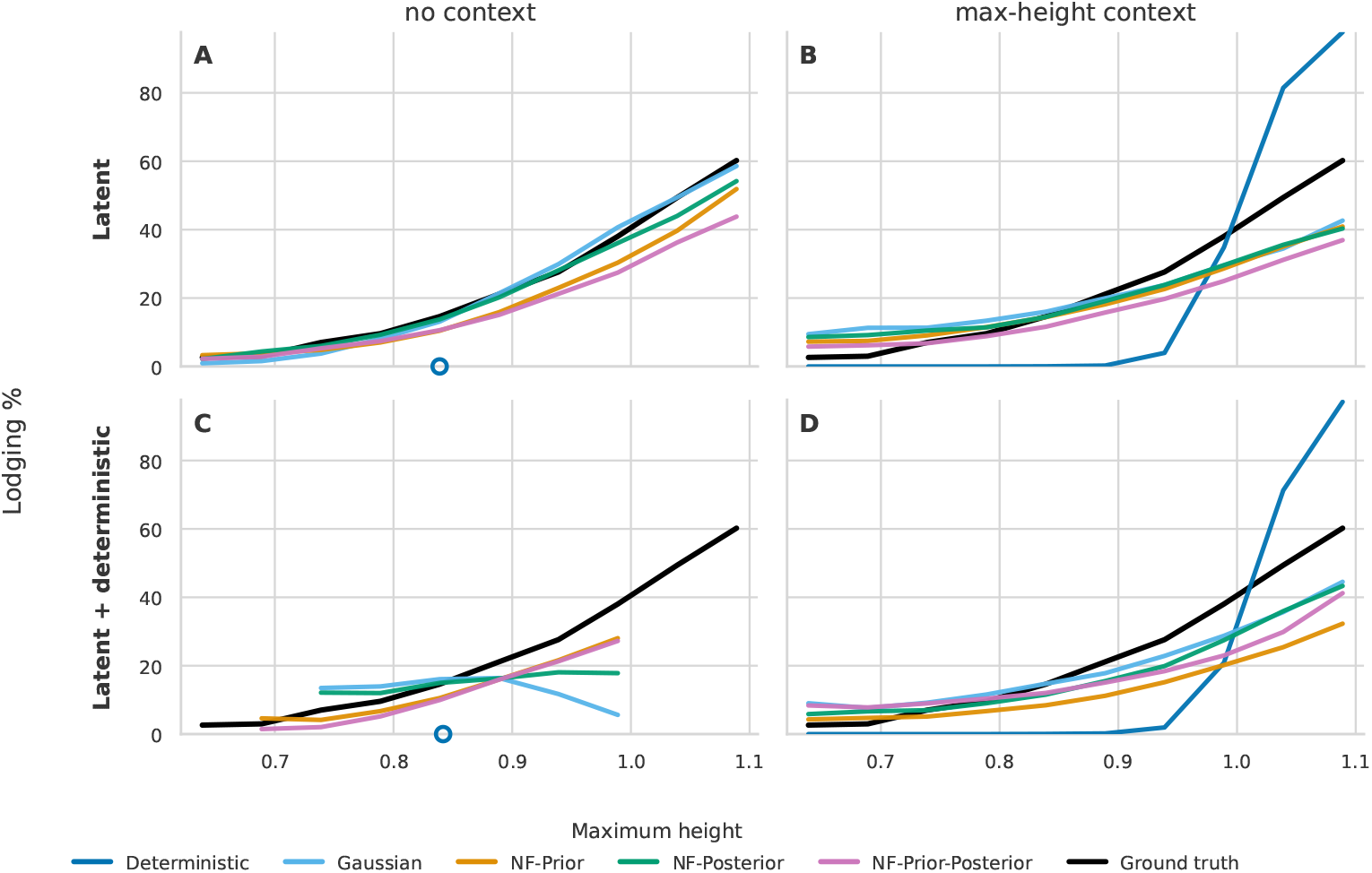
Lodging probability over maximum height for the global and per-target model groups. In the synthetic generator, lodging probability increases with maximum height. Predicted trajectories are counted as lodged using the maximum-to-final-height drop rule described in Section 7.7. The *Latent* row contains the global models, the conditional (*CNP*) and latent (*LNP*) neural processes and the *LNP* variants with normalizing flows (*NF*), *LNP-NF-\**, and the *Latent + deterministic* row the per-target models, the attentive conditional (*ACNP*) and attentive (*ANP*) neural processes and *ANP-NF-\**. Within each row, *Deterministic* denotes the conditional model (*CNP*/*ACNP*), *Gaussian* the latent model (*LNP*/*ANP*), and *NF-\** its flow variants, with a normalizing flow on the prior (*NF-Prior*), the posterior (*NF-Posterior*), or both (*NF-Prior-Posterior*). Columns are context regimes. In the *no context* column models receive no height points. In the *max-height context* column they receive all noisy context points up to maximum height of the trajectory. The black *Ground truth* curve is the lodging rate of the simulator trajectories. Without context, trajectories are grouped by their predicted maximum height, and with context by their true maximum height in the simulator, in bins of 0.05 m. Bins with fewer than 1 % of the trajectories are not shown. All predictions use no covariates (∅). For conditional models, the curve is the fraction of predictive mean trajectories classified as lodged. Without context, the *CNP* and *ACNP* predict one trajectory for all conditions, shown as one open circle at its maximum height.

Figure 11 shows lodging severity as the mean absolute drop from maximum to lodged height. In the simulator the final lodged height is a uniformly sampled fraction of maximum height, so the absolute drop increases linearly with maximum height. The *LNP* variants reproduce this linear relationship well, whereas the *ANP* variants do not without context (Figure 11C) and only approach it given max-height context (Figure 11D).

**Figure 11:**
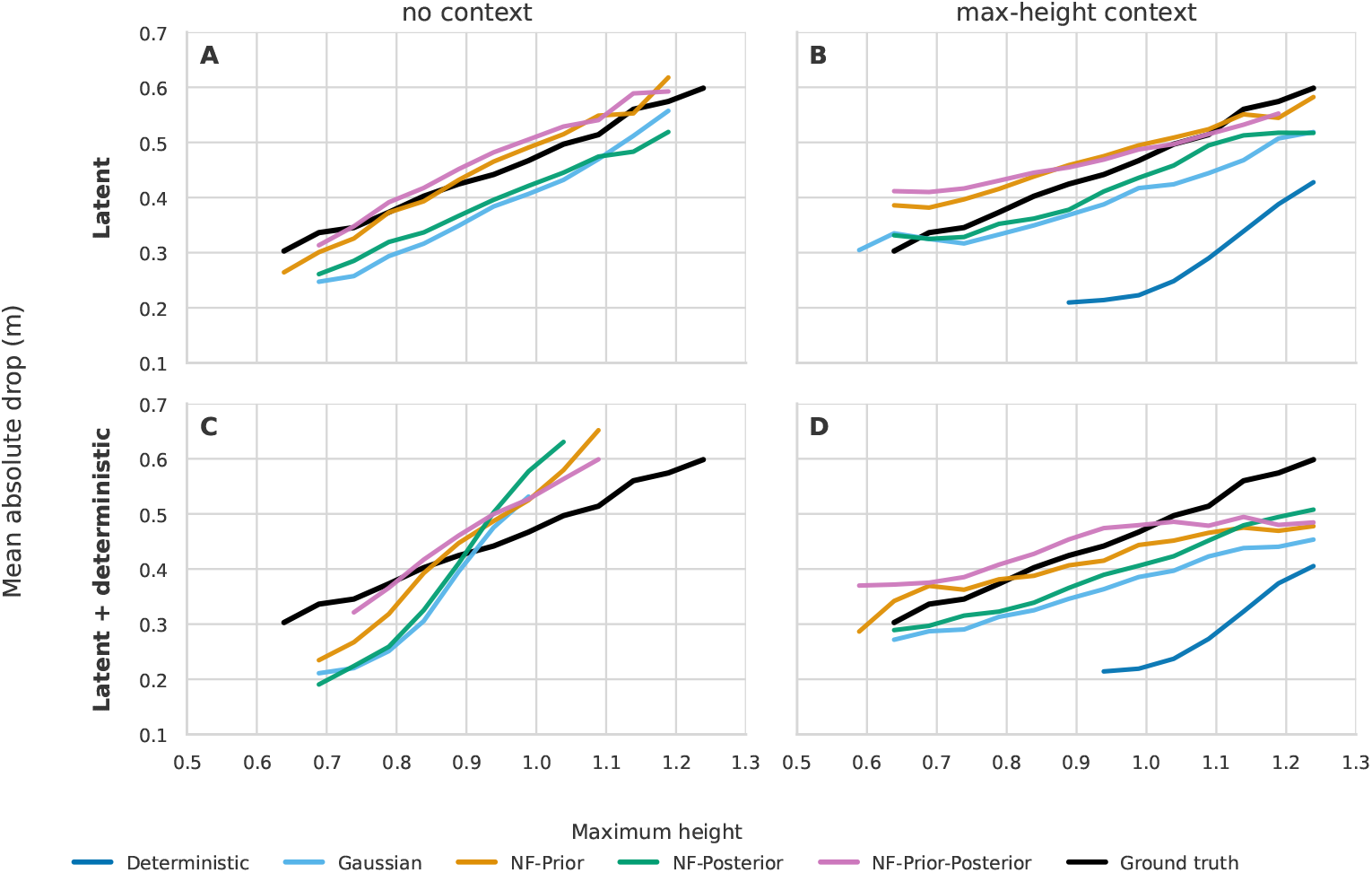
Lodging severity over maximum height for the global and per-target model groups. In the simulator, the final lodged height is sampled as a fraction of maximum height, so the expected absolute drop increases with maximum height. The plotted drop is the maximum-to-final-height drop averaged over trajectories counted as lodged. The *Latent* row contains the global models, the conditional (*CNP*) and latent (*LNP*) neural processes and the *LNP* variants with normalizing flows (*NF*), *LNP-NF-\**, and the *Latent + deterministic* row the per-target models, the attentive conditional (*ACNP*) and attentive (*ANP*) neural processes and *ANP-NF-\**. Within each row, *Deterministic* denotes the conditional model (*CNP*/*ACNP*), *Gaussian* the unflowed latent model (*LNP*/*ANP*), and *NF-\** its flow variants, with a normalizing flow on the prior (*NF-Prior*), the posterior (*NF-Posterior*), or both (*NF-Prior-Posterior*). Columns are context regimes. In the *no context* column models receive no height points. In the *max-height context* column they receive all noisy context points up to maximum height of the trajectory. The black *Ground truth* curve is the mean drop of the lodged simulator trajectories. Without context, trajectories are grouped by their predicted maximum height, and with context by their true maximum height in the simulator, in bins of 0.05 m. Bins with fewer than 25 lodged trajectories are not shown. All predictions use no covariates (∅). For conditional models, the drop is measured on the predictive mean trajectory. Without context, the *CNP* and *ACNP* predict no lodged trajectory and therefore have no curve.

### 8.4 Distributional scores

This experiment compares the model variants under the matched simulator distribution (Section 6.1), with each score computed within a fixed setting defined by the evaluation split, covariate regime, and context regime. Figure 12 reports the overall *Sig-MMD* over the context regimes, evaluation splits, and covariate regimes. The latent models clearly improve on the conditional *CNP* and *ACNP*, with substantially lower scores. The baseline scores below every neural process in most settings, in particular on unseen year-sites, whereas the neural processes score below it with both covariates on seen year-sites and, with max-height context, with both covariates on every split (Appendix N). Each model was trained five times with different randomness, and the score figures summarize all five runs, so that a difference between models can be compared with the variation between runs (Appendix O). The prior flow lowers the *Sig-MMD* of its base model in every setting, by 27 to 40 % without context and with sparse context, and the difference is significant in every context regime except for the *ANP-NF-Prior* with max-height context. The posterior flow alone lowers it by 8 to 9 % in the same regimes, which is close to the variation between runs. The *LNP* scores lower than the *ANP* in every context regime, significantly without context and with sparse context, and its prior-flow variants score below those of the *ANP* in these regimes, by more than three standard deviations over runs without context and by 1.6 to 2.3 with sparse context (Table 14). Within each family, the two prior-flow variants differ by less than the variation between runs.

**Figure 12:**
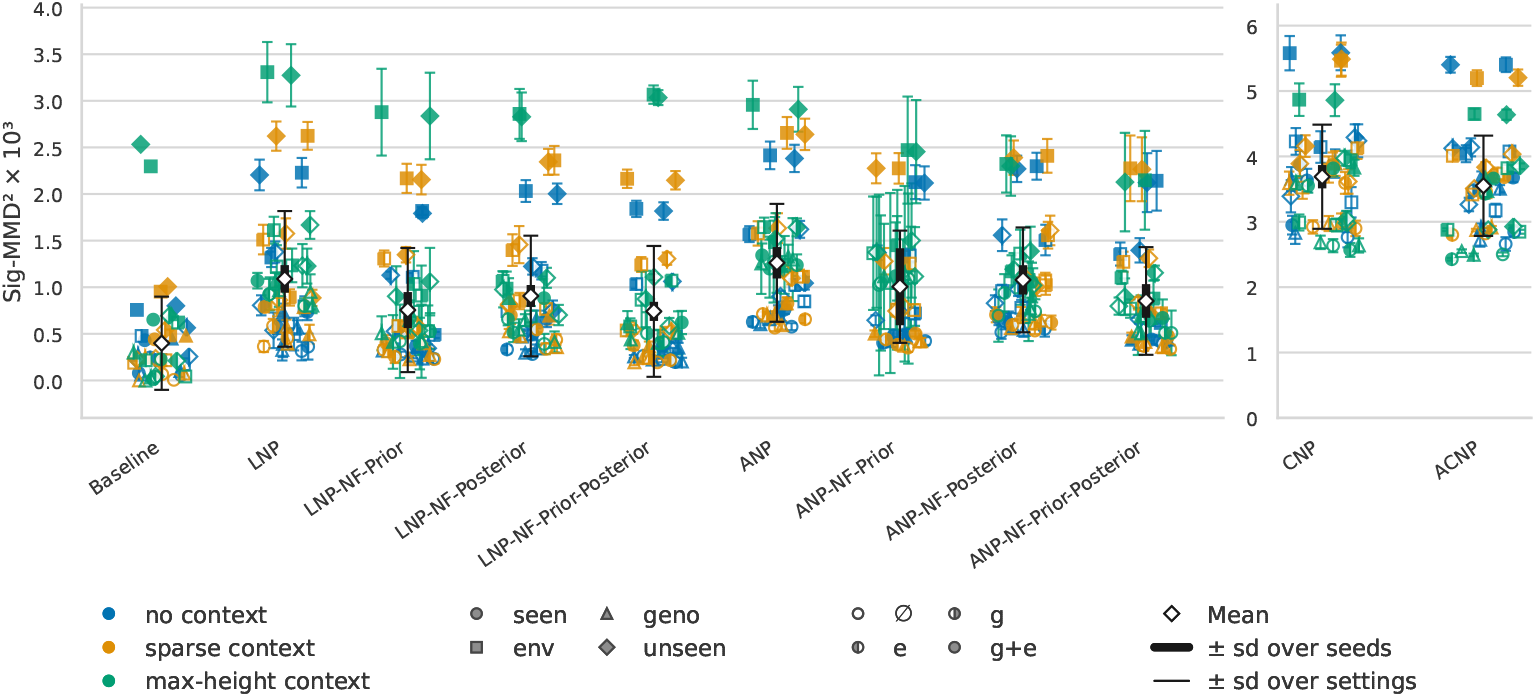
Signature maximum mean discrepancy (*Sig-MMD*) scores under the matched simulator distribution, across model variants, evaluation splits, covariate regimes, and context regimes. The models are the conditional (*CNP*), attentive conditional (*ACNP*), latent (*LNP*), and attentive (*ANP*) neural processes. Variants with a normalizing flow (*NF*) on the prior, the posterior, or both carry the suffixes *NF-Prior, NF-Posterior*, and *NF-Prior-Posterior*. The evaluation splits are seen genotypes and environments (*seen*), held-out genotypes (*geno*), held-out environments (*env*), and both held out (*unseen*). Scores compare models within fixed covariate and context regimes. *Baseline* is the lodging-mixture spline model with genotype and environment effects, shown as a single fit. Each marker is the score of one combination of context regime (color), evaluation split (marker shape), and covariate regime (marker fill), as the mean over five training runs, with a bar of ±1 standard deviation (*SD*) over the runs. The covariate regimes are no covariates (∅), the genotype-parameter vector (*g*), the temperature trajectory (*e*), and both (*g* + *e*). The diamond is the mean over all settings of a model. The thick bar spans ± the root mean square (*RMS*) of the per-setting *SD*s over runs (*sd over seeds*), and the thin bar ±1 *SD* over the settings (*sd over settings*). The *CNP* and *ACNP* are shown in the right panel with their own vertical axis. The vertical axes show the squared *Sig-MMD* scaled by 10^3^. Lower is better.

The *CSig-MMD* evaluation, shown in Figure 13, makes the variant differences in the tail more pronounced than *Sig-MMD*, and it also spreads the training runs of a model more widely. On the tail score, the baseline is lowest in most settings without *g*, whereas with both covariates on seen year-sites the *LNP-NF-Prior* is lower in every context regime. Among the latent models, *LNP-NF-Prior* has the lowest tail score in most settings and in every setting with *g*, and its prior flow is the only one that lowers the tail score of its base model significantly in every context regime, by 20 to 43 % (Appendix O). Without *g* and with context, the *ANP* and its variants with a posterior flow are among the lowest. The other flows change the tail score of their base model in both directions. The posterior flow of the *ANP* lowers it significantly without context and with sparse context, whereas the two *ANP* variants with a prior flow score significantly higher than the *ANP* in up to eight settings of a regime. The order of the flow variants on the tail therefore depends on the covariate and context regime. The order of the *LNP* and the *ANP* on the tail score also depends on the tail threshold, since a higher threshold keeps only the most severe lodging in the tail, whereas the order of *LNP-NF-Prior-Posterior* and *ANP-NF-Prior-Posterior* holds at every threshold and kernel bandwidth tested (Appendix F).

**Figure 13:**
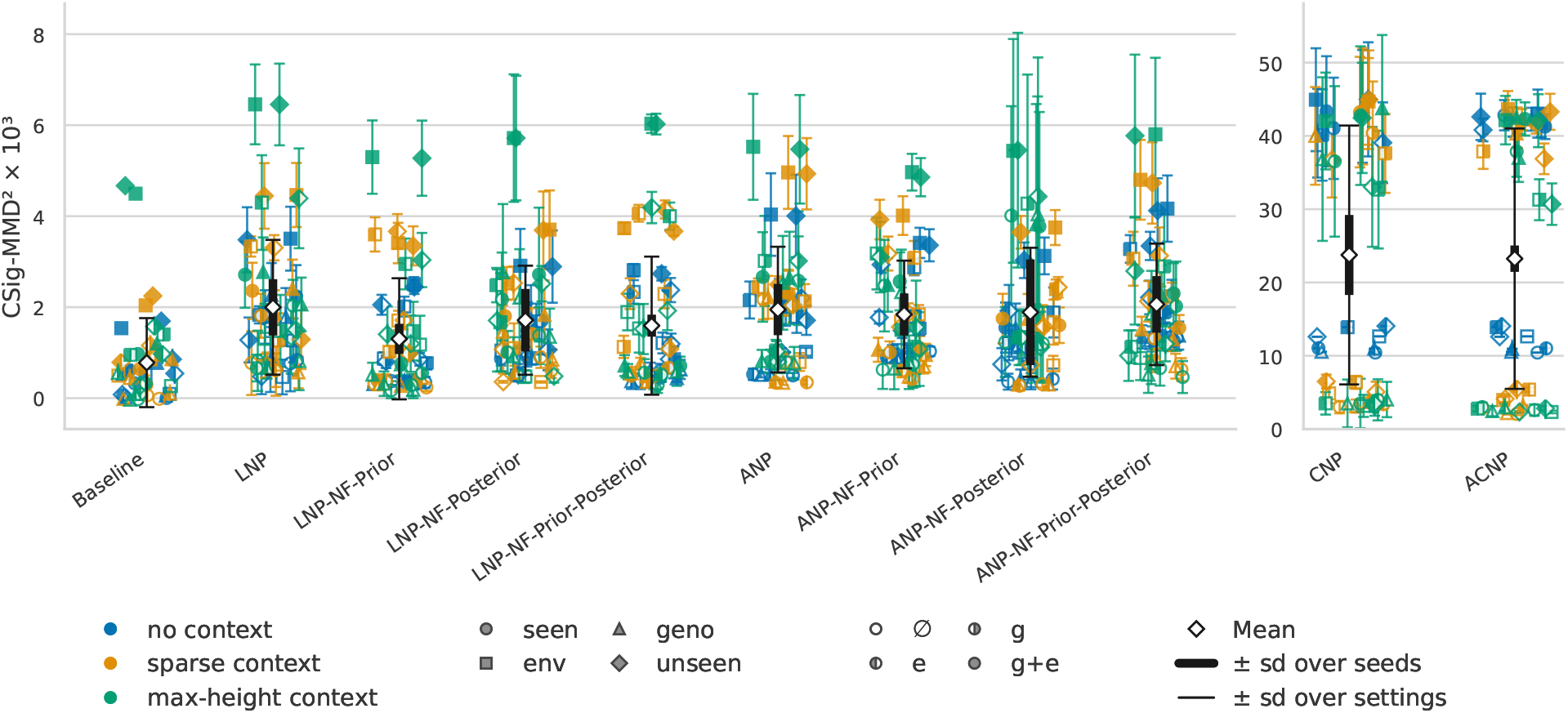
Censored signature maximum mean discrepancy (*CSig-MMD*) scores under the matched simulator distribution, across model variants, evaluation splits, covariate regimes, and context regimes. *CSig-MMD* concentrates the comparison on atypical trajectories, in particular lodged ones. The models are the conditional (*CNP*), attentive conditional (*ACNP*), latent (*LNP*), and attentive (*ANP*) neural processes. Variants with a normalizing flow (*NF*) on the prior, the posterior, or both carry the suffixes *NF-Prior, NF-Posterior*, and *NF-Prior-Posterior*. The evaluation splits are seen genotypes and environments (*seen*), held-out genotypes (*geno*), held-out environments (*env*), and both held out (*unseen*). Scores compare models within fixed covariate and context regimes. *Baseline* is the lodging-mixture spline model with genotype and environment effects, shown as a single fit. Each marker is the score of one combination of context regime (color), evaluation split (marker shape), and covariate regime (marker fill), as the mean over five training runs, with a bar of ±1 standard deviation (*SD*) over the runs. The covariate regimes are no covariates (∅), the genotype-parameter vector (*g*), the temperature trajectory (*e*), and both (*g* + *e*). The diamond is the mean over all settings of a model. The thick bar spans ± the root mean square (*RMS*) of the per-setting *SD*s over runs (*sd over seeds*), and the thin bar ±1 *SD* over the settings (*sd over settings*). The *CNP* and *ACNP* are shown in the right panel with their own vertical axis. The vertical axes show the squared *CSig-MMD* scaled by 10^3^. Lower is better.

Figure 9 showed that the *LNP* and the *ANP* underestimate the lodging probability at lower lodging rates. This holds for all models (Appendix P). At a lodging rate of 2.7 %, the latent models lodge 0.6 to 1.7 % of the plants and the conditional models almost none. The scores at this rate therefore compare models that all miss most of the lodging, and their order does not follow how much lodging a model reproduces. Under *CSig-MMD*, only the last two places of the conditional models hold at every rate. At the observation noise of *FIP1*, 0.02 instead of 0.05 m, the *LNP*, the *ANP*, and their *NF-Prior-Posterior* variants keep their *Sig-MMD* order, and the two flow variants rank first and second under both scores.

### 8.5 Application to FIP1

The models were also trained on the *FIP1* plant-height data and evaluated on held-out plots, held-out genotypes, and the held-out year 2019, with the observed plots as the reference (Appendix Q). Figure 22 shows samples without context next to the measured plots. With max-height context, the mean score over models and covariate regimes is 2.24 on held-out plots and 0.42 on held-out genotypes, the latter at the resolution of the metric of about 0.4, whereas in the held-out year it remains at 4.77 for seen and 3.39 for held-out genotypes. With context, the *ANP* variants score below the *LNP* variants in every covariate regime (Table 18), whereas without context and covariates the *LNP* variants score lowest. Initializing from the checkpoints of the synthetic study instead of at random lowers the mean score in every evaluation split and context regime (Table 19), and the baseline is at or below the best neural process in every combination of context and covariate regime except without context and covariates. *CSig-MMD* is not reported on *FIP1*, since its tail is defined by the lodging events of the simulator, whereas on *FIP1* lodging is known only from a detector that flags 1.6 % of the plots, and the trained models predict at most 0.6 %.

## 9 Discussion and Conclusion

Modeling wheat height as a distribution over growth trajectories captures the variation of the growth process that a single curve discards. We studied this problem with *NP*s and their *NF* extensions on a synthetic generator calibrated against Swiss weather records and the *FIP1* dataset. Its ground truth allows predicted and simulated trajectory distributions to be compared directly with *Sig-MMD* and *CSig-MMD*, with lodging as the tail event. The same models were then trained and evaluated on *FIP1* itself.

Only the latent *LNP* and *ANP* capture this distribution, whereas the conditional models reduce to a single mean trajectory. The per-target conditioning of the *ANP* variants sharpens the prediction where the covariates nearly determine the trajectory. Under partial covariates, the same conditioning makes them overconfident, and without context they miss the lodging rate of the simulator. The global *LNP* reproduces this rate and scores lower on *Sig-MMD*. The prior flow lowers the *Sig-MMD* of its base model in both families, which is consistent with the prior, conditioned on less information, needing the more expressive distribution. On the tail score, only the prior flow of the *LNP* improves its base model in every context regime, and *LNP-NF-Prior* is lowest in most settings. The posterior flow helps in some regimes although it is used only in training, which indicates that a better posterior fit can improve the learned prior. The training objective also matters. Scoring the observed context points leads to overfitting of their noise, and excluding them from the scoring set avoids this, which matters for the noisy and sparse measurements of phenotyping.

These conclusions rest on five training runs per model. On *Sig-MMD*, the gains of the prior flows exceed the variation between runs. On the tail score, most differences among the flow variants do not, and their order changes with the covariate and context regime. The tail order of the *LNP* and the *ANP* depends on the threshold, whereas that of their *Prior-Posterior* variants holds at every threshold and kernel bandwidth tested. The *Sig-MMD* order of these four models also holds at the observation noise of *FIP1*. All rankings were established at a lodging rate far above that of real wheat. At realistic rates, every model misses most of the lodging and the rankings change, so they characterize the models where lodging is frequent enough to be learned.

The lodging-mixture spline baseline scores below the neural processes in most synthetic settings. Its functional forms match the generator, and wherever the genotype or year-site was seen in training, or no covariate is given, it looks up a fitted effect instead of predicting one. The neural processes improve on it with both covariates on seen year-sites, with max-height context and both covariates on every evaluation split, and on *FIP1* in a single combination of context and covariate regime. Their advantage is therefore not a lower score in this study. It is a single amortized model that conditions on any subset of covariates and context without refitting and represents the growth process in its latent space, whose principal components modulate the maximum height and the late drop of lodging (Appendix L).

The application to *FIP1* makes the distance to real data measurable. On the training seasons, the predictions with max-height context reach the resolution of the metric on held-out genotypes and remain above it on held-out plots. This was reached with one million training samples from 2,930 plots after pre-training on the simulator, in line with the synthetic scaling (Appendix K). The heldout year remains about ten times above the resolution, and since 2019 is the only held-out year, a property of that season cannot be separated from a general limitation. With context, the *ANP* variants score below the *LNP* variants on *FIP1* in every covariate regime, the reverse of the synthetic order, and we have no explanation for this. The experiment further covers one dataset, and the reported spread reflects the variation between training runs, not between datasets. Lodging, which separates the real data most from the synthetic setting, is too rare in *FIP1* to be learned or to be evaluated per condition.

The generator is deliberately simple. It models a single trait from a temperature-only environment, represents the genotype by a parameter vector, and sets the lodging model rather than fitting it. Across genotypes, its spread of growth-cessation day is too wide and its spread of elongation rate too narrow compared with *FIP1*. Both the generator and the models assume homoscedastic Gaussian noise, whereas real measurement noise is heterogeneous, real sampling is sparser and more irregular, and devices, years, and sites introduce systematic shifts. The synthetic results identify the modeling choices that decide whether a model captures a stochastic event such as lodging, but the ranking they establish need not carry over to real data, as *FIP1* shows.

These limitations define the role of the simulator. Field data provide one realization per plot and few replicate plots per genotype and season, so the distribution a model should predict for a condition is unknown, and a predicted distribution can only be checked against the marginal over the observed plots. The simulator provides this distribution per condition. Whether a model captures a stochastic event at all, and under which covariates and context, can therefore be established in simulation, and a rare event such as lodging can be evaluated on its tail, which the real data do not allow. Testing the models in simulation before they are trained on real data, as in this study, separates the question whether a model class captures the event from the question whether the data suffice to learn it. The simulator further serves as a prior for the real data, since initializing from the synthetic checkpoints lowered the mean score on *FIP1* in every evaluation split and context regime. Evaluation can draw on both sources, with the simulator providing the replicates per condition that a tail score requires and the real data the check of the marginal distribution.

Several directions follow from these results. Held-out seasons require training data from several years and platforms, so that the variation between seasons is part of the training distribution. Lodging calls for targeted collection or augmentation of lodging events and for methods from imbalanced learning. Beyond height, more detailed environmental covariates and multiple correlated traits are where signatures and *NF*s could prove most useful. Together, the results indicate that *NP*s paired with signature-based distributional metrics offer a way to model and evaluate the variability of a plant trait and to capture it in a latent representation.

## A Synthetic Height Generator Details

Sampled free control points are clipped elementwise to [max(*M −* 4σ_cp_, 0), *M* + 4σ_cp_]. This clipping is part of the generative process rather than a training-time constraint. The height time axis uses the 274-day weather window starting at day 61 plus a 30-day flat post-season extension. Unless stated otherwise, synthetic datasets use the height-generator parameters listed in Table 7. One B-spline surface sampled around the calibrated mean is visualized in Figure 2.

**Table 7:**
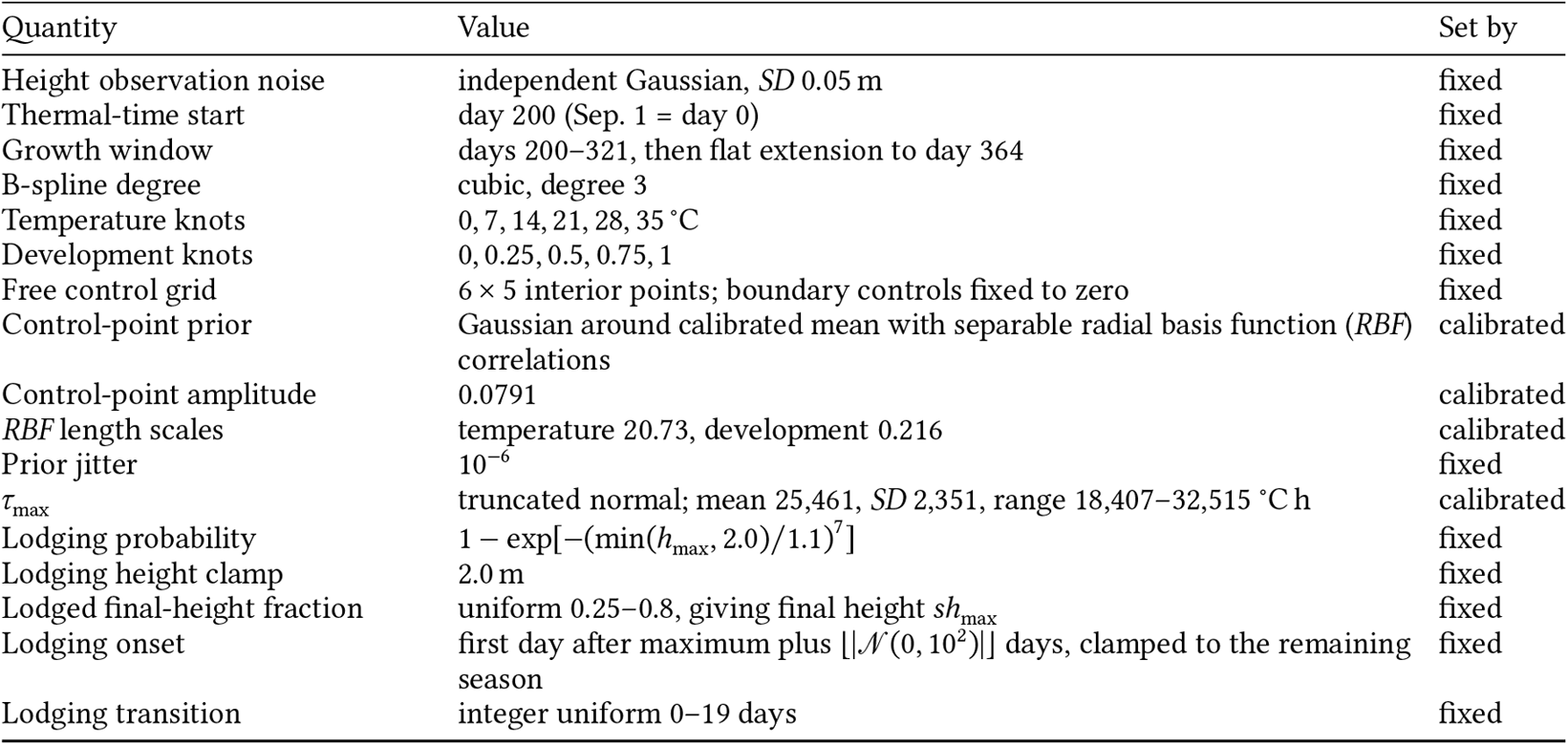
Default synthetic height-generator parameters. Distribution widths are given as standard deviation (*SD*). The last column states whether a value was calibrated or fixed.

Table 7 marks each parameter as calibrated or fixed. The fixed values reflect the simplifying assumptions of the generator. Thermal time is accumulated from day 200 (20 March), before stem elongation begins in *FIP1*, to day 321, around the last *FIP1* height measurements. The generator does not represent vernalization or photoperiod, so thermal time accumulated over a mild winter would advance development unchecked. The knots are equally spaced, in temperature between the growth limits of 0 and 35 ^°^C and in development on [0, 1]. They fix only the resolution of the surface, and its shape is learned in calibration. The observation-noise *SD* of 0.05 m is about twice the *FIP1* measurement noise of about 0.02 m, so that the treatment of noise by the models can be assessed. The lodging parameters are fixed so that the lodging rate can be set and varied in the lodging-scale experiment. *FIP1*, with about 1.6 % lodged plots, contains too few events to constrain them. The Weibull shape gives a probability that rises steeply with maximum height, in line with the observed events, and the scale sets the overall rate.

### A.1 Comparison with Observed Trajectories

Table 8 compares simulated and observed trajectories per year on the shared measurement grid of Appendix B. Figure 14 shows the trajectories of each year. The simulated trajectories include lodging and observation noise, with the lodging rate and the noise level set to match *FIP1*, and each simulator score is the mean over five realizations of both. Because the absolute scores are hard to interpret on their own, the table also gives two references computed from the observed plots. Within-year scores compare disjoint genotype groups of the same year and between-year scores compare the plots of two years, so the two references bracket the discrepancy to be expected within and between seasons.

**Table 8:** Agreement between the calibrated height generator and the Field Phenotyping Platform 1.0 (*FIP1*) dataset per harvest year, as squared signature maximum mean discrepancy (*Sig-MMD*) scaled by 10^3^. The simulator column scores simulated against observed trajectories of the year, as mean and standard deviation (*SD*) over five lodging and noise realizations. The remaining columns score observed trajectories against each other. Diagonal entries compare disjoint genotype groups of the same year, as mean and *SD* over 50 splits, and off-diagonal entries compare the plots of two years.

| Year | Simulator | 2016 | 2017 | 2018 | 2019 | 2021 | 2022 |
| --- | --- | --- | --- | --- | --- | --- | --- |
| 2016 | $53.704 \pm 0.231$ | $0.126 \pm 0.258$ | 42.870 | 85.234 | 33.241 | 99.203 | 58.089 |
| 2017 | $2.783 \pm 0.092$ | 42.870 | $0.158 \pm 0.286$ | 20.724 | 10.455 | 27.424 | 23.765 |
| 2018 | $0.397 \pm 0.029$ | 85.234 | 20.724 | $0.131 \pm 0.204$ | 19.194 | 11.791 | 8.593 |
| 2019 | $6.862 \pm 0.172$ | 33.241 | 10.455 | 19.194 | $0.239 \pm 0.342$ | 19.679 | 11.856 |
| 2021 | $2.619 \pm 0.073$ | 99.203 | 27.424 | 11.791 | 19.679 | $0.126 \pm 0.180$ | 19.361 |
| 2022 | $3.091 \pm 0.056$ | 58.089 | 23.765 | 8.593 | 11.856 | 19.361 | $0.122 \pm 0.281$ |

**Figure 14:**
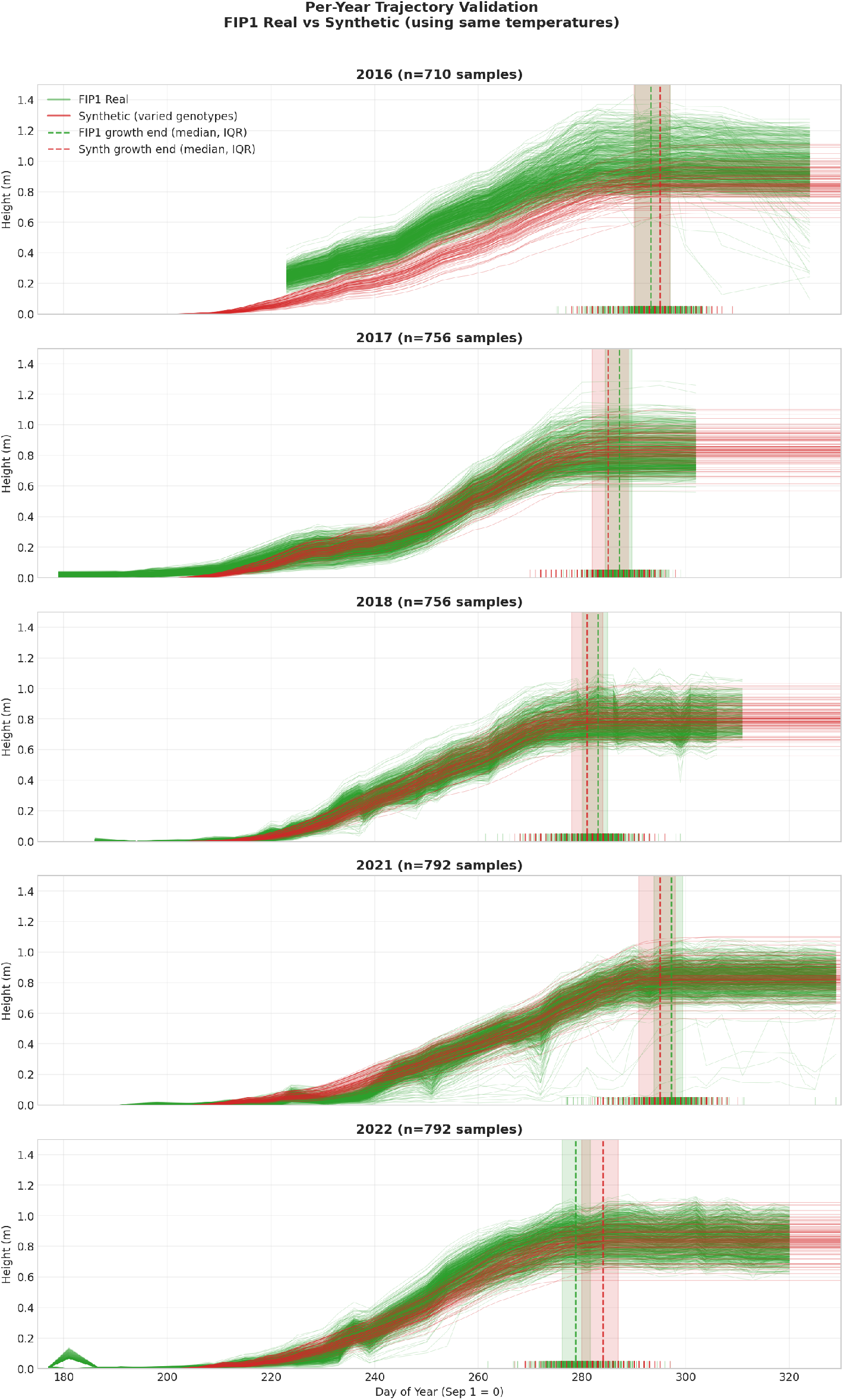
Per-year validation of the synthetic height generator against the Field Phenotyping Platform 1.0 (*FIP1*) dataset. Green curves show observed *FIP1* height trajectories and red curves show synthetic trajectories generated using the corresponding yearly temperature inputs and sampled genotypes. Ticks at the bottom mark the growth-cessation day of each trajectory, and dashed lines and shaded bands their median and interquartile range (*IQR*). The held-out year 2019 is not shown.

The height distributions were fitted on 2017, 2018, 2021, and 2022, whereas the 2016 plots entered only the growth-cessation term of the calibration objective. In these four years, the simulator scores lie between the two references. In 2016 and 2019, the observed plants are already taller than the simulated ones at the first grid date, and this initial difference is included in the scores. In 2016 it is large enough that the simulator score exceeds every between-year reference among the four fitted years.

### A.2 Genotype Variability

Each simulated genotype is a vector of control points drawn from the calibrated prior. Since simulated genotypes do not correspond to *FIP1* cultivars, the prior is assessed through the traits it produces. For each *FIP1* harvest year, 1,000 genotypes are drawn from the prior and simulated with the recorded temperatures of that year, without lodging and observation noise. Their spread across genotypes is compared with that of the *FIP1* genotypes of the same year in three traits of a trajectory, its maximum height, its growth-cessation day, and its elongation rate. The growth-cessation day is defined as in the calibration objective. In the simulator it is the day on which growth stops, and in *FIP1* it is the day on which the smoothed trajectory levels off. The elongation rate is the mean height gain per day while the trajectory rises from 40 % to 80 % of its maximum height.

Simulated genotypes have no replicate plots and no field effects, so the *FIP1* reference is the spread across genotypes with replicate noise removed. For maximum height, *FIP1* provides a spatially corrected best linear unbiased estimate (*BLUE*) per genotype and a generalized heritability per year [38]. The heritability is the fraction of the variance of the *BLUE*s that is due to genotype, and the reference is the *SD* of this genotypic part. For the other two traits, the reference is the *SD* of the genotype means with the variance between replicate plots of the same genotype subtracted. Figure 15 shows the ratio of simulated to *FIP1* spread per year, with bootstrap intervals over genotypes, and Table 9 gives the underlying values.

**Table 9:** Standard deviation (*SD*) across genotypes of maximum height, growth-cessation day, and elongation rate per harvest year, for the Field Phenotyping Platform 1.0 (*FIP1*) dataset and the calibrated height generator. Observed values are the spread across *FIP1* genotypes with replicate noise removed. Simulated values are over 1,000 genotypes drawn from the calibrated prior and simulated with the recorded temperatures of the year. The year 2019 was not used in the calibration.

| Year | Maximum height (cm) |  | Growth-cessation day (days) |  | Elongation rate (mm/day) |  |
| --- | --- | --- | --- | --- | --- | --- |
|  | *FIP1* | Simulator | *FIP1* | Simulator | *FIP1* | Simulator |
| 2016 | 11.87 | 11.09 | 4.77 | 5.76 | 2.66 | 1.47 |
| 2017 | 10.77 | 11.05 | 3.28 | 5.02 | 2.72 | 2.15 |
| 2018 | 7.82 | 9.42 | 2.47 | 5.28 | 1.64 | 1.50 |
| 2019 | 7.21 | 11.11 | 3.16 | 4.79 | 1.65 | 2.03 |
| 2021 | 8.59 | 10.97 | 3.32 | 5.10 | 2.05 | 1.60 |
| 2022 | 8.98 | 10.72 | 3.81 | 5.21 | 3.61 | 1.73 |

**Figure 15:**
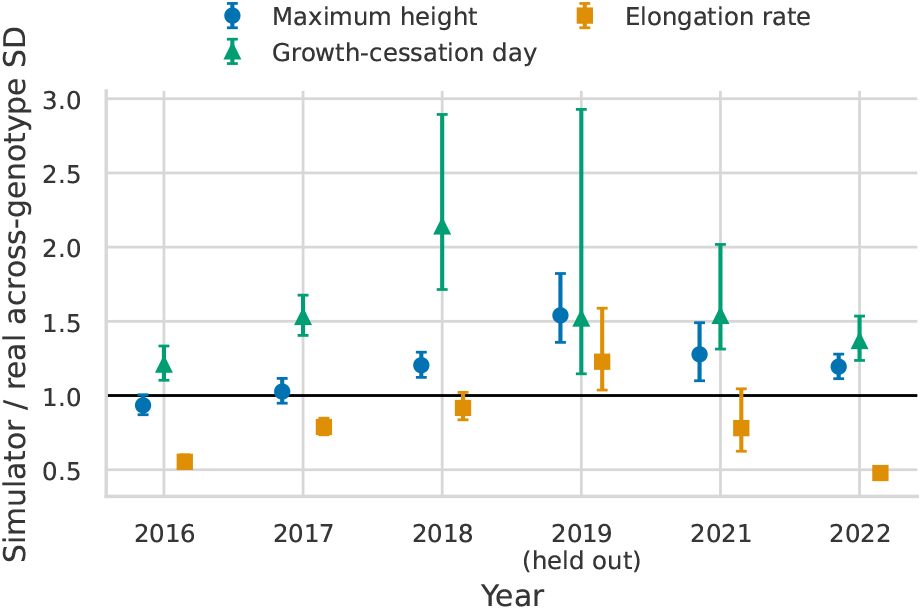
Ratio of the simulated to the observed standard deviation (*SD*) across genotypes of maximum height, growth-cessation day, and elongation rate, per harvest year of the Field Phenotyping Platform 1.0 (*FIP1*) dataset. Simulated values are from 1,000 genotypes drawn from the calibrated prior and simulated with the recorded temperatures of the year. Observed values are the spread across *FIP1* genotypes after removing replicate noise. Error bars are 95 % percentile intervals from 1,000 bootstrap resamples of observed and simulated genotypes. The horizontal line marks equal spread. The year 2019 was not used in the calibration.

The simulated spread of maximum height is close to the *FIP1* spread in the calibration years, whereas the spread of growth-cessation day is too wide in every year and the spread of elongation rate too narrow in most years. The excess spread is consistent with the calibration matching the spread of plots, which includes variation between replicate plots that the simulator can only represent as variation between genotypes. Elongation rate is the most informative of the three traits, as it was not a direct calibration target.

Rank stability is compared with the Spearman correlation of maximum height between the genotypes of two years, over the 15 pairs of *FIP1* years. Simulated values receive noise matched to the precision of the *BLUE*s, so that replicate noise affects both correlations alike. The mean correlation is 0.93 for *FIP1* and 0.96 for the simulator, so simulated rankings are slightly more stable across years than observed ones. The comparison tests populations rather than individual genotypes, and the spread of each trait rather than the shape of its distribution or the correlations between traits.

## B Shared FIP1 Measurement Grid

The *FIP1* year-sites have different measurement schedules and share no measurement date, so comparisons with observed trajectories place all plots on one grid of 12 dates from day 223 to day 302. Each year-site serves a grid date with its nearest measurement date, at most 3 days away. Only the time coordinate of a measurement moves, its height stays as measured. This shift displaces the heights by about 0.017 m root mean square (*RMS*), below the observation noise of about 0.02 m. Plots without a measurement on one of the selected dates are dropped.

## C Training Hyperparameters

The training loop consumes 3,000,000 trajectory samples. AdamW uses learning rate 10^*−*3^, weight decay 0.01, betas (0.95, 0.98), gradient clipping at 0.1, and a cosine learning-rate schedule with a 3,072-sample warmup.

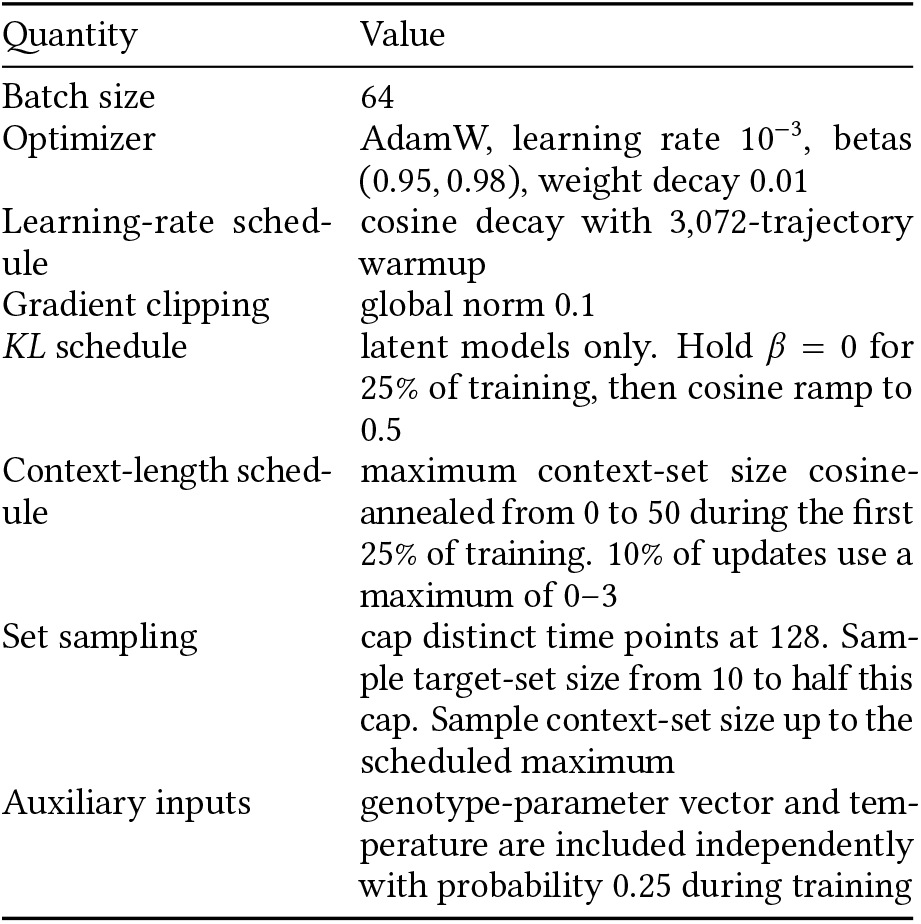

| Quantity | Value |
| --- | --- |
| Batch size | 64 |
| Optimizer | AdamW, learning rate $10^{-3}$, betas (0.95, 0.98), weight decay 0.01 |
| Learning-rate schedule | cosine decay with 3,072-trajectory warmup |
| Gradient clipping | global norm 0.1 |
| KL schedule | latent models only. Hold $\beta = 0$ for 25% of training, then cosine ramp to 0.5 |
| Context-length schedule | maximum context-set size cosine-annealed from 0 to 50 during the first 25% of training. 10% of updates use a maximum of 0–3 |
| Set sampling | cap distinct time points at 128. Sample target-set size from 10 to half this cap. Sample context-set size up to the scheduled maximum |
| Auxiliary inputs | genotype-parameter vector and temperature are included independently with probability 0.25 during training |

## D Architecture Details

All variants share the transformer backbone and covariate encoders. They differ along the conditional/latent and global/per-target axes and, for the flow variants, in the latent flows. In the per-target models, observation and covariate tokens cannot attend to query tokens, and query tokens attend to all non-query tokens and themselves but not each other, which keeps the queried points conditionally independent. The per-target representations 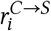 are produced only in the context pass. Because the target pass has no query tokens, the target observations *y*_*T*_ never enter them.

D Architecture Details
| Component | Configuration |
| --- | --- |
| *Shared transformer backbone* |  |
| Hidden width | 128 |
| Transformer blocks | 12, pre-norm |
| Attention heads | 4 (head dim 32) |
| Normalization | root mean square layer normalization (*RMSNorm*) |
| Feed-forward | Swish-gated linear unit (*SwiGLU*), inner width 4× hidden |
| Input embeddings | bias-free linear $1 \rightarrow 128$ for $x$ and $y$. Context token = $x\text{-emb} + y\text{-emb}$ |
| Prefix tokens | learnable *global* token; *flow-condition* token (flow variants) |
| *Covariate encoders* |  |
| Temperature | hierarchical 1D-conv stem (hourly→daily kernel/stride 24, weekly kernel 7, strided pooling) to 6 tokens + learnable positional encoding; input $274 \times 24 = 6,576$ |
| Genotype-parameter vector | multi-layer perceptron (*MLP*) $31 \rightarrow 64 \rightarrow 64 \rightarrow 128$ |
| *Latent head* |  |
| Latent dimension | 32 |
| Latent encoder | *MLP* $128 \rightarrow 2 \times 32$; scale = $0.1 + 0.9 \sigma(\cdot)$ |
| Latent decoder | *MLP* $32 \rightarrow 128$ |
| Output decoder | 2-layer *MLP* + linear $\rightarrow 1$ |
| Likelihood | homoscedastic Gaussian, single learned variance |
| Latent samples | 8 latent draws during training; up to 64 draws at prediction and evaluation (deterministic variants effectively one draw). |
| *Latent flows (NSFs)* |  |
| Prior flow | coupling *NSF* with 6 layers, 8 bins, tail bound 5, conditioner *MLP* width 64 |
| Posterior flow | autoregressive *NSF* with 3 transforms, 8 bins, conditioner width (64, 64) |
| Flow-condition input | *MLP* $128 \rightarrow 128 \rightarrow 32$ from the flow-condition token |

## E Evaluation Metric Details

Unless otherwise stated, the synthetic metrics use up to 64 stored trajectories per condition. The prediction day axis is normalized by the simulator height day range. Deterministic *CNP* and *ACNP* models produce one trajectory per model input. Blocks with several such inputs are empirical distributions over distinct trajectories. Both metrics compare trajectories through the signature kernel. Each height trajectory is divided by a fixed scale, lifted with a lead-lag transform, and augmented with a normalized-time channel that maps days to [0, 1], so that the one-dimensional height curve retains its full shape. *Sig-MMD* estimates the squared *MMD* (eq. (5)) with the cosine-normalized signature kernel. The metric uses the unbiased U-statistic for every block with at least two trajectories. Only singleton blocks use the diagonal kernel term. This rule also applies to deterministic *CNP* and *ACNP* predictions.

*CSig-MMD* adds a tail stage. Each trajectory is described by a truncated signature, normalized by its own kernel norm to remove the size effect, and its robust Mahalanobis distance to the reference distribution is computed from an *MRCD* estimate fitted on pooled normalized signatures of simulator draws for the training conditions. The *MRCD* regularizes the robust scatter toward a diagonal target, so the distance stays well-defined although the signatures have directions of near-zero variance. The tail threshold is the distance that keeps 97.5 % of the lodged training draws in the tail, and *c* denotes its square. The quantile α of all training distances that this corresponds to depends on the lodging rate of the training data, and is 0.81 at the default rate. Censoring acts on each trajectory’s signature-kernel embedding rather than on the path itself. The embedding is a logistic blend between the trajectory’s own embedding and that of a flat zero-height pivot **0**. Equivalently, expectations under the censored distribution

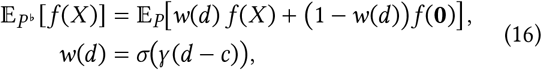

where *d* is the trajectory’s squared tail distance and *c* the squared threshold distance. The weight *w* grows smoothly with *d*, so trajectories far into the tail, where *d* ≫ *c*, keep their contribution while typical non-tail trajectories are down-weighted toward the pivot, with *w* = ^1^ at the threshold. The censored *MMD* uses the same signature kernel as *Sig-MMD*.

Figure 16 visualizes the *CSig-MMD* tail construction. The robust signature-Mahalanobis distance separates lodged trajectories from the main non-lodged band. For non-lodged trajectories, the distance also rises at the lowest and highest maximum heights, showing sensitivity to unusually short or tall growth curves. The same score can therefore serve as a practical outlier score for growth trajectories.

**Figure 16:**
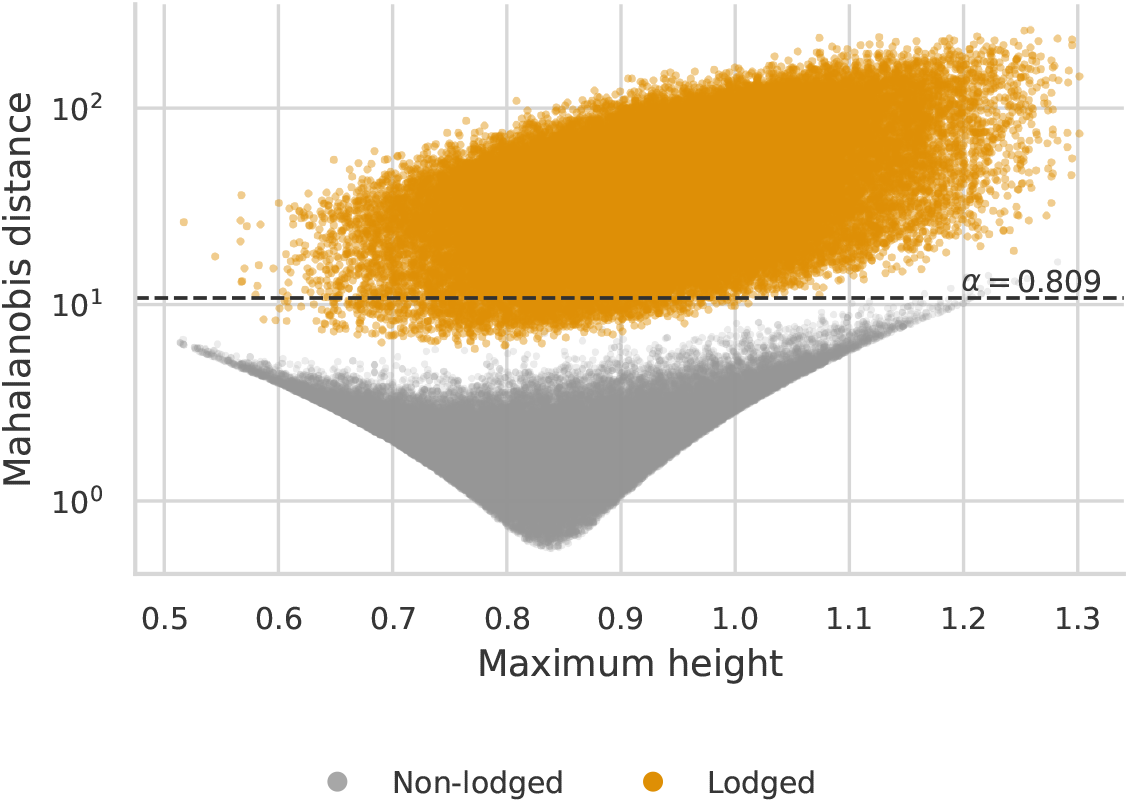
*Signature-Mahalanobis separation* for simulator trajectories. The vertical axis shows the robust Mahalanobis distance in signature space as a function of maximum height. The dashed line marks the tail threshold used by the censored signature maximum mean discrepancy (*CSig-MMD*), labeled with its quantile α of the training distances at the default lodging rate. Lodged trajectories form a high-distance tail. The non-lodged band also rises at low and high maximum heights, showing sensitivity to unusually short or tall trajectories.

Estimating the matched simulator distribution of Section 6.1 exactly would mean comparing, for each covariate regime, the model’s pooled predictions against simulator draws from the entire candidate pool. Each score is therefore estimated by Monte Carlo sampling over fixed-size blocks. Within a covariate pool, a balanced block of model trajectories is repeatedly drawn across the pool’s conditions, paired with an equally sized block of trajectories from the matched simulator distribution, scored by the *Sig-MMD* of the block, and averaged over blocks. The block size and the number of blocks per covariate regime bound the number of kernel evaluations while covering the pool. The simulator trajectories are drawn from the posterior-weighted mixture, whose context weights use the height-observation noise scale σ_obs_ of Table 7. The block size is capped by the draws available per condition. For *g* + *e*, this gives one model draw for deterministic variants and up to 64 for latent variants. The simulator side uses 64 draws. All block sampling is deterministically seeded.

| Quantity | Value |
| --- | --- |
| *Shared signature kernel* |  |
| Static kernel | radial basis function (RBF), $\sigma = 1$ |
| Partial differential equation (PDE) dyadic order | 2 |
| Path lift | lead-lag on height + normalized-time channel |
| Height scale | fixed 1.5 m, shared across conditions |
| Kernel normalization | cosine, $k(a, b)/\sqrt{k(a, a)k(b, b)}$ |
| *MMD* estimator | U-statistic for blocks with at least two trajectories and diagonal term for singleton blocks |
| *Matched simulator distribution* |  |
| Observation-noise scale | $\sigma_{\text{obs}} = 0.05$ m (Table 7) |
| Block size | 512 per side, model vs. simulator |
| Blocks per covariate regime | 1000 for ∅, 16 per environment for $e$, 1 per genotype for $g$, 1 per condition for $g + e$ |
| Stored draws per condition | up to 64 model draws, one for *CNP* and *ACNP*, and 64 simulator draws |
| *CSig-MMD tail* |  |
| Tail features | depth-4 truncated signatures, level 0 dropped, kernel-normalized |
| Robust scatter | *MRCD*, support fraction 0.8, fit on $\leq 10^4$ pooled simulator signatures of the training conditions |
| Tail threshold | keeps 97.5 % of lodged training draws in the tail ($\alpha = 0.81$ quantile of training distances at the default lodging rate) |
| Censoring weight | logistic $\sigma(\gamma(d - c))$ with boundary steepness $\gamma = 25.6/c$, so $w$ rises from 0.12 to 0.88 within $\pm 8\%$ of $c$ |
| Pivot | flat zero-height path |

## F Sensitivity of the Metrics to their Hyperparameters

Both metrics depend on the bandwidth σ of the static kernel, and *CSig-MMD* also on the tail threshold *c* and the steepness γ of the censoring boundary, which is set relative to the threshold as γ *c*. The paper uses σ = 1, the threshold at the quantile α = 0.81 of the training distances, and γ *c* = 25.6. We rescored the *LNP*, the *ANP*, and their *NF-Prior-Posterior* variants without context in the 16 settings of the four evaluation splits and four covariate regimes at σ = 0.5, 1, and 2, at α = 0.81, 0.85, and 0.95, which keep 97.5 %, 76.5 %, and 25.5 % of the lodged training draws in the tail, and at γ *c* = 12.8, 25.6, and 51.2. Table 10 ranks the four models in each setting and gives the mean rank over the settings. The steepness is not tabulated, since it changes a score by at most 5 % in the median and the order of the models in at most five settings. Under *Sig-MMD*, all 16 differences between the *LNP* and the *ANP* exceed twice their standard error over the paired block scores, and the *LNP* is lower in 14. Under *CSig-MMD*, 13, 14, and 16 differences are resolved in this sense at the three thresholds, and the *LNP* is lower in 7, 12, and 16.

**Table 10:**
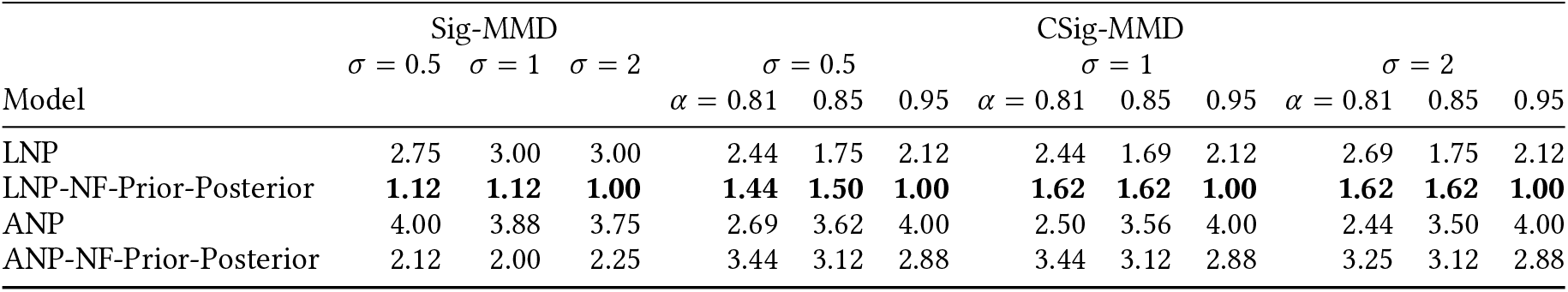
Mean rank of four models under the signature maximum mean discrepancy (*Sig-MMD*) and its censored variant (*CSig-MMD*) at three bandwidths σ of the static kernel and, for *CSig-MMD*, three tail thresholds, given as the quantile α of the training distances. In each of the 16 settings of the four evaluation splits and four covariate regimes without context, the models are ranked from 1 (lowest score) to 4, and the table gives the mean over the settings. The models are the latent (*LNP*) and attentive (*ANP*) neural processes and their variants with a normalizing flow (*NF*) on the prior and the posterior (suffix *NF-Prior-Posterior*). The steepness of the censoring boundary is γ *c* = 25.6. Bold marks the best mean rank in each column.

## G Temperature Generation Details

The generated observations additionally include output measurement noise with standard deviation 0.3 ^°^C.

The annual harmonic amplitude and phase receive site/year adjustments, while the semiannual component is shared. The hourly AR(1) weather anomaly is mixed between a year-shared component and site-local component, then soft-capped with *a* tanh(*W* /*a*) using the calibrated anomaly soft cap. The seasonal component *T*_seasonal_ follows a two-harmonic Fourier model whose amplitude and phase vary by site and year, capturing regional differences and inter-annual variability. Weather variability *T*_weather_ is an hourly *AR(1)* process. A single shared hourly series *W*_*y*_(*d, h*) is generated per year. Each site combines it with an independent local hourly process 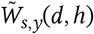:

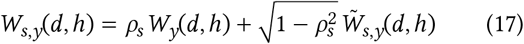

where ρ_*s*_ is a per-site correlation coefficient.

The diurnal component *T*_diurnal_ produces an asymmetric daily cycle. Let *u* = (*h − h*_trough_) mod 24 be the time since the daily trough, and let Δ_*r*_ and Δ_*f*_ = 24 *−* Δ_*r*_ be the rise and fall durations.

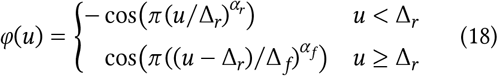

Since φ is asymmetric, its daily average 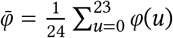 is generally nonzero, so we center the diurnal component as

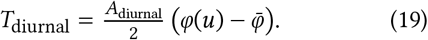

This spans the full daily range *A*_diurnal_ with a smooth analytical mean correction, so it leaves the daily-mean seasonal component *T*_seasonal_ unbiased. The rise/fall exponents are seasonally interpolated calibrated values. The defaults are approximately 1.044/0.754 for winter/summer rise and 0.509/0.768 for winter/summer fall.

### G.1 Calibration

The temperature model is calibrated against 151 complete station-year records from eight winter wheat variety testing sites, Changins, Delley, Ellighausen, Grangeneuve, Lindau, Sulz-Künten, Vouvry, and Zollikofen, over harvest years from 2004 to 2024. The temperature parameters are calibrated in stages. The seasonal harmonics are fitted to observed daily means. The *AR(1)* persistence coefficient is then fitted to the residual daily mean anomalies after removing the seasonal cycle. Diurnal timing and waveform shape parameters are extracted from the observed hourly cycle. The remaining variance parameters governing site and year variability are optimized using Optuna [1] to match marginal statistics of the observed temperatures.

## H Synthetic Lodging Details

The analytical lodging curves use the same probability function as the synthetic generator. Given maximum height *h*, the lodging probability is

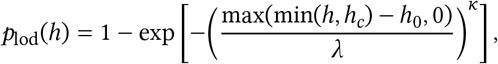

where *h*_*c*_ is the height clamp, *h*_0_ is the height offset, λ is the scale, and *κ* is the Weibull shape parameter. In the default dataset we use *h*_*c*_ = 2.0 m, *h*_0_ = 0, λ = 1.1 m, and *κ* = 7. The lodging-scale experiment varies λ while keeping the other parameters fixed.

If lodging is sampled, let *d*_max_ be the first maximum-height day. The onset offset is Δ = ⌊|*N* (0, 10^2^)|⌋ days, so lodging starts at *d*_*l*_ = *d*_max_ + 1 + Δ, and the event is recorded only when this day lies within the simulated season. We then sample a final-height fraction *s* ∼ Uniform(0.25, 0.8), giving final lodged height *sh*_max_, and a transition duration *L* ∼ Uniform{0, …, 19} days, clamped to the available days after onset. Before *d*_*l*_, the trajectory is unchanged. During the transition it decays exponentially toward *sh*_max_, and after the transition it remains at this height. When *L* = 0, the final height applies only from the onset day onward. This sampled event is used as the simulator lodging label.

## I Condition-Specific Scoring

The condition view compares every prediction to the exact simulator distribution for the same genotype and year-site pair. This condition-view score is distinct from the matched and context-matched blocked views, which pool conditions by covariate scope before constructing fixed-size MMD blocks.

The main results score each prediction against the matched simulator distribution (Section 6.1), whose spread contracts as covariates are supplied. Because that reference differs across covariate regimes, those scores cannot be compared across them. Here we instead score every prediction against the individual condition it was generated from, *P*_sim_(**H** ∣ *u*), regardless of which covariates the model received. The reference is then the same single condition across all covariate regimes, so the scores become comparable across them. A model given no covariates must cover the full simulator spread and is scored against this single condition, while a model given both covariates can concentrate on it. The scores therefore reflect how much each covariate helps the model predict the actual trajectory.

Figure 17 shows these scores for the latent models. Scores fall as more covariate information is supplied, from the ∅ regime through the single-covariate regimes to *g* + *e*, so the models do use the additional covariates to predict the specific condition more closely. Added height context lowers the scores within each covariate regime in the same way, with the ∅ no-context predictions highest and the *g* + *e* max-height-context predictions lowest. The flow variants are again among the best, and the global *LNP* variants tend to score lower than their per-target *ANP* counterparts.

**Figure 17:**
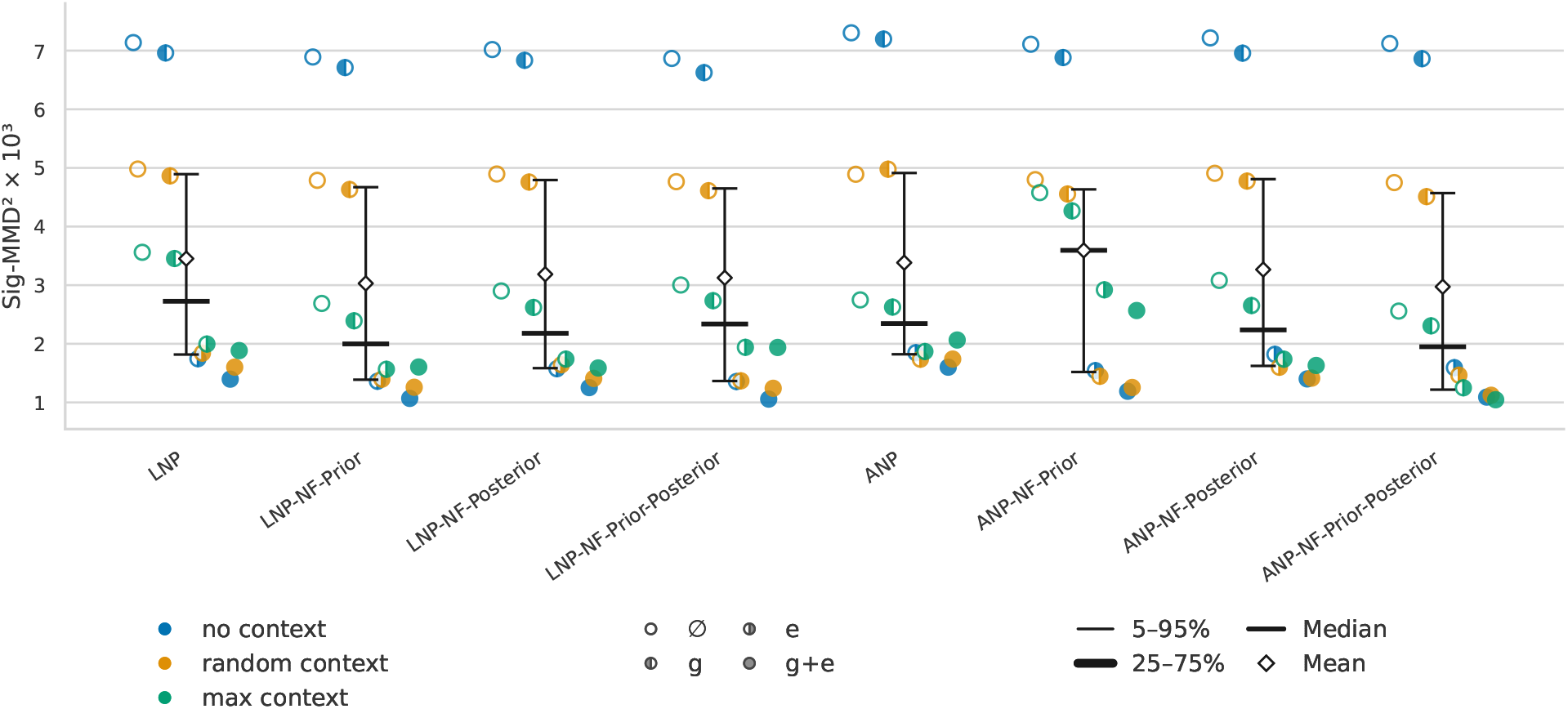
Condition-specific signature maximum mean discrepancy (*Sig-MMD*) scores for the latent models, with each prediction scored against the individual condition it was generated from rather than the matched simulator distribution. The models are the latent (*LNP*) and attentive (*ANP*) neural processes. Variants with a normalizing flow (*NF*) on the prior, the posterior, or both carry the suffixes *NF-Prior, NF-Posterior*, and *NF-Prior-Posterior*. Each marker is one combination of covariate regime (marker fill) and context regime (color). The covariate regimes are no covariates (∅), the genotype-parameter vector (*g*), the temperature trajectory (*e*), and both (*g* + *e*). The context regimes are *no context, sparse context* (*random context* in the legend), and *max-height context* (*max context* in the legend). The vertical axis shows the squared *Sig-MMD* scaled by 10^3^. Summary markers give each model’s median and mean and its 25–75% and 5–95% ranges over the test conditions. Because the reference is the same single condition for every covariate regime, scores are comparable across covariate regimes here. Lower is better.

## J Active Context Selection

Uncertainty selection computes the standard deviation of the predicted heights across sampled trajectories at each candidate day, discards days already observed, and adds the day of highest predictive standard deviation to the context set. Random selection instead picks uniformly at random among the days not yet observed.

Figure 18 contrasts random and uncertainty-based context selection as points of a held-out trajectory are revealed one at a time. Each score measures how closely the predictions match that trajectory as context is added. Under random selection both models improve only gradually, while uncertainty sampling reaches a much lower score from the first point. The latent models therefore produce informative uncertainty estimates, and the *LNP* stays below the *ANP* throughout.

**Figure 18:**
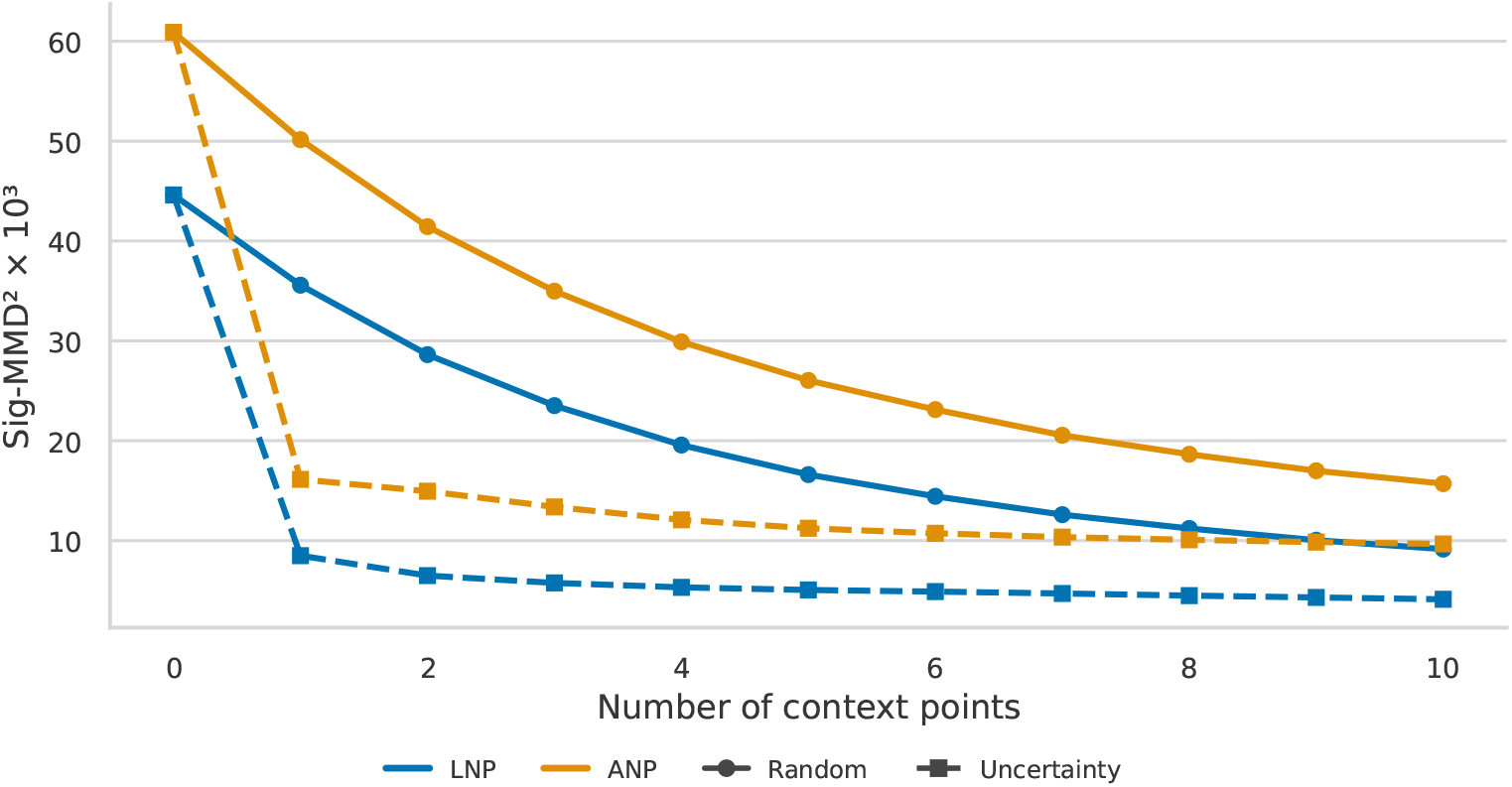
Signature maximum mean discrepancy (*Sig-MMD*) versus the number of observed context points, for the latent (*LNP*) and attentive (*ANP*) neural processes, under random context selection and uncertainty sampling, where the next point is the day of highest predictive standard deviation. For each held-out condition, points of one simulator trajectory are revealed one at a time, and the *Sig-MMD* is computed against that trajectory rather than the matched simulator distribution. Each curve is the mean over the held-out trajectories. Solid lines with circles show random selection, and dashed lines with squares uncertainty sampling. The vertical axis shows the squared *Sig-MMD* scaled by 10^3^. Predictions use no covariates (∅). Lower is better.

## K Training Length and Dataset Size

Figure 19 shows the effect of training length and dataset size. The dataset size is the number of distinct conditions, and the training length is the total number of seen trajectories. A large improvement appears when training longer than *100k* seen trajectories. At *100k*, both model families remain substantially worse and show large variation across settings. Training to *1m* accounts for most of the improvement, while additional training to *3m* or *5m* gives smaller and less uniform gains. For dataset size, even *1k* conditions already achieves reasonable scores, with a clear improvement when going to *8k*, after which only a small improvement remains. Training is very robust, and only the *1k* dataset trained for *3m* or *5m* samples shows clear signs of overfitting. All other experiments use *3m* seen trajectories and *524k* conditions.

**Figure 19:**
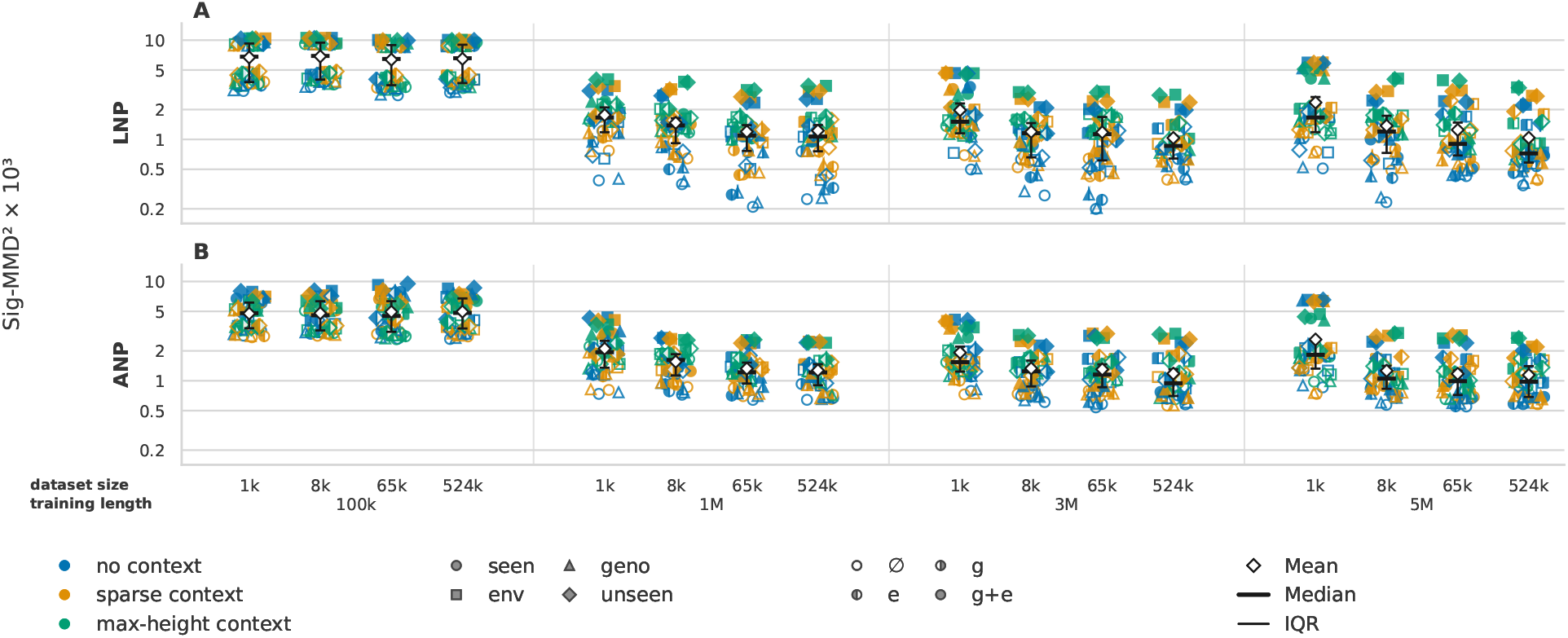
Signature maximum mean discrepancy (*Sig-MMD*) scores under the matched simulator distribution for the latent (*LNP*) and attentive (*ANP*) neural processes across dataset sizes and training lengths. The groups along the horizontal axis are the training lengths in seen trajectories, and the positions within a group the dataset sizes in distinct conditions. Each marker is the score of one combination of context regime (color), evaluation split (marker shape), and covariate regime (marker fill). The evaluation splits are seen genotypes and environments (*seen*), held-out genotypes (*geno*), held-out environments (*env*), and both held out (*unseen*). The covariate regimes are no covariates (∅), the genotype-parameter vector (*g*), the temperature trajectory (*e*), and both (*g* + *e*). Overlaid symbols give the mean, median, and interquartile range (*IQR*) over the markers of a training length and dataset size. The vertical axis shows the squared *Sig-MMD* scaled by 10^3^ on a logarithmic scale. Lower is better.

## L Latent Space Structure

Figure 20 shows the structure of the learned prior latent space along its principal components. Some components appear to capture interpretable modes of variation, though they are not fully disentangled. A few mainly modulate the maximum height, for example PC1 of the *LNP* (Figure 20A), while others affect the late drop characteristic of lodging, for example PC2 of the *ANP* (Figure 20G).

**Figure 20:**
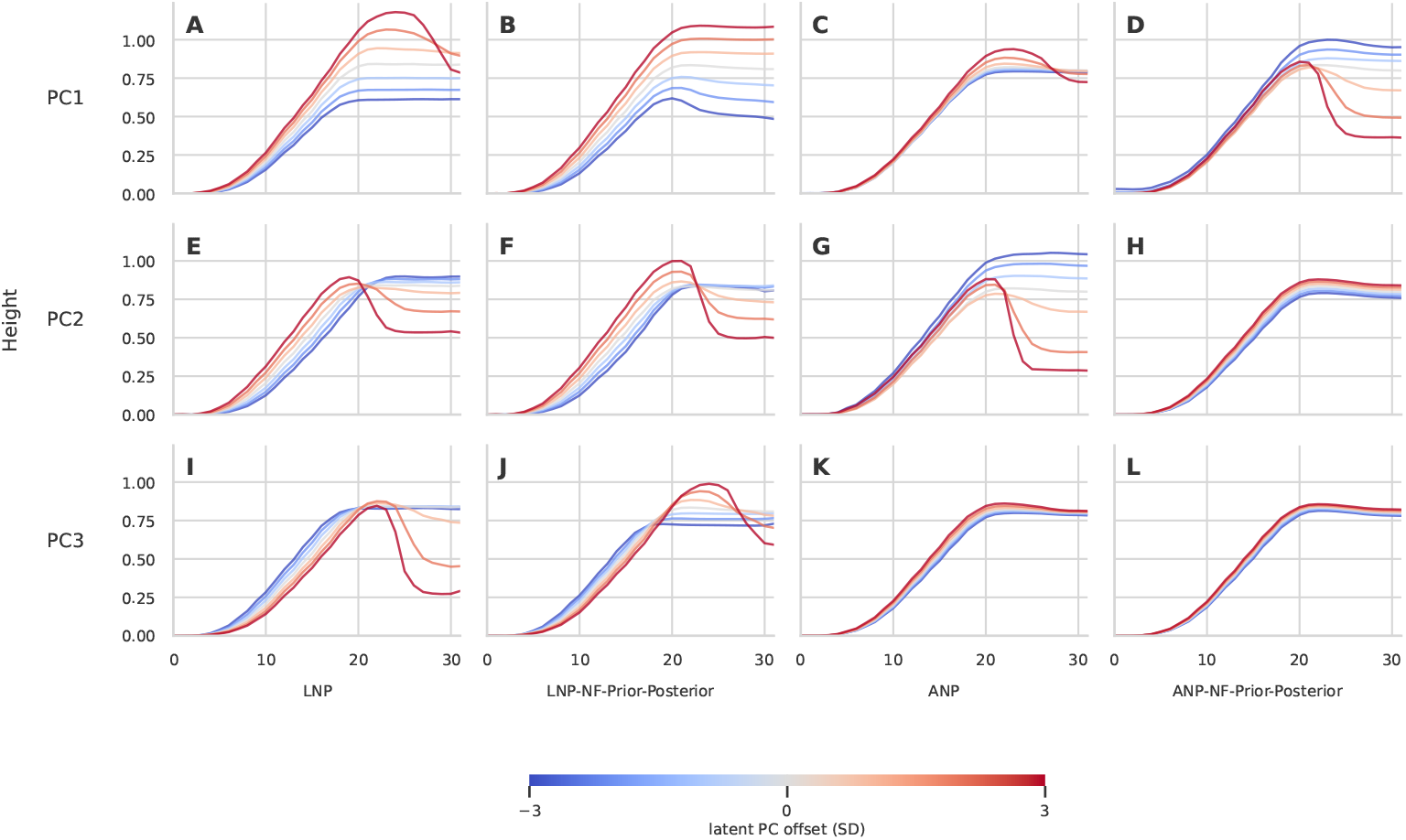
Decoded height trajectories along the principal axes of the prior latent space. The models are the latent (*LNP*) and attentive (*ANP*) neural processes, each without and with normalizing flows (*NF*) on both prior and posterior (*LNP-NF-Prior-Posterior, ANP-NF-Prior-Posterior*). For each model (columns) we fit a principal component analysis (*PCA*) to samples from the prior latent and vary each of the first three principal components (rows, PC1–PC3) from *−*3 to +3 standard deviations (*SD*), decoding every offset into a height curve. Color encodes the latent offset along the component, from *−*3 (blue) to +3 (red). The prior is conditioned on no height context and no covariates. The horizontal axis is the index of the 32 prediction time points, which span the season. Some components mainly modulate the maximum height, while others govern the late drop associated with lodging.

## M Tabulated Distributional Scores

The boxplots in Figures 12 and 13 show the full distribution of setting-level scores for each model. This section reports the same scores as numbers, averaged over the test conditions and over the five training runs, so that the model variants can be read off directly within each context and covariate regime. Table 11 gives the mean *Sig-MMD* and Table 12 the mean *CSig-MMD*. The orderings match the corresponding figures, with the latent models scoring well below the conditional *CNP* and *ACNP* and the flow variants among the best, and the variant differences are more pronounced on the tail score of Table 12.

**Table 11:** Mean squared signature maximum mean discrepancy (*Sig-MMD*^2^), scaled by 10^3^, under the matched simulator distribution and averaged over the test conditions and over five training runs, for every model across the context and covariate regimes. The models are the conditional (*CNP*), attentive conditional (*ACNP*), latent (*LNP*), and attentive (*ANP*) neural processes. Variants with a normalizing flow (*NF*) on the prior, the posterior, or both carry the suffixes *NF-Prior, NF-Posterior*, and *NF-Prior-Posterior. LNP-bigger* and *ANP-bigger* are the *LNP* and *ANP* with four instead of two decoder layers, trained once. *Baseline* is the lodging-mixture spline model with genotype and environment effects, fitted once. The column labels give the covariate regime, with no covariates (*none*), the temperature trajectory (*e*), the genotype-parameter vector (*g*), or both (*e+g*). Cells are shaded relative to the plain *LNP*, green where a model scores lower and red where it scores higher. Bold marks the lowest value in each column, together with every value within two standard errors of it. The standard error is estimated over the five runs, and the single runs of the bigger variants and the single fit of the baseline have none. Lower is better.

| Model | Mean Sig-MMD<sup>2</sup> × 10<sup>3</sup> (↓) |  |  |  |  |  |  |  |  |  |  |  |
| --- | --- | --- | --- | --- | --- | --- | --- | --- | --- | --- | --- | --- |
|  | no context |  |  |  | sparse context |  |  |  | max-height context |  |  |  |
|  | none | e | g | e+g | none | e | g | e+g | none | e | g | e+g |
| CNP | 3.071 | 3.582 | 3.952 | 4.732 | 3.258 | 3.398 | 3.867 | 4.680 | 2.812 | 2.810 | 3.763 | 4.343 |
| ACNP | 2.955 | 3.480 | 3.796 | 4.543 | 3.155 | 3.321 | 3.739 | 4.445 | 2.694 | 2.681 | 3.642 | 4.156 |
| LNP | 0.419 | 0.863 | 0.686 | 1.474 | 0.611 | 1.008 | 0.734 | 1.705 | 1.084 | 1.292 | 1.016 | 2.178 |
| LNP-NF-Prior | 0.306 | 0.677 | 0.416 | 1.085 | 0.377 | 0.795 | 0.451 | 1.280 | 0.476 | 0.726 | 0.712 | **1.757** |
| LNP-NF-Posterior | 0.390 | 0.769 | 0.622 | 1.338 | 0.567 | 0.941 | 0.686 | 1.525 | 0.548 | 0.807 | 0.816 | 1.864 |
| LNP-NF-Prior-Posterior | 0.268 | 0.624 | **0.368** | 1.093 | 0.361 | 0.751 | **0.388** | 1.263 | 0.492 | 0.758 | **0.688** | 1.834 |
| ANP | 0.732 | 1.118 | 0.865 | 1.579 | 0.865 | 1.143 | 0.904 | 1.737 | 1.264 | 1.364 | 1.437 | 2.134 |
| ANP-NF-Prior | 0.529 | 0.883 | 0.602 | 1.301 | 0.566 | 0.833 | 0.591 | 1.389 | 1.073 | 1.297 | 1.208 | **1.790** |
| ANP-NF-Posterior | 0.666 | 1.048 | 0.801 | 1.474 | 0.822 | 1.101 | 0.816 | 1.551 | 0.857 | 0.969 | 1.159 | **1.685** |
| ANP-NF-Prior-Posterior | 0.533 | 0.898 | 0.529 | 1.285 | 0.579 | 0.844 | 0.540 | 1.371 | 0.651 | 0.897 | 0.698 | **1.404** |
| LNP-bigger | 0.466 | 0.999 | 0.694 | 1.740 | 0.637 | 1.184 | 0.778 | 1.965 | 0.712 | 1.040 | 0.908 | 2.382 |
| ANP-bigger | 0.826 | 1.212 | 0.831 | 1.667 | 0.841 | 1.098 | 0.930 | 1.787 | 0.609 | 0.798 | 0.939 | 1.713 |
| Baseline | **0.127** | **0.164** | 0.391 | **0.609** | **0.108** | **0.146** | **0.367** | **0.720** | **0.024** | **0.124** | **0.460** | **1.548** |

**Table 12:** Mean squared censored signature maximum mean discrepancy (*CSig-MMD*^2^), scaled by 10^3^, under the matched simulator distribution and averaged over the test conditions and over five training runs, for every model across the context and covariate regimes. The models are the conditional (*CNP*), attentive conditional (*ACNP*), latent (*LNP*), and attentive (*ANP*) neural processes. Variants with a normalizing flow (*NF*) on the prior, the posterior, or both carry the suffixes *NF-Prior, NF-Posterior*, and *NF-Prior-Posterior. Baseline* is the lodging-mixture spline model with genotype and environment effects, fitted once. The column labels give the covariate regime, with no covariates (*none*), the temperature trajectory (*e*), the genotype-parameter vector (*g*), or both (*e+g*). Shading and bold follow the same conventions as Table 11. Lower is better.

| Model | Mean $CSig-MMD^2 \times 10^3$ (↓) | | | | | | | | | | | |
| --- | --- | --- | --- | --- | --- | --- | --- | --- | --- | --- | --- | --- |
|  | no context |  |  |  | sparse context |  |  |  | max-height context |  |  |  |
|  | none | e | g | e+g | none | e | g | e+g | none | e | g | e+g |
| CNP | 11.555 | 12.495 | 40.149 | 44.326 | 4.024 | 4.986 | 38.680 | 44.230 | 3.329 | 3.759 | 34.768 | 42.754 |
| ACNP | 11.555 | 12.481 | 41.925 | 42.194 | 3.087 | 4.230 | 39.047 | 42.686 | 2.430 | 2.893 | 34.213 | 42.254 |
| LNP | 0.612 | 1.339 | 1.529 | 2.873 | 0.736 | 1.955 | 1.558 | 3.418 | 1.074 | 2.505 | 1.768 | 4.599 |
| LNP-NF-Prior | 0.624 | 1.188 | **0.548** | 1.419 | 0.982 | 2.020 | **0.654** | 1.887 | 0.683 | 1.648 | **0.986** | **3.019** |
| LNP-NF-Posterior | 0.701 | 1.275 | 1.267 | 2.309 | 0.616 | 1.556 | 1.261 | 2.757 | 0.921 | 1.732 | 1.939 | 4.229 |
| LNP-NF-Prior-Posterior | 0.908 | 1.498 | **0.613** | 1.609 | 1.407 | 2.410 | **0.733** | 2.096 | 1.109 | 2.321 | **1.051** | 3.371 |
| ANP | 0.760 | 1.351 | 1.884 | 3.059 | 0.572 | 1.413 | 2.138 | 3.631 | 0.777 | 0.923 | 2.788 | 4.075 |
| ANP-NF-Prior | 1.415 | 1.903 | 1.235 | 2.160 | 1.150 | 1.892 | 1.301 | 2.525 | 0.858 | **1.149** | 2.771 | 3.720 |
| ANP-NF-Posterior | 0.560 | 1.159 | 1.177 | 2.310 | 0.514 | 1.351 | 1.615 | 2.722 | 1.169 | 1.264 | 4.190 | **4.619** |
| ANP-NF-Prior-Posterior | 1.791 | 2.314 | 1.582 | 2.749 | 1.392 | 1.957 | 1.746 | 3.147 | 0.699 | 0.939 | 2.393 | 4.031 |
| Baseline | **0.047** | **0.297** | **0.578** | **1.154** | **0.240** | **0.451** | **0.713** | **1.446** | **0.123** | **0.526** | **0.963** | **2.834** |

## N Lodging-Mixture Spline Baseline

The baseline is the lodging-mixture spline model of eq. (15), fitted to the same training data as the neural processes. In this form, the covariates *u* enter only as the identity of a genotype and of a year-site, so the model can neither predict for an unseen genotype or year-site nor use observed heights. The regression of effects and the importance sampling of Section 7.2 provide both. The model as stated, conditioned on context by the same importance sampling, is reported as an ablation.

### N.1 Growth Model

Each trajectory is a cubic P-spline in calendar day on days 200 to 322 with six uniform interior knots and ten coefficients, and the curve is constant outside this window. The coefficients are fitted by penalized least squares on the observation days before the detected lodging day, with a first-difference penalty of weight 0.01 that keeps the curve flat where observations are missing. They are decomposed into a mean, a genotype effect, a year-site effect, and a Gaussian residual, where each effect is the mean deviation over its trajectories. The residual covariance is estimated from the trajectories without a detected lodging drop, after removing the estimation noise of the single fit per condition.

### N.2 Lodging

Lodging is detected from the heights, without the labels of the simulator. A day is a lodging day if the mean of the seven observations before it exceeds the mean of all observations from that day on by more than 0.1 m, with at least five later observations. The lodging probability π is a logistic function of the peak height, the maximum of the three-day running mean of the observed heights. Each detected lodging event yields a triple of the delay from the end of growth to the lodging day, the severity as the ratio of final to peak height, and the duration of the drop. The pool of triples is corrected for the bias of the detection by re-detecting trajectories sampled from the fitted model.

### N.3 Sampling

A draw adds the genotype effect, the year-site effect, and a residual to the mean coefficients and evaluates the growth curve. With the probability π of its peak height, it takes a triple from the pool and falls linearly from the height at the lodging day to the severity times the peak height. Each condition receives 64 draws, as for the neural processes. If the covariate regime does not include the genotype or the year-site, its effect is drawn at random from the training effects, as is the effect of an unseen genotype or year-site in the model as stated. With the regression of effects, the effect of an unseen genotype or year-site is instead predicted by ridge regression on the genotype covariate, as in genomic best linear unbiased prediction, or on the cumulative thermal time of the season, with the ridge weight chosen by leave-one-out prediction of the training effects. Each draw adds a Gaussian prediction error with the covariance of the leave-one-out errors. With context, a pool of 16,384 prior draws per condition is weighted by eq. (8), with the noise scale estimated from the training data, and 64 draws are resampled. In the max-height regime, the effective number of draws falls below 10 for up to a quarter of the conditions without covariates on unseen year-sites.

### N.4 Advantage from the Evaluation

On the synthetic data, the baseline profits from the evaluation itself. The test conditions come from the same generative distributions as the training conditions, and the matched simulator distribution is the simulator marginal over the conditions consistent with the covariates and the context. A retrieval model that returns the training trajectories consistent with the covariates, resampled with the context weights, reproduces this distribution by construction. The baseline is a smoothed retrieval model wherever it looks up a fitted effect rather than predicting one, that is on seen genotypes and year-sites and in every split without covariates.

### N.5 Scores

The baseline is scored with the blocked context-matched *Sig-MMD* and *CSig-MMD* in every setting of the neural processes, and the score figures of the main text include it in every context regime. Table 13 compares the model as stated with the regression of effects.

**Table 13:** Mean squared signature maximum mean discrepancy (*Sig-MMD*^2^), scaled by 10^3^, of the baseline as stated and with regression of effects, in the settings with an unseen genotype or year-site, next to the reported run of the latent neural process (*LNP*) with a normalizing flow on the prior and the posterior (*NF-Prior-Posterior*). The baseline as stated draws the missing effect from the training effects, and the baseline with regression predicts the genotype effect from the genotype-parameter vector and the year-site effect from the thermal time. With context, both are conditioned by importance sampling. The evaluation splits are held-out genotypes (*geno*), held-out environments (*env*), and both held out (*unseen*), and the covariate regimes are temperature (*e*), genotype-parameter vector (*g*), and both (*e+g*). Lower is better.

| Context regime | Model | geno/g | geno/e+g | env/e | env/e+g | unseen/e | unseen/g | unseen/e+g |
| --- | --- | --- | --- | --- | --- | --- | --- | --- |
| No context | LNP-NF-Prior-Posterior | 0.24 | 0.32 | 0.97 | 1.81 | 1.01 | 0.47 | 1.76 |
| No context | Baseline as stated | 5.73 | 6.11 | 0.85 | 1.75 | 0.90 | 5.65 | 6.85 |
| No context | Baseline with regression | 0.27 | 0.45 | 0.22 | 0.76 | 0.26 | 0.57 | 0.80 |
| Sparse context | LNP-NF-Prior-Posterior | 0.27 | 0.34 | 1.17 | 2.16 | 1.26 | 0.52 | 2.11 |
| Sparse context | Baseline as stated | 2.55 | 3.61 | 0.57 | 1.72 | 0.62 | 2.19 | 4.56 |
| Sparse context | Baseline with regression | 0.25 | 0.48 | 0.22 | 0.95 | 0.23 | 0.54 | 1.01 |

**Table 14:** Mean and standard deviation (*SD*) over five training runs of the averaged squared signature maximum mean discrepancy (*Sig-MMD*^2^) and its censored variant (*CSig-MMD*^2^), scaled by 10^3^, under the matched simulator distribution. Each value averages the 16 combinations of evaluation split and covariate regime within a context regime. The models are the conditional (*CNP*), attentive conditional (*ACNP*), latent (*LNP*), and attentive (*ANP*) neural processes. Variants with a normalizing flow (*NF*) on the prior, the posterior, or both carry the suffixes *NF-Prior, NF-Posterior*, and *NF-Prior-Posterior*. Bold marks the lowest mean in each column. Lower is better.

| Model | $Sig\text{-}MMD^2 \times 10^3$ (↓) | | | $CSig\text{-}MMD^2 \times 10^3$ (↓) | | |
| --- | --- | --- | --- | --- | --- | --- |
|  | no context | sparse | max-height | no context | sparse | max-height |
| CNP | $3.83 \pm 0.19$ | $3.80 \pm 0.16$ | $3.43 \pm 0.12$ | $27.13 \pm 3.18$ | $22.98 \pm 2.66$ | $21.15 \pm 5.34$ |
| ACNP | $3.69 \pm 0.07$ | $3.67 \pm 0.04$ | $3.29 \pm 0.03$ | $27.04 \pm 0.81$ | $22.26 \pm 0.64$ | $20.45 \pm 1.41$ |
| LNP | $0.86 \pm 0.08$ | $1.01 \pm 0.07$ | $1.39 \pm 0.15$ | $1.59 \pm 0.41$ | $1.92 \pm 0.28$ | $2.49 \pm 0.18$ |
| LNP-NF-Prior | $0.62 \pm 0.03$ | $0.73 \pm 0.07$ | $0.92 \pm 0.28$ | **$0.94 \pm 0.09$** | **$1.39 \pm 0.17$** | **$1.58 \pm 0.39$** |
| LNP-NF-Posterior | $0.78 \pm 0.04$ | $0.93 \pm 0.06$ | $1.01 \pm 0.11$ | $1.39 \pm 0.40$ | $1.55 \pm 0.42$ | $2.21 \pm 0.74$ |
| LNP-NF-Prior-Posterior | **$0.59 \pm 0.03$** | **$0.69 \pm 0.04$** | $0.94 \pm 0.13$ | $1.16 \pm 0.14$ | $1.66 \pm 0.11$ | $1.96 \pm 0.23$ |
| ANP | $1.07 \pm 0.03$ | $1.16 \pm 0.03$ | $1.55 \pm 0.19$ | $1.76 \pm 0.23$ | $1.94 \pm 0.23$ | $2.14 \pm 0.62$ |
| ANP-NF-Prior | $0.83 \pm 0.06$ | $0.84 \pm 0.04$ | $1.34 \pm 0.67$ | $1.68 \pm 0.08$ | $1.72 \pm 0.07$ | $2.12 \pm 0.32$ |
| ANP-NF-Posterior | $1.00 \pm 0.06$ | $1.07 \pm 0.08$ | $1.17 \pm 0.16$ | $1.30 \pm 0.32$ | $1.55 \pm 0.21$ | $2.81 \pm 1.64$ |
| ANP-NF-Prior-Posterior | $0.81 \pm 0.06$ | $0.83 \pm 0.06$ | **$0.91 \pm 0.16$** | $2.11 \pm 0.29$ | $2.06 \pm 0.33$ | $2.02 \pm 0.71$ |

## O Variance Across Training Runs

Each model was trained five times with different randomness and identical training data, architecture, and training budget, and every run was scored as in the main results. The score figures of the main text and the tables of Appendix M summarize all five runs, and the other figures and tables of the synthetic study show the reported run, one of the five, except for the retrained models of Appendix P. For each model and context regime, Table 14 gives the mean and *SD* over the runs of the score averaged over the settings of the regime. Table 15 tests pairs of models with a Welch t-test over the runs, per setting with a Holm correction over the settings of a regime, and on the averaged score.

**Table 15:**
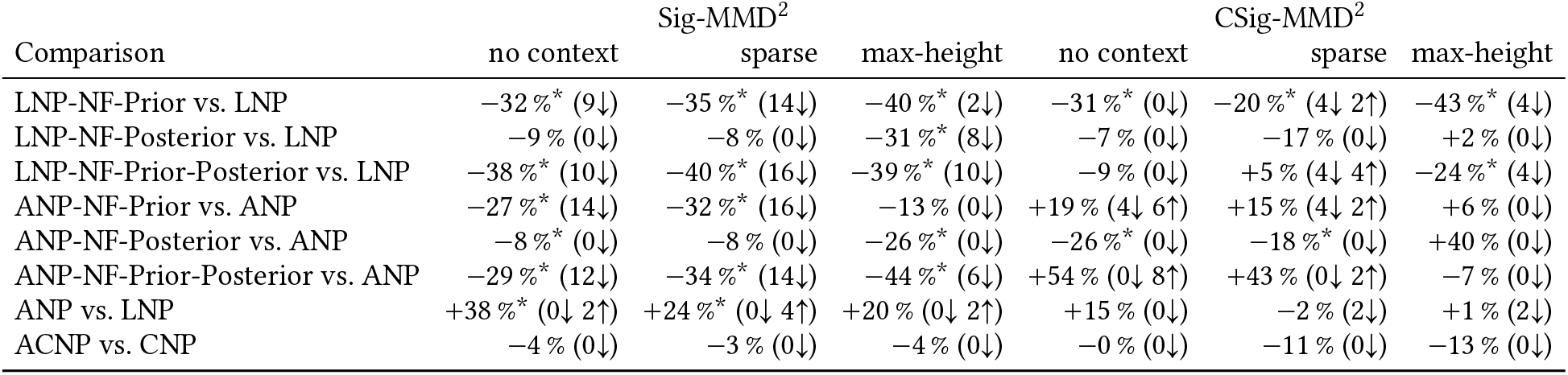
Differences between models over five training runs on the squared signature maximum mean discrepancy (*Sig-MMD*^2^) and its censored variant (*CSig-MMD*^2^). Each entry gives the mean relative difference of the first model to the second over the 16 combinations of evaluation split and covariate regime within a context regime, so that a negative value means that the first model scores lower. An asterisk marks a significant Welch t-test over the five runs on the averaged score (*p* < 0.05). The brackets give the number of settings in which the first model is significantly lower (↓) or higher (↑) than the second, by a Welch t-test over the runs with a Holm correction over these 16 combinations (family-wise level 0.05). The models are the conditional (*CNP*), attentive conditional (*ACNP*), latent (*LNP*), and attentive (*ANP*) neural processes. Variants with a normalizing flow (*NF*) on the prior, the posterior, or both carry the suffixes *NF-Prior, NF-Posterior*, and *NF-Prior-Posterior*.

## P Sensitivity to Simulator Settings

Two simulator settings were varied, the lodging rate and the observation noise. For each changed setting, the models were trained once on training data with that setting and evaluated on evaluation splits with the same setting, with the evaluation splits, covariate regimes, context regimes, and blocking of the main results. The *CSig-MMD* reference and its threshold were fitted on the training data of each setting with the rule of Appendix E. Scores are therefore not comparable across settings, and the tables rank the models within each setting. Each of the 12 combinations of context regime and covariate regime, averaged over the four evaluation splits, ranks the models from 1 (lowest score) upwards, and the tables give the mean rank over these combinations.

The lodging rates of 7.0 % and 2.7 % follow from Weibull scales of λ = 1.3 m and 1.5 m in the lodging model of Appendix H, instead of 1.1 m for the default rate of 19.7 %. All ten models were trained at each rate. At 2.7 %, 65 % of the test conditions keep at least one simulator draw above the tail threshold, compared with 98 % at 19.7 %. Table 16 gives the mean ranks under both metrics and Figure 21 the lodging rate of every model against maximum height. The observation noise was lowered from the default *SD* of 0.05 m to 0.02 m, the noise measured on *FIP1*, for the *LNP*, the *ANP*, and their *NF-Prior-Posterior* variants. Table 17 gives their mean ranks at both noise levels.

**Table 16:**
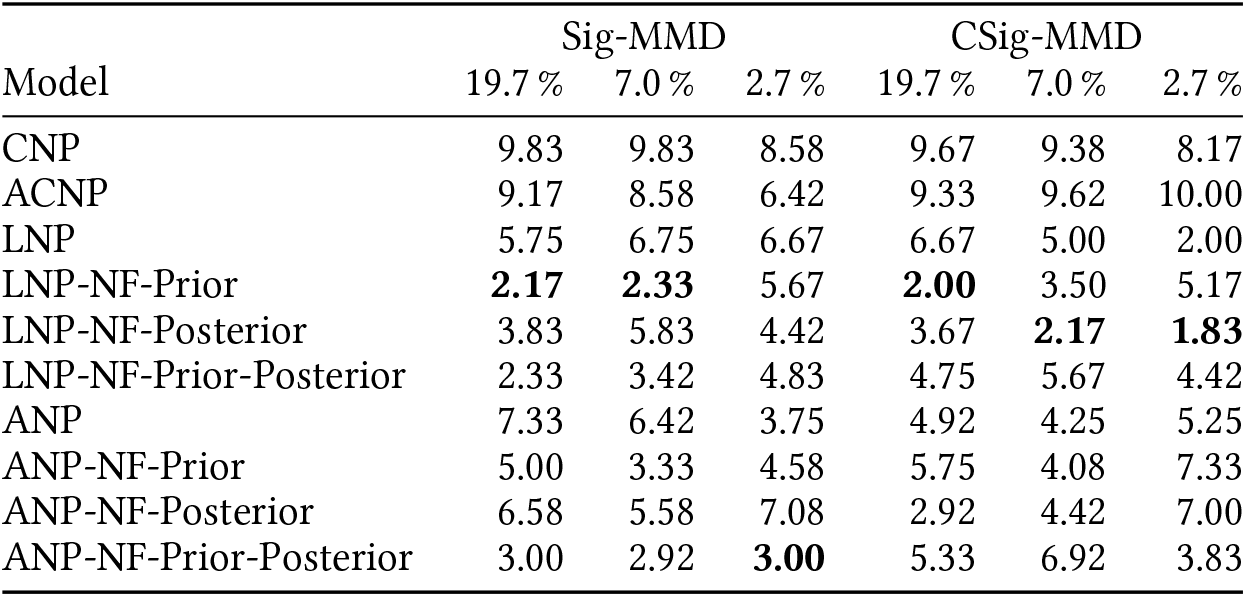
Mean rank of the models at three lodging rates under the signature maximum mean discrepancy (*Sig-MMD*) and its censored variant (*CSig-MMD*), with the models trained and evaluated at each rate. Each of the 12 combinations of context regime and covariate regime, averaged over the four evaluation splits, ranks the models from 1 (lowest score) to 10, and the mean over these combinations is given. The models are the conditional (*CNP*), attentive conditional (*ACNP*), latent (*LNP*), and attentive (*ANP*) neural processes. Variants with a normalizing flow (*NF*) on the prior, the posterior, or both carry the suffixes *NF-Prior, NF-Posterior*, and *NF-Prior-Posterior*. Bold marks the best mean rank in each column.

| Model | Sig-MMD |  |  | CSig-MMD |  |  |
| --- | --- | --- | --- | --- | --- | --- |
|  | 19.7 % | 7.0 % | 2.7 % | 19.7 % | 7.0 % | 2.7 % |
| CNP | 9.83 | 9.83 | 8.58 | 9.67 | 9.38 | 8.17 |
| ACNP | 9.17 | 8.58 | 6.42 | 9.33 | 9.62 | 10.00 |
| LNP | 5.75 | 6.75 | 6.67 | 6.67 | 5.00 | 2.00 |
| LNP-NF-Prior | **2.17** | **2.33** | 5.67 | **2.00** | 3.50 | 5.17 |
| LNP-NF-Posterior | 3.83 | 5.83 | 4.42 | 3.67 | **2.17** | **1.83** |
| LNP-NF-Prior-Posterior | 2.33 | 3.42 | 4.83 | 4.75 | 5.67 | 4.42 |
| ANP | 7.33 | 6.42 | 3.75 | 4.92 | 4.25 | 5.25 |
| ANP-NF-Prior | 5.00 | 3.33 | 4.58 | 5.75 | 4.08 | 7.33 |
| ANP-NF-Posterior | 6.58 | 5.58 | 7.08 | 2.92 | 4.42 | 7.00 |
| ANP-NF-Prior-Posterior | 3.00 | 2.92 | **3.00** | 5.33 | 6.92 | 3.83 |

**Table 17:** Mean rank of four models at two observation-noise levels under the signature maximum mean discrepancy (*Sig-MMD*) and its censored variant (*CSig-MMD*), with the models trained and evaluated at each noise level. Each of the 12 combinations of context regime and covariate regime, averaged over the four evaluation splits, ranks the models from 1 (lowest score) to 4, and the mean over these combinations is given. The models are the latent (*LNP*) and attentive (*ANP*) neural processes and their variants with a normalizing flow (*NF*) on the prior and the posterior (*NF-Prior-Posterior*). Bold marks the best mean rank in each column.

| Model | Sig-MMD |  | CSig-MMD |  |
| --- | --- | --- | --- | --- |
|  | 0.05 m | 0.02 m | 0.05 m | 0.02 m |
| LNP | 2.92 | 3.17 | 3.08 | 2.58 |
| LNP-NF-Prior-Posterior | **1.58** | **1.33** | **2.25** | **1.92** |
| ANP | 3.83 | 3.75 | **2.25** | 3.25 |
| ANP-NF-Prior-Posterior | 1.67 | 1.75 | 2.42 | 2.25 |

**Table 18:**
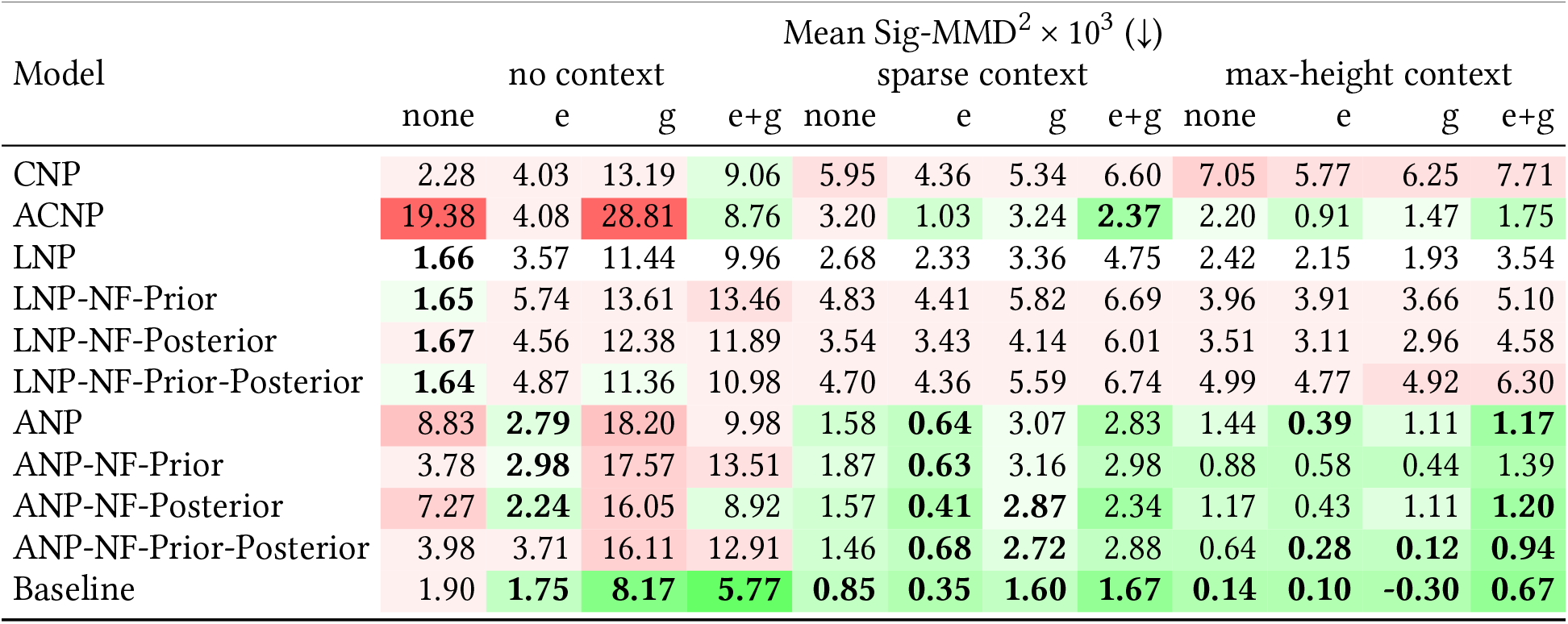
Mean squared signature maximum mean discrepancy (*Sig-MMD*^2^), scaled by 10^3^, on the Field Phenotyping Platform 1.0 (*FIP1*) dataset after pre-training on the simulator, averaged over the four evaluation splits and, for the neural processes, over five training runs, for every model across the context and covariate regimes. The models are the conditional (*CNP*), attentive conditional (*ACNP*), latent (*LNP*), and attentive (*ANP*) neural processes. Variants with a normalizing flow (*NF*) on the prior, the posterior, or both carry the suffixes *NF-Prior, NF-Posterior*, and *NF-Prior-Posterior. Baseline* is the lodging-mixture spline model with genotype and environment effects, fitted once. The column labels give the covariate regime, with no covariates (*none*), the temperature record (*e*), the markers (*g*), or both (*e+g*). Cells are shaded relative to the plain *LNP*, green where a model scores lower and red where it scores higher. Bold marks the lowest value in each column, together with every value within two standard errors of it. A perfect model scores the floor of its evaluation split, between *−*0.49 and *−*0.04, so a value below the floor counts as the floor. Lower is better.

**Table 19:**
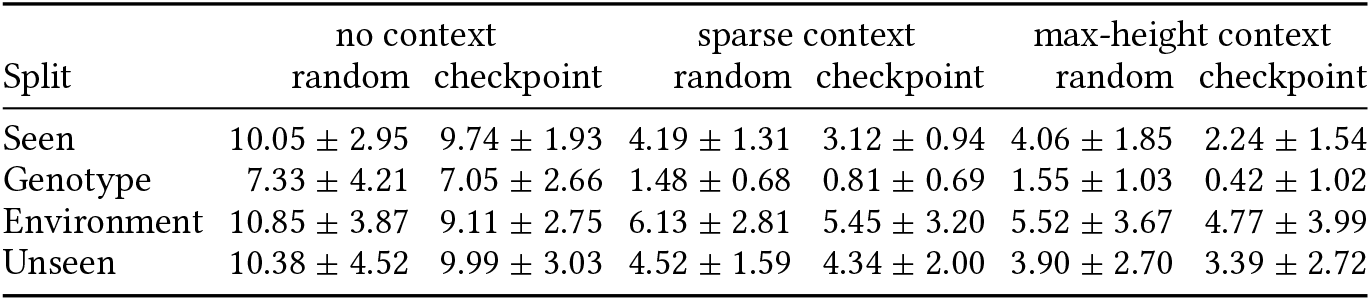
Mean squared signature maximum mean discrepancy (*Sig-MMD*^2^), scaled by 10^3^, on the Field Phenotyping Platform 1.0 (*FIP1*) dataset by evaluation split, context regime, and initialization. Each value is the mean over the ten neural processes and the four covariate regimes, with the standard deviation (*SD*) over the neural processes. *Random* is training from random initialization and *checkpoint* training from the checkpoints of the synthetic study. Lower is better.

| Split | no context |  | sparse context |  | max-height context |  |
| --- | --- | --- | --- | --- | --- | --- |
|  | random | checkpoint | random | checkpoint | random | checkpoint |
| Seen | $10.05 \pm 2.95$ | $9.74 \pm 1.93$ | $4.19 \pm 1.31$ | $3.12 \pm 0.94$ | $4.06 \pm 1.85$ | $2.24 \pm 1.54$ |
| Genotype | $7.33 \pm 4.21$ | $7.05 \pm 2.66$ | $1.48 \pm 0.68$ | $0.81 \pm 0.69$ | $1.55 \pm 1.03$ | $0.42 \pm 1.02$ |
| Environment | $10.85 \pm 3.87$ | $9.11 \pm 2.75$ | $6.13 \pm 2.81$ | $5.45 \pm 3.20$ | $5.52 \pm 3.67$ | $4.77 \pm 3.99$ |
| Unseen | $10.38 \pm 4.52$ | $9.99 \pm 3.03$ | $4.52 \pm 1.59$ | $4.34 \pm 2.00$ | $3.90 \pm 2.70$ | $3.39 \pm 2.72$ |

**Figure 21:**
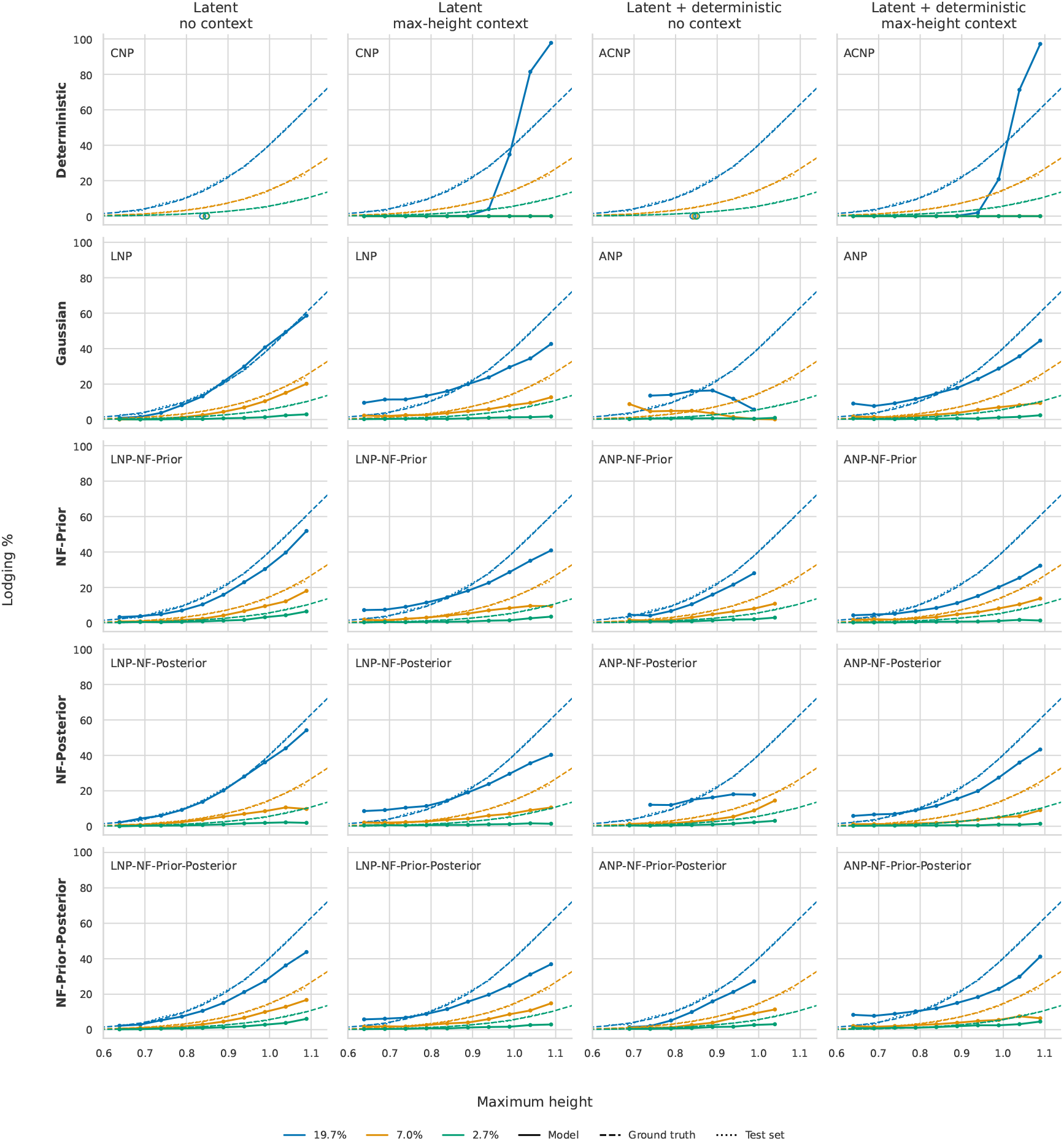
Lodging rate over maximum height at lodging rates of 19.7 %, 7.0 %, and 2.7 %, with each model trained and evaluated at the same rate. Solid curves are the models, dashed curves the analytical lodging probability of the synthetic generator (*Ground truth*), and dotted curves the lodging rate of the evaluation data (*Test set*). Predicted trajectories are counted as lodged when their height falls by at least 20 % and at least 10 cm below their maximum. The *Latent* columns contain the global models, the conditional (*CNP*) and latent (*LNP*) neural processes and the *LNP* variants with normalizing flows (*NF*), and the *Latent + deterministic* columns the per-target models, the attentive conditional (*ACNP*) and attentive (*ANP*) neural processes and the *ANP* variants. Within each column pair, the rows are the conditional model (*Deterministic*), the latent model (*Gaussian*), and its flow variants with a flow on the prior (*NF-Prior*), the posterior (*NF-Posterior*), or both (*NF-Prior-Posterior*). In the *no context* columns models receive no height points, and in the *max-height context* columns all noisy context points up to the maximum height of the trajectory. Without context, plants are grouped by their predicted maximum height, and with context by their true maximum height. Bins with fewer than 1 % of the plants are not shown. Without context, the *CNP* and *ACNP* predict one trajectory for all plants, shown as one point at its maximum height. All predictions use no covariates (∅).

**Figure 22:**
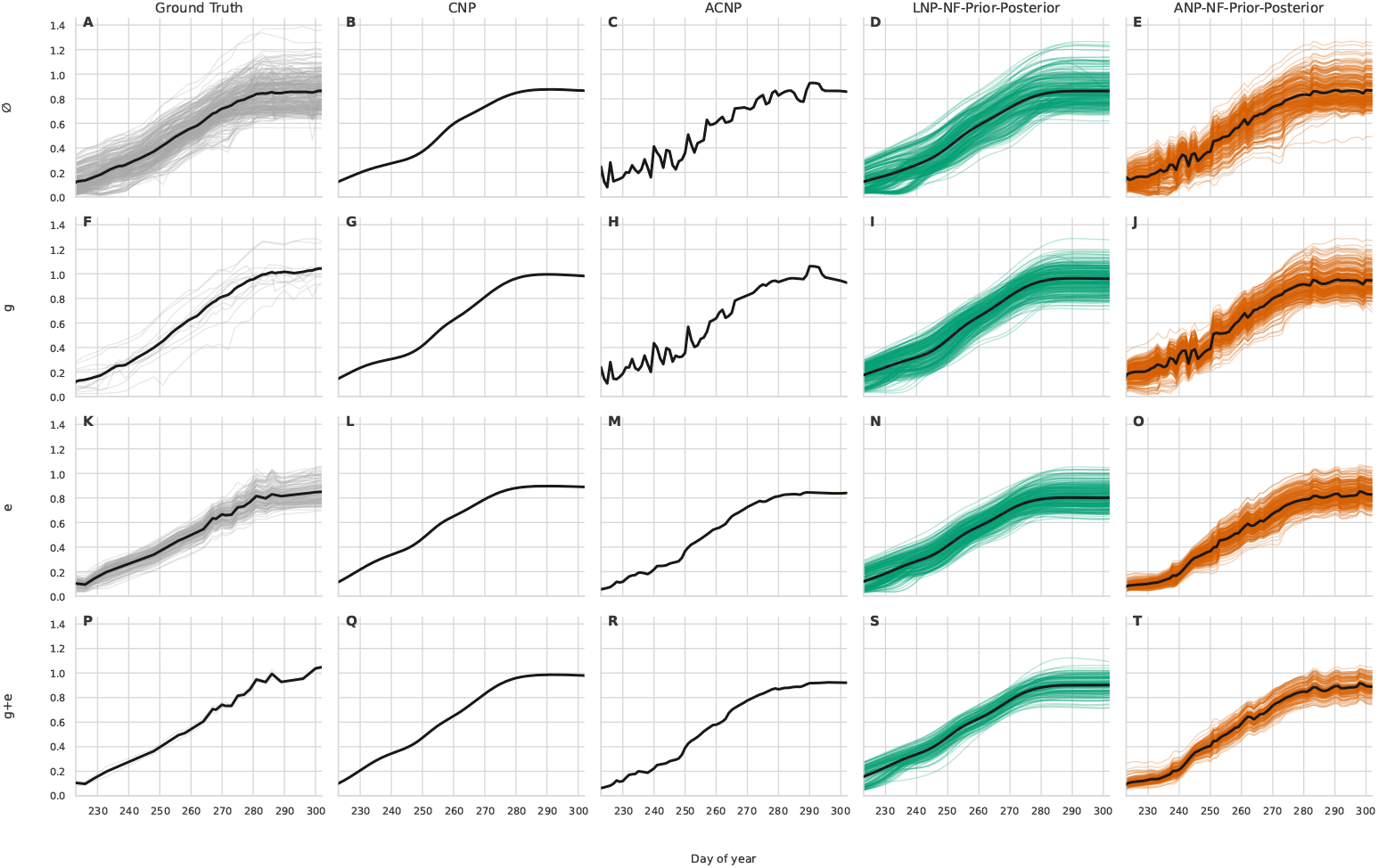
Height trajectory samples on the Field Phenotyping Platform 1.0 (*FIP1*) dataset under each covariate regime, without context. Columns show the measured plots (*Ground Truth*), the conditional (*CNP*) and attentive conditional (*ACNP*) neural processes, and the latent (*LNP*) and attentive (*ANP*) neural processes with normalizing flows (*NF*) on both prior and posterior (*LNP-NF-Prior-Posterior, ANP-NF-Prior-Posterior*), after pre-training on the simulator. Rows show the covariate regimes with no covariates (∅), the markers (*g*), the temperature record (*e*), and both (*g* + *e*), and the row label gives the covariates passed to the model. The ∅ row draws from all four evaluation splits, the *g* row shows one genotype across its year-sites, the *e* row one year-site, and the *g* + *e* row their common plots. Thin lines are individual samples, and thick lines are their means. Days are counted from the first of September. Since the *CNP* and *ACNP* have no latent variable, they produce a single predictive mean trajectory for each covariate regime.

## Q Application to FIP1

### Q.1 Data and Training

*FIP1* provides 2,930 training and 380 validation plots from five harvest years, with heights on 20 to 48 irregular dates per plot, temperature records, and 18,845 markers per genotype. The four evaluation splits hold 250 plots of seen genotypes and years (*seen*), 246 plots of held-out genotypes (*geno*), 190 plots of seen genotypes in the held-out year 2019 (*env*), and 62 plots of held-out genotypes in that year (*unseen*). The 2019 plots come from two trials with one temperature record and form one environment. All ten configurations were trained with five seeds and one million training samples each, once from random initialization and once from the checkpoints of the synthetic study, which initialize all weights except the marker encoder, whose input dimension differs. Markers are available for 261 of the 849 training genotypes, and a genotype without markers enters the neural processes without a genotype token, as in the covariate regimes without *g*.

### Q.2 Evaluation

The evaluation uses the context regimes, the covariate regimes, and the *Sig-MMD* of the synthetic study. Since *FIP1* has no simulator, the reference of an evaluation split consists of its observed plots, each weighted with the weight of eq. (8) evaluated on its own trajectory, and all evaluation splits are placed on the shared measurement grid of Appendix B. A covariate unit with fewer draws than its blocks require fills them in repeated reshuffled passes, so that no block contains a draw twice, and a unit whose draws fit into one block is scored once. Scoring one half of the observed plots of an evaluation split against the other half gives the score of a perfect model, the floor of the split, which lies between *−*0.49 ± 0.29 and *−*0.04 ± 0.06 and sets the resolution of the metric at about 0.4. *CSig-MMD* is not reported, since its tail is defined by the lodging events of the simulator, which *FIP1* does not provide.

### Q.3 Baseline

The baseline of Section 7.2 was fitted with the same settings to the training plots. The spline of a plot is fitted on its own measurement dates, and repeated measurements on one date are averaged. Since not every genotype occurs in every year-site, the genotype and year-site effects are fitted by backfitting, which alternates the two group means until they converge. The ridge regression on the markers is fitted on the 261 training genotypes with markers, and a held-out genotype without markers receives a random training effect. The thermal-time regression is fitted on the five training year-sites. The noise scale is 0.019 m. The drop detector of the baseline, which requires at least five observations after the drop, finds 12 lodged training plots (0.4 %), fewer than the 1.6 % of lodged *FIP1* plots (Appendix A). Four of them are still observed 25 days after the drop and provide the lodging triples.

### Q.4 Scores

Figure 22 shows predictions without context next to the measured plots, Table 18 gives the scores after pre-training as the mean over the four evaluation splits, and Figure 23 shows them per split. In the table, two values differ when their difference exceeds twice its standard error, estimated over the five runs for a neural process and over the covariate units of a setting for the baseline. A setting with a single unit, whose blocks share the observed plots, takes the *SD* of the floor of its split instead. A value below the floor of its split counts as the floor. Table 19 compares the two initializations per evaluation split and context regime as the mean over the neural processes and the covariate regimes.

**Figure 23:**
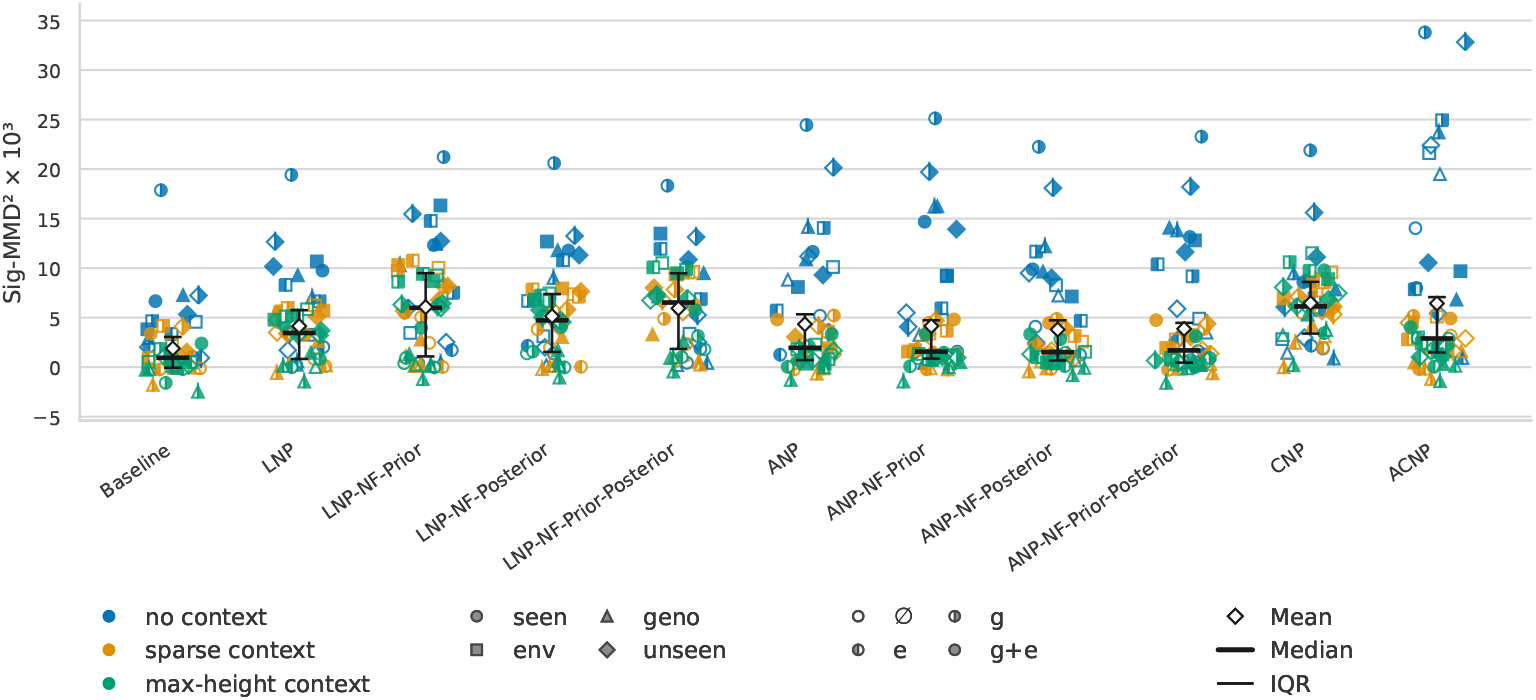
Signature maximum mean discrepancy (*Sig-MMD*) scores on the Field Phenotyping Platform 1.0 (*FIP1*) dataset after pre-training on the simulator, across model variants, evaluation splits, covariate regimes, and context regimes. The models are the conditional (*CNP*), attentive conditional (*ACNP*), latent (*LNP*), and attentive (*ANP*) neural processes. Variants with a normalizing flow (*NF*) on the prior, the posterior, or both carry the suffixes *NF-Prior, NF-Posterior*, and *NF-Prior-Posterior. Baseline* is the lodging-mixture spline model with genotype and environment effects, shown as a single fit. The evaluation splits are seen genotypes and years (*seen*), held-out genotypes (*geno*), the held-out year (*env*), and held-out genotypes in the held-out year (*unseen*). Each marker is the score of one combination of context regime (color), evaluation split (marker shape), and covariate regime (marker fill), as the mean over five training runs. The covariate regimes are no covariates (∅), the markers (*g*), the temperature record (*e*), and both (*g* + *e*). The diamond is the mean over all settings of a model, the thick bar the median, and the thin bar the interquartile range (*IQR*). The vertical axis shows the squared *Sig-MMD* scaled by 10^3^. A perfect model scores the floor of its split, between *−*0.49 and *−*0.04, so some scores are below zero. Lower is better.

## R Failure Modes of the Latent and Attentive Neural Processes

The *LNP* and the *ANP* fail in different ways. Figure 24 shows one condition of the Environment split with max-height context. Without covariates, the *LNP* draws follow the context points (Figure 24A). With both covariates, they rise earlier than the context points (Figure 24B), whereas the *ANP* follows them (Figure 24C).

**Figure 24:**
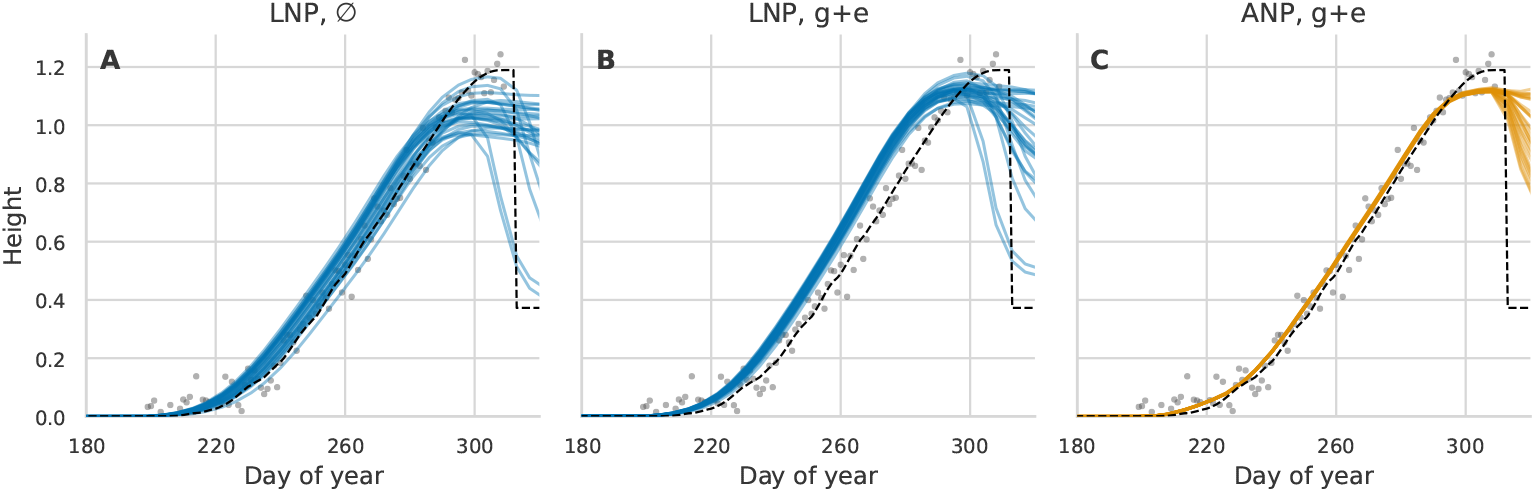
Prediction draws for one condition of the Environment split of the synthetic data, a held-out year-site with a seen genotype, with max-height context. The panels show the latent neural process (*LNP*) without covariates (∅, A) and with the genotype-parameter vector and the temperature trajectory (*g* + *e*, B), and the attentive neural process (*ANP*) with *g* + *e* (C). Gray dots are the context points, noisy observations of the trajectory up to its maximum height. The dashed line is the noise-free simulator trajectory, which lodges after its maximum. Thin lines are 28 prediction draws. Heights are in meters, and days are counted from the first of September.

Without context, the *ANP* fails instead. On *FIP1*, the mean prediction of the per-target models jumps from day to day and follows the per-day mean of the measured plots (Figure 25A), whereas that of the global models is smooth (Figure 25B). The per-day mean jumps because the siteyears measured on a day change from day to day. On synthetic data, the lodging rate of the *ANP* falls as the predicted maximum height increases beyond about 0.9 m, whereas the *LNP* follows the simulator (Figure 25C).

**Figure 25:**
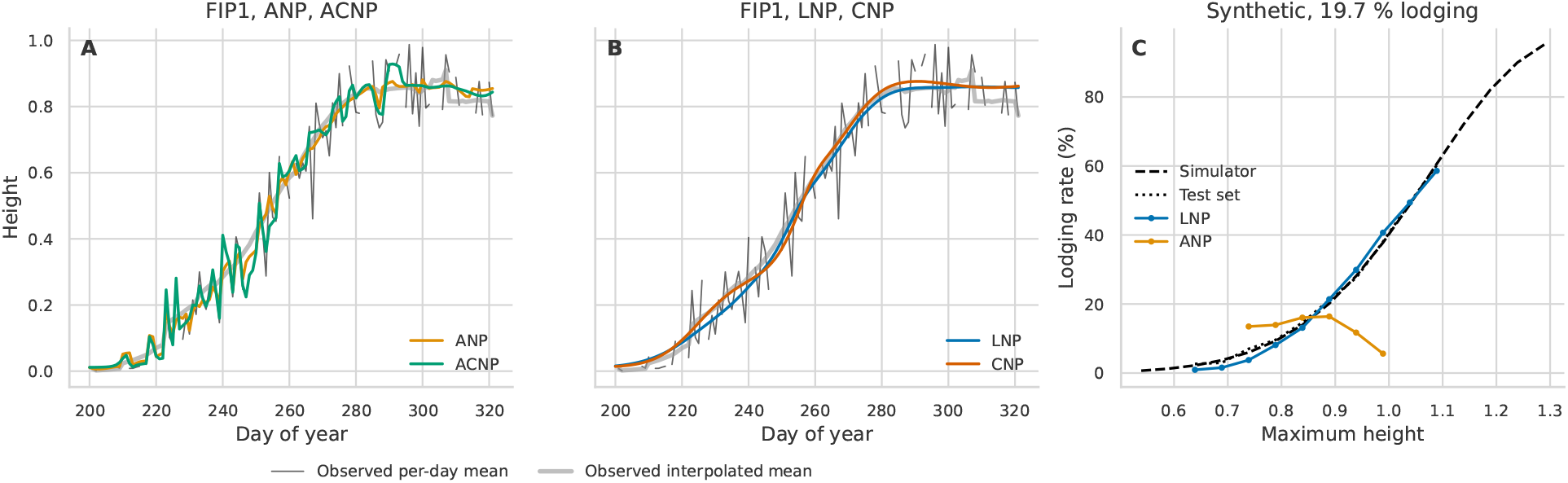
Predictions of the attentive models without context. Panels A and B show the mean prediction without covariates on the seen evaluation split of the Field Phenotyping Platform 1.0 (*FIP1*) dataset, held-out plots of seen genotypes and years, for the attentive (*ANP*) and attentive conditional (*ACNP*) neural processes (A) and the latent (*LNP*) and conditional (*CNP*) neural processes (B), from one training run after pre-training on the simulator. The thin black line is the per-day mean, the mean over the plots measured on each day, and the thick gray line the interpolated mean, the mean over all plots after interpolating each plot between its measurement dates. Days are counted from the first of September. Panel C shows the lodging rate against the predicted maximum height on the Seen split of the synthetic data, for the *LNP* and the *ANP* without covariates. The dashed curve is the analytical lodging probability of the synthetic generator (*Simulator*) and the dotted curve the lodging rate of the evaluation data (*Test set*). Predicted trajectories are counted as lodged when their height falls by at least 20 % and at least 10 cm below their maximum. Heights are in meters.

### List of Abbreviations

*ACNP*: attentive conditional neural process
*ANP*: attentive neural process
*ANP-NF-Prior*: attentive neural process with normalizing flow prior
*AR(1)*: first-order autoregressive
*ANP-NF-Prior-Posterior*: attentive neural process with normalizing flow prior and posterior
*BLUE*: best linear unbiased estimate
*CNP*: conditional neural process
*CRPS*: continuous ranked probability score
*CSig-MMD*: censored signature maximum mean discrepancy
*ELBO*: evidence lower bound
*FIP1*: Field Phenotyping Platform 1.0
*KL*: Kullback–Leibler divergence
*LNP*: latent neural process
*LNP-NF-Prior*: latent neural process with normalizing flow prior
*MLP*: multi-layer perceptron
*LNP-NF-Prior-Posterior*: latent neural process with normalizing flow prior and posterior
*MMD*: maximum mean discrepancy
*MRCD*: minimum regularized covariance determinant
*NF*: normalizing flow
*NP*: neural process
*NSF*: neural spline flow
*PDE*: partial differential equation
*RBF*: radial basis function
*RMS*: root mean square
*RMSE*: root mean squared error
*RMSNorm*: root mean square layer normalization
*SD*: standard deviation
*Sig-MMD*: signature maximum mean discrepancy
*SwiGLU*: Swish-gated linear unit
*VAE*: variational auto-encoder

## Acknowledgments

The authors thank Achim Walter (ETH Zürich, Institute of Agricultural Sciences) for providing the research environment in which this work was carried out.

LR discloses support for the research of this work from the Swiss Data Science Center [grant no. PHENO-MINE C21–04] and from the Swiss National Science Foundation [grant no. IZSEZ0 230796].

This work was supported in part by the Swiss National Science Foundation [grant no. PreDiMix 10002207].

## Data and Code Availability

The code for the synthetic generator, the models, and the evaluation is available at http://hdl.handle.net/20.500.11850/806730. The *FIP1* dataset is published separately [38].

The authors disclose the use of AI tools to assist with code development and manuscript writing. All such contributions were verified by the authors.

